# A nanodomain anchored-scaffolding complex is required for PI4Kα function and localization in plants

**DOI:** 10.1101/2020.12.08.415711

**Authors:** Lise C Noack, Vincent Bayle, Laia Armengot, Rozier F Frédérique, Adiilah Mamode-Cassim, Floris D Stevens, Marie-Cécile Caillaud, Teun Munnik, Sébastein Mongrand, Roman Pleskot, Yvon Jaillais

**Affiliations:** Laboratoire Reproduction et Développement des Plantes, Univ Lyon, ENS de Lyon, Université Claude Bernard Lyon 1, CNRS, INRAE, F-69342, Lyon, France; Laboratoire de Biogenèse Membranaire, UMR5200, Université de Bordeaux, CNRS, 33140 Villenave d’Ornon, France; Research Cluster Green Life Sciences, Section Plant Cell Biology, Swammerdam Institute for Life Sciences, University of Amsterdam, PO Box 94215, Amsterdam, 1090 GE, The Netherlands; Agroécologie, AgroSup Dijon, CNRS, INRA, Univ. Bourgogne Franche-Comté, F-21000 Dijon, France; Institute of Experimental Botany, Academy of Sciences of the Czech Republic, Rozvojová 263, 16502 Prague 6, Czech Republic

## Abstract

Phosphoinositides are low-abundant lipids that participate in the acquisition of membrane identity through their spatiotemporal enrichment in specific compartments. PI4P accumulates at the plant plasma membrane driving its high electrostatic potential, and thereby facilitating interactions with polybasic regions of proteins. PI4Kα1 has been suggested to produce PI4P at the plasma membrane, but how it is recruited to this compartment is unknown. Here, we pin-point the mechanism that tethers PI4Kα1 to the plasma membrane via a nanodomain-anchored scaffolding complex. We identified that PI4Kα1 is part of a complex composed of proteins from the NO-POLLEN-GERMINATION, EFR3-OF-PLANTS, and HYCCIN-CONTAINING families. Comprehensive knock-out and knock-down strategies revealed that subunits of the PI4Kα1 complex are essential for pollen, embryonic and post-embryonic development. We further found that the PI4Kα1 complex is immobilized in plasma membrane nanodomains. Using synthetic mis-targeting strategies, we demonstrate that a combination of lipid anchoring and scaffolding localizes PI4Kα1 to the plasma membrane, which is essential for its function. Together, this work opens new perspectives on the mechanisms and function of plasma membrane nanopatterning by lipid kinases.

## INTRODUCTION

Eukaryotic cells are composed of several membrane-surrounded compartments. Each compartment has a unique physicochemical environment delimited by a membrane with a specific biochemical and biophysical identity (Bigay and Antonny, 2012). The membrane identity includes the nature of the lipids, the curvature, the electrostaticity and the density of lipids at the membrane. The identity of each membrane allows the proper localization of membrane-associated proteins.

Phosphoinositides are rare anionic lipids present in membranes. Five types of phosphoinositides exist in plants - PI3P, PI4P, PI5P, PI(4,5)P2 and PI(3,5)P2 - depending on the number and position of phosphates around the inositol ring (Munnik and Nielsen, 2011; Munnik and Vermeer, 2010). They accumulate differently at the plasma membrane and intracellular compartments and interact with proteins through stereo-specific or electrostatic interactions (Barbosa et al., 2016; Hirano et al., 2017; Lemmon, 2008; Simon et al., 2016). Recent work uncovered that PI4P concentrates according to an inverted gradient by comparison to their yeast and animal counterpart (Hammond et al., 2009, 2014; Levine and Munro, 1998, 2002; Noack and Jaillais, 2017, 2020; Roy and Levine, 2004; Simon et al., 2016). Indeed in yeast, the major PI4P pool is at the Golgi/trans-Golgi Network (TGN) compartments while a minor pool is present at the plasma membrane (Roy and Levine, 2004). The plasma membrane pool of PI4P is produced by the PI4-Kinases (PI4K) Stt4p, while Pik1p produces the PI4P pool at the TGN (Audhya and Emr, 2002; Audhya et al., 2000; Balla et al., 2005; Nakatsu et al., 2012; Roy and Levine, 2004). These two PI4P pools are essential for yeast survival and at least partially independent (Roy and Levine, 2004). In animal, three PI4K isoforms, PI4KIII β/PI4KIIα/PI4KIIβ, are responsible for synthetizing PI4P at the Golgi/TGN and in endosomes (Balla et al., 2002; Wang et al., 2003; Wei et al., 2002). Similar to the Δstt4 in yeast, PI4KIIIα loss-of-function mutant is lethal in mammals (Nakatsu et al., 2012). In PI4KIIIα conditional mutants, the pool of PI4P disappears from the plasma membrane while the TGN structures seem to remain untouched suggesting that the two pools could be independent (Nakatsu et al., 2012).

In plants, PI4P massively accumulates at the plasma membrane and is less abundant at the TGN (Simon et al., 2014, 2016; Vermeer et al., 2009). This PI4P accumulation at the cell surface drives the plasma membrane electrostatic field, which in turn recruits a host of signalling proteins to this compartment (Barbosa et al., 2016; Platre et al., 2018; Simon et al., 2016). Moreover, the plant TGN is the site of vesicular secretion but is also involved in endocytic sorting and recycling, which might imply regulatory mechanisms of lipid exchanges or maintenance of membrane identity between the plasma membrane and the TGN (Noack and Jaillais, 2017).

The Arabidopsis genome codes four PI4-kinases: PI4Kα1, PI4Kα2, PI4Kβ1 and PI4Kβ2 (Szumlanski and Nielsen, 2010). Due to the absence of Expressed Sequence Tags of PI4Kα2, it is considered as a pseudogene (Mueller-Roeber and Pical, 2002). pi4kβ1pi4kβ2 double mutant displays mild growth defects including tip growth phenotype with bulged root hairs and cell plate defects, which suggest a defective secretory pathway (Antignani et al., 2015; Delage et al., 2012; Kang et al., 2011; Lin et al., 2019; Preuss et al., 2006; Šašek et al., 2014). In addition, pi4kβ1pi4kβ2 presents fewer and misshaped secretory vesicles at the TGN (Kang et al., 2011). PI4Kβ1 and PI4Kβ2 have first been described to be localized to the Trans-Golgi Network/Early Endosomes (TGN/EE) in root hairs (Preuss et al., 2006). This localization, as well as its accumulation at the cell plate, has later been validated by electron tomography and confocal microscopy in root meristematic cells (Kang et al., 2011; Lin et al., 2019). The targeting mechanism of PI4Kβ1 at the TGN involves RabA4b, a small GTPase (Preuss et al., 2006). In addition, PI4Kβ1 recognizes, and interacts with, the curved electronegative membrane of the TGN/EE via an amphipatic lipid packing sensor (ALPS) motif preceded by cationic amino acids (Platre et al., 2018).

In contrast, PI4Kα1 localizes at the plasma membrane (Okazaki et al., 2015) and its catalytic activity was confirmed in vitro (Stevenson-Paulik et al., 2003). Thus, it is a prime candidate for producing PI4P at the plasma membrane. However, PI4Kα1 is a soluble protein with no protein-lipid interaction domains or anchoring mechanism characterized in planta. How PI4Kα1 is recruited and participates to the architecture of the plasma membrane are open questions.

Here, we uncovered that PI4Kα1 belongs to a 4-subunit complex composed of proteins from the NPG (NO POLLEN GERMINATION), HYC (HYCCIN-CONTAINING) and EFOP (EFR3 OF PLANTS) protein families. Using fluorescent protein tagging, immunolocalization and subcellular fractionation, we confirmed the presence of the PI4Kα1 complex at the plasma membrane. Furthermore, we show that pi4kα 1 loss-of-function leads to full male sterility. Mutant pollen grains collapse and display abnormal cell walls. Knockout of any subunits of the PI4Kα1 complex mimics pi4kα1 pollen lethality. Moreover, we established that the four subunits of the complex are essential for PI4Kα1 function. By using mutant variants and chimeric constructs, we showed that the function of this complex is to target PI4Kα1 to the plasma membrane via lipid anchors. Finally, we observed that this heterotetrameric complex is not homogenously present on the plasma membrane but enriched in nanodomains. Although all the subunits of the complex are peripheral proteins and lack a transmembrane domain, they show very little lateral mobility at the plasma membrane. These results suggest that PI4Kα1 is not localized homogeneously at the plasma membrane but rather accumulates in distinct hotspots at the inner leaflet of the plasma membrane. Consequently, the targeting of this lipid kinase by a multiprotein scaffold might allow its precise spatiotemporal recruitment in order to maintain the proper electrostatic landscape of plant cell membranes.

## RESULTS

### PI4Kα1 is a soluble protein with a potential lipid-binding domain

To determine how PI4Kα1 is recruited to the plasma membrane, we first analysed its protein sequence in silico. Using TMHMM Server v. 2.0, no transmembrane helices could be predicted in PI4Kα1 protein sequence suggesting that PI4K α1 is a soluble cytosolic protein. We then looked for lipid binding domains. Indeed, a Plecthrin Homology (PH) domain was previously reported in its C-terminal, upstream from the catalytic domain (Stevenson et al., 1998; Stevenson-Paulik et al., 2003; Xue et al., 1999). This PH domain was first thought to localize PI4Kα1 at the plasma membrane through interaction with anionic phospholipids but its role is now discussed (De Jong and Munnik, 2021). Fat blot experiments showed affinity of the putative PH domain for PI4P and in a lesser extent for PI(4,5)P2 (Stevenson et al., 1998; Stevenson-Paulik et al., 2003). However, no experiment in planta validates this result and using the Simple Modular Architecture Research Tool (SMART) software, we were not able to retrieve the PH domain. Because of the lack of predicted domain (except for the kinase domain), we decided to consider other targeting mechanisms involving possible protein partners. Indeed, protein targeting to a membrane can be multifactorial and may require coincidence binding of lipids and protein partners.

### PI4Kα1 interacts with NO POLLEN GERMINATION proteins

To investigate this last hypothesis, we screened for PI4Kα 1-protein partners. We performed a yeast-two-hybrid screen with the large N-terminal part of PI4Kα1 (1-1468 aa) (Figure 1A). We recovered 267 in frame clones, which corresponded to 48 different proteins. Among them, the screen revealed interactions between PI4Kα1 and the three members of a protein family called NO POLLEN GERMINATION (NPG): NPG1 (At2g43040), NPG-Related 1 (NPGR1 – At1g27460) and NPGR2 (At4g28600) (Golovkin and Reddy, 2003). In the screen, we retrieved 39 clones (7 independent clones) for NPG1, 32 clones (6 independent clones) for NPGR1 and 2 clones (1 independent clone) for NPGR2. The clones from the NPG family corresponded to about 30% of the total clones obtained from the screen, suggesting that they were over-represented.

**Figure 1.**
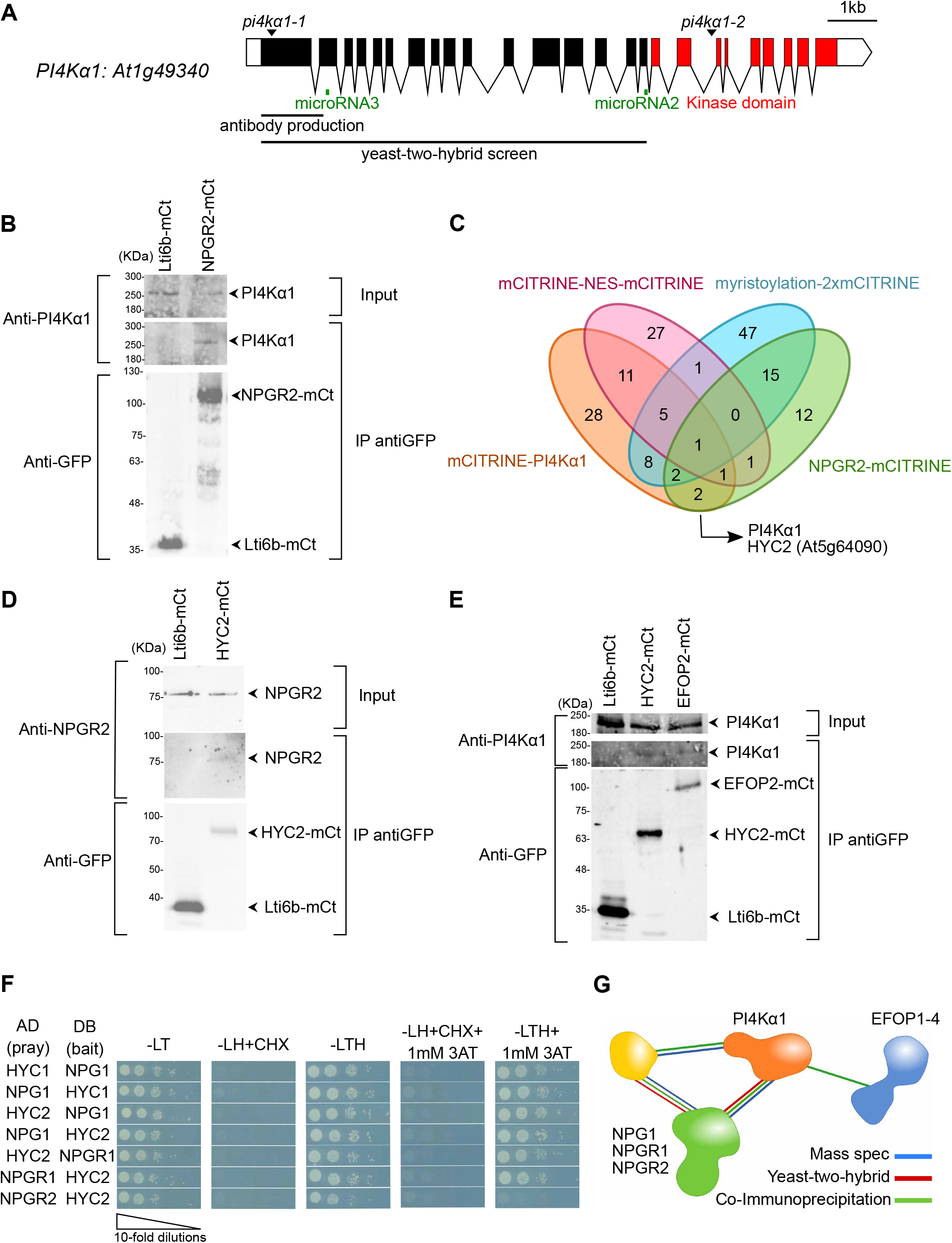
PI4Kα1 interacts with proteins from the NPG, HYC and EFOP families. (A) Schematic representation of PI4Kα1 (At1g49340) gene. Boxes and lines represent exons and introns, respectively. The kinase domain is shown in red. The T-DNA positions of the pi4kα1-1 and pi4kα1-2 alleles are indicated. Parts of the protein used for the yeast-two-hybrid screen and the antibody production are also shown. The regions targeted by microRNA#2 and #3 are indicated in green. (B) Co-immunoprecipitation of PI4Kα1 with NPGR2. Arabidopsis transgenic plants overexpressing NPGR2-mCITRINE (NPGR2-mCt) or Lti6b-mCITRINE (Lti6b-mCt) were used for immunoprecipitation using anti-GFP beads. Immunoblots used anti-PI4Kα1 (upper panel) and anti-GFP (lower panel). (C) Venn diagram of proteins identified by mass spectrometry from immunoprecipitation of mCITRINE-PI4Kα1, NPGR2-mCITRINE, mCITRINE-NES-mCITRINE and myristoylation-2x-mCITRINE. (D) Co-immunoprecipitation of NPGR2 with HYC2. Arabidopsis transgenic plants overexpressing HYC2-mCITRINE (HYC2-mCt) or Lti6b-mCITRINE were used for immunoprecipitation using anti-GFP beads. Immunoblots used anti-GFP (lower panel) and anti-NPGR2 (upper panel). (E) Co-immunoprecipitation of PI4Kα1 with EFOP2 and HYC2. Arabidopsis transgenic plants overexpressing EFOP2-mCITRINE (EFOP2-mCt), HYC2-mCITRINE or Lti6b-mCITRINE were used for immunoprecipitation using anti-GFP beads. Immunoblots used anti-PI4Kα1 (upper panel) or anti-GFP (lower panel). (F) Yeast-two hybrid assay of HYC1 with NPG1, HYC2 with NPG1 or NPGR1 and NPGR2. Indicated combinations of interactions between HYCCIN-CONTAINING and NPG proteins were assessed by growth on plates with yeast growth media lacking Leu, Trp, and His (−LTH). Yeast growth on plates lacking Leu and Trp (−LT) shows the presence of the bait and prey vectors. Absence of growth when cycloheximide was added (+CHX) shows the absence of auto-activation of the DB vectors. Addition of 3-Amino-1,2,4-Triazol (+3AT) shows the strength of the interaction. (G) Summary of experiments showing interactions between PI4Kα1, NPG, HYC and EFOP2 proteins.

NPG proteins contain tetratricopeptide repeats (TPR) motifs that are protein-protein interaction motifs. In the yeast-two-hybrid screen, the selected interaction domain identified for NPG1, NPGR1 and NPGR2 correspond to the C-terminal part of the proteins (aa 444-704 for NPG1; 501-694 for NPGR1 and 414-739 for NPGR2). This is also the part of the sequence that contains the highest density of predicted TPR motifs suggesting that the interaction between PI4Kα1 and NPGs could be mediated by the C-terminal TPR motifs.

Because all three members of the NPG family interacted with PI4Kα1 in yeast-two hybrids and given the high degree of identity and similar architecture of the three proteins, we decided to focus on one member of the family to confirm the interaction in planta. We guided this choice based on the RNAseq expression data compiled on eFP browser (https://bar.utoronto.ca/efp/cgi-bin/efpWeb.cgi). We chose NPGR2, as it is the family member with the highest and more widespread expression. Indeed, NPG1 was predicted to be specifically expressed in the pollen, while NPGR1 expression matched that of NPGR2 but was predicted to be much weaker.

To confirm the interaction between NPGR2 and PI4Kα1, we produced stable transgenic lines expressing UBQ10:NPGR2-mCITRINE. We raised antibodies against the native PI4Kα1 (residues 1 to 344 of PI4Kα1). In western blot the antibody recognized PI4Kα1 around the expected size (225kDa) and the tagged version of PI4Kα1 with mCITRINE and 2xmCHERRY slightly higher (Supplemental Figure 1A; Table S1). We immunoprecipitated NPGR2-mCITRINE or the plasma membrane protein Lti6b-mCITRINE as control using anti-GFP antibodies and probed whether they could co-immunoprecipitate PI4Kα1, using our native antibody. We efficiently immunoprecipitated NPGR2-mCITRINE or Lti6b-mCITRINE, but PI4Kα1 was only co-immunoprecipitated with NPGR2-mCITRINE (Figure 1B). Together, these experiments suggest that PI4Kα1 can interact in yeast with the C-terminus of all the three members of the NPG family and is at least found in complex with NPGR2 in planta.

### NPG proteins interact with HYCCIN-CONTAINING proteins

Next, we asked whether the PI4Kα1 and NPG proteins could interact with additional protein partners. To this end, we used the lines expressing mCITRINE-PI4Kα1 and NPGR2-mCITRINE to perform immunoprecipitation (IP) followed by mass spectrometry analyses. Two lines expressing membrane-associated (myristoylation-2xmCITRINE) and nuclear-excluded (mCITRINE-NES-mCITRINE) proteins were also used as generic controls for plasma membrane and cytosolic proteins, respectively. In the NPGR2 IP, we found PI4Kα1, further confirming that these two proteins are present in the same complex in plants. Only one common protein was found in both NPGR2 and PI4Kα1 IPs but excluded from the two controls (Figure 1C). This protein was coded by the At5g64090 locus and contains a HYCCIN domain. Arabidopsis genome codes for only two proteins with a HYCCIN domain subsequently called HYCCIN-CONTAINING1 (HYC1, At5g21050) and HYCCIN-CONTAINING2 (HYC2, At5g64090).

HYC2 is broadly expressed, while HYC1 expression is restricted to pollen according to the eFP browser data set (https://bar.utoronto.ca/efp/cgi-bin/efpWeb.cgi). Hence, we chose to confirm whether HYC2 was interacting with PI4Kα 1 and NPGR2 in sporophytic tissues. To this end, we raised a UBQ10::HYC2-mCITRINE expressing lines and successfully isolated antibodies raised against NPGR2 (residues 1 to 273 of NPGR2). The expected size of NPGR2 is 82kDa. The antibody recognized a band at ca. 80kDa that is not present in npgr2-1 or npgr2-3 knock out mutants or npgr1npgr2-1 double mutant (Supplemental Figure 1B; Table S1). Moreover, the antibody recognized NPGR2-mCITRINE around 110kDa, but did not recognize NPGR1-mCITRINE indicating that the antibody specifically recognized NPGR2 (Supplemental Figure 1B). We found that NPGR2 co-immunoprecipitated with HYC2-mCITRINE but not Lti6b-mCITRINE (Figure 1D). Similarly, PI4Kα1 also co-immunoprecipitated with HYC2-mCITRINE but not with Lti6b-mCITRINE (Figure 1E). Next, we used yeast-two hybrid to check whether the two HYC family members may directly interact with PI4Kα1/NPGs (Figure 1F). We found that the two isoforms that are pollen specific, HYC1 and NPG1 interacted in yeast. Likewise, HYC2 interacted with NPG1, NPGR1 and NPGR2 in yeast. Together, our results indicate that HYC family members directly interact with NPG proteins. These data suggest that HYC2 is present in complex with PI4Kα1 and with NPGR2 in the Arabidopsis sporophyte, while HYC1/NPG1/PI4Kα1 could form a similar complex in the male gametophyte.

### A structural modelling approach suggests that PI4Kα1, NPG and HYC form a heterotrimeric complex in plants

HYCCIN domain-containing proteins are also found in Metazoa. In human cells, one of the members of the HYCCIN family, FAM126, is part of a complex containing PI4KIIIα (Baskin et al., 2016; Dornan et al., 2018). In this complex, TETRATRICOPEPTIDE REPEAT PROTEIN 7 (TTC7) bridges together PI4KIIIα and FAM126 (Baskin et al., 2016; Dornan et al., 2018; Lees et al., 2017a; Wu et al., 2014a). While no HYCCIN containing proteins are found in yeast, the plasma membrane PI4-Kinase, Stt4p also forms a complex containing Ypp1p, which like NPG and TTC7 proteins contain TPR motives (Baird et al., 2008; Nakatsu et al., 2012; Wu et al., 2014a). Based on these information, we hypothesized that NPGs and HYCs in plants could be functional homologs of TTC7/Ypp1p and FAM126, respectively. The structure of the human trimeric complex formed by PI4KIIIα-TTC7-FAM126 has been determined by cryo-electron microscopy (Baskin et al., 2016; Dornan et al., 2018; Lees et al., 2017a). We thus decided to use a templated-based modeling approach together with protein-protein docking to analyze the conservation of the overall structure of individual subunits and of their respective binding interface in particular. Using the structure of the human PI4KIII α, TTC7 and FAM126 as templates, we modelled the structure of PI4Kα1 (aa 859-2028), NPG1 (aa 43-704) and HYC1 (aa 51-331) from Arabidopsis. Using this approach, we obtained a highly confident structural model for each Arabidopsis subunit (Supplemental Figure 2 A-C). We next predicted binding interfaces of the heterodimer composed of PI4Kα1-NPG1 and NPG1-HYC1 by a hybrid docking method using the HDOCK algorithm (Figure 2 A-B). We found that these interfaces are formed by highly conserved amino acids, suggesting that they are structurally important and supporting their potential key roles in the formation of supporting their potential key roles in the formation of the PI4Kα1-NPG1 and NPG1-HYC1 dimers. Of note, a majority of amino acid residues forming the PI4Kα1-binding interface of NPG1 (aa 585-655) corresponded to the selected interaction domain identified experimentaly in the yeast two hybrid screen (aa 444-704) (Figure 2A). Finally, we compared the overall experimental structure of the human PI4KIII α complex with the model of the Arabidopsis PI4Kα1 complex obtained using the template-based modelling and protein-protein docking approach (Figure 2C). The structural comparison between the two structures revealed a highly similar arrangement of both complexes, again supporting the notion that Arabidopsis PI4Kα1-NPG1-HYC1 are structural homologs of human PI4KIII α-TTC7-FAM126.

**Figure 2.**
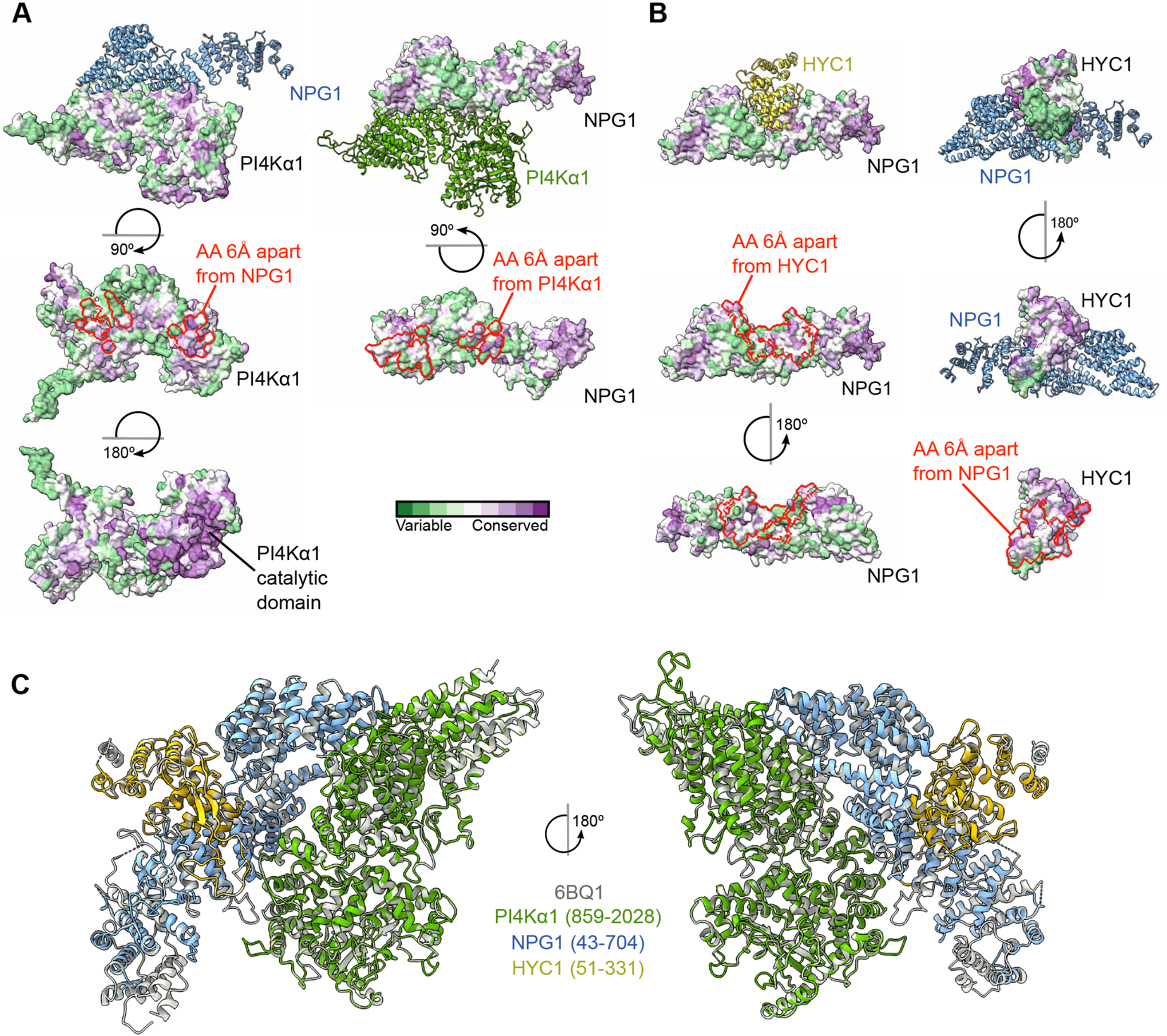
Template-based modelling and protein-protein docking suggest that the plant PI4Kα1 forms a stable heterotrimeric complex. (A) Heterodimer formed by PI4Kα1 (green) and NPG1 (blue) as calculated by a hybrid docking approach using the HDOCK algorithm. Analysis of conserved amino acid residues was performed utilizing the Consurf server and mapped on the solvent-excluded surface of each protein. Calculated protein-protein interfaces indicated by red lines are formed by highly conserved amino acid residues. (B) Heterodimer formed by NPG1 (blue) and HYC1 (yellow) as calculated by a hybrid docking approach using the HDOCK algorithm. Analysis of conserved amino acid residues was performed utilizing the Consurf server and mapped on the solvent-excluded surface of each protein. Calculated protein-protein interfaces indicated by red lines are formed by highly conserved amino acid residues. (C) Comparison of the experimental structure of the human PI4KIIIα complex (grey, PDB code 6BQ1) with the heterotrimeric Arabidopsis PI4Kα1 complex obtained using template-based modelling and protein-protein docking.

The modelling approach together with interaction data suggest that PI4Kα1, HYC and NPG proteins form a stable heterotrimeric complex in planta, which is likely equivalent to its metazoan counterpart.

### EFOPs proteins are part of the PI4Kα1-NPG-HYC complex

In human cells and yeast, the PI4-Kinase complex contains an additional subunit called EFR3/Efr3p (Baird et al., 2008; Baskin et al., 2016; Dornan et al., 2018; Lees et al., 2017a; Nakatsu et al., 2012; Wu et al., 2014b). We thus searched whether EFR3 homologs could exist in Arabidopsis using blast and sequence alignments. We found four potential candidates that we named EFR3 OF PLANT (EFOPs): EFOP1 (At5g21080), EFOP2 (At2g41830), EFOP3 (At1g05960) and EFOP4 (At5g26850).

Yeast Efr3p is a rod-shaped protein made of ARMADILLO-(ARM) and HEAT-like repeats. ARM and HEAT repeats are difficult to be distinguished bioinformatically, but all four EFOP proteins belong to the ARM-repeat super-family, which includes both ARM and HEAT repeat containing proteins (Wu et al., 2014a). In Marchantia polymorpha, mutants for MpPI4Kα1 (the homolog of PI4Kα1) and MpSRI4, the homolog of EFOP2, display short rhizoids, suggesting that they could act in the same pathway and/or protein complex (Honkanen et al., 2016). In addition, based on RNAseq data, EFOP2 have a rather large pattern of expression. Thus, we decided to concentrate on EFOP2 to test whether it is indeed present in the sporophytic PI4Kα 1/NPGR2/HYC2 complex. To this end, we raised UBQ10prom:EFOP2-mCITRINE transgenic lines and immunoprecipitated EFOP2-mCITRINE and Lti6b-mCITRINE using an anti-GFP antibody. We found that PI4Kα1 co-immunoprecipitated with EFOP2-mCITRINE while it did not with Lti6b-mCITRINE (Figure 1E), suggesting that EFOP2 may belong to the PI4Kα1/NPGR2/HYC2 complex. The summary of these interactions suggests that PI4Kα1 is part of an heterotetrameric complex in which NPG proteins may act as a scaffold that bridges EFOP, HYC and PI4Kα1 proteins together (Figure 1G).

### *pi4kα1* mutants are pollen lethal

Next, we took a genetic approach to confirm whether NPG, HYC and EFOP family members indeed may function together with PI4Kα1 in planta. To this end, we isolated single mutants for all the genes encoding for a subunit of the PI4Kα1 complex (Table S2). We started our analysis with PI4Kα1 because it is the catalytic subunit and it is present as a single-copy gene in the Arabidopsis genome for which we isolated two T-DNA insertion alleles. The first allele (*pi4kα1-1*; GK_502D11) had an insertion in the first exon, while the second insertion (*pi4kα1-2*; FLAG_275H12) was in the 20th intron (Figure 1A). T-DNAs of the first and second allele possess a sulfadiazine and glufosinate resistant gene, respectively. We failed to obtain homozygous mutant plants for both alleles. The segregations obtained after self-fertilization of heterozygous plants were 38% of sulfadiazine resistant seedlings for *pi4kα1-1* and 9% of glufosinate resistant seedlings for *pi4kα1-2* (Table 1). Because there were less than 50% of resistant plants in the progeny of self-fertilized *pi4kα1* alleles, these segregations indicated likely gametophyte lethality, which might explain the absence of homozygous *pi4kα1* mutant. To address whether *pi4kα1* could be female and/or male gametophytic lethal, we performed reciprocal crosses, using *pi4kα1+/−* and wild-type as either male or female. For both alleles, we recovered 0% resistant plants when *pi4kα1+/−* was used as the male indicating no transmission of the mutation via the pollen and thus complete male sterility (Table 2). When *pi4k α1+/−* was used as female, we obtained 39% and 9% of resistant plants for each allele (Table 2). This result shows that the *pi4kα1* mutation did not fully impair the transmission through the female gametophyte but led to a partial distortion of the segregation.

**Table 1:** Segregation analyses of the indicated self-fertilized heterozygous mutants. n represent the number of seedlings analysed.

| Segregation |  |  |  |  |  |  |
| --- | --- | --- | --- | --- | --- | --- |
| Gene | Allele | Cross |  | % of resistant plants | % expected | n |
|  |  | Female | Male |  |  |  |
| PI4Kα1 | *pi4ka1-1* | *pi4ka1-1 +/-* | *pi4ka1-1 +/-* | 38,2 | 75 | 418 |
|  | *pi4ka1-2* | *pi4ka1-2 +/-* | *pi4ka1-2 +/-* | 9,4 | 75 | 359 |
| NPG1 | *npg1-1* | *npg1-1 +/-* | *npg1-1 +/-* | 50,3 | 75 | 296 |
|  | *npg1-2* | *npg1-2 +/-* | *npg1-2 +/-* | 32 | 75 | 504 |
| HYC1 | *hyc1* | *hyc1 +/-* | *hyc1 +/-* | 38,2 | 75 | 128 |
| HYC2 | *hyc2-2* | *hyc2-2 +/-* | *hyc2-2 +/-* | 59,9 | 75 | 133 |
|  | *hyc2-3* | NO RESISTANCE |  |  |  |  |

**Table 2:** Segregation analyses of reciprocal crosses between wild-type and the indicated mutants. n represent the number of seedlings analysed.

| Reciprocal Crosses |  |  |  |  |  |  |
| --- | --- | --- | --- | --- | --- | --- |
| Gene | Allele | Cross |  | % of resistant plants | % expected | n |
|  |  | Female | Male |  |  |  |
| PI4Kα1 | *pi4ka1-1* | *pi4ka1-1 +/-* | Col0 | 39 | 50 | 424 |
|  |  | Col0 | *pi4ka1-1 +/-* | 0 | 50 | 268 |
|  | *pi4ka1-2* | *pi4ka1-2 +/-* | Col0 | 9,3 | 50 | 106 |
|  |  | Col0 | *pi4ka1-2 +/-* | 0 | 50 | 293 |
| NPG1 | *npg1-1* | *npg1-1 +/-* | Col0 | 40,6 | 50 | 214 |
|  |  | Col0 | *npg1-1 +/-* | 0 | 50 | 163 |
|  | *npg1-2* | *npg1-2 +/-* | Col0 | 30,8 | 50 | 26 |
|  |  | Col0 | *npg1-2 +/-* | 0 | 50 | 111 |
| HYC1 | *hyc1* | *hyc1 +/-* | Col0 | 47 | 50 | 185 |
|  |  | Col0 | *hyc1 +/-* | 0 | 50 | 58 |
| HYC2 | *hyc2-2* | *hyc2-2 +/-* | Col0 | 29,6 | 50 | 54 |
|  |  | Col0 | *hyc2-2 +/-* | 29,8 | 50 | 47 |
|  | *hyc2-3* | NO RESISTANCE |  |  |  |  |
| ELP3 ELP4 | *elp3-1 elp4-2* | *elp3-1 +/- elp4-2/-* | Col0 | 55,7 | 50 | 97 |
|  |  | Col0 | *elp3-1 +/- elp4-2/-* | 2,2 | 50 | 45 |
|  | *elp3-2 elp4-2* | *elp3-2 +/- elp4-2/-* | Col0 | 32,4 | 50 | 68 |
|  |  | Col0 | *elp3-2 +/- elp4-2/-* | 0 | 50 | 31 |
|  | *elp3-1 elp4-4* | *elp3-1 +/- elp4-4/-* | Col0 | 48,1 | 50 | 27 |
|  |  | Col0 | *elp3-1 +/- elp4-4/-* | ND | 50 | ND |
|  | *elp3-2 elp4-4* | *elp3-2 +/- elp4-4/-* | Col0 | 50 | 50 | 12 |
|  |  | Col0 | *elp3-2 +/- elp4-4/-* | 0 | 50 | 6 |

Next, we observed *pi4kα1* pollen grains using scanning electron microscopy (SEM) to test whether they showed morphological defects (Figure 3A). For both alleles, half of the pollen grains were shrivelled and likely not able to germinate explaining the pollen lethality (Figure 3A-B). However, using Alexander staining, we observed that the pi4kα1-1 pollens were still alive (Supplemental Figure 3B). DAPI staining also revealed the presence of the vegetative nucleus and the two sperm cell nuclei indicating that meiosis likely occurred normally (Supplemental Figure 3C). Further analysis by transmission electron microscopy showed that the *pi4kα1-1* pollen grains displayed an abnormally thick intine layer (Figure 3C).

**Figure 3.**
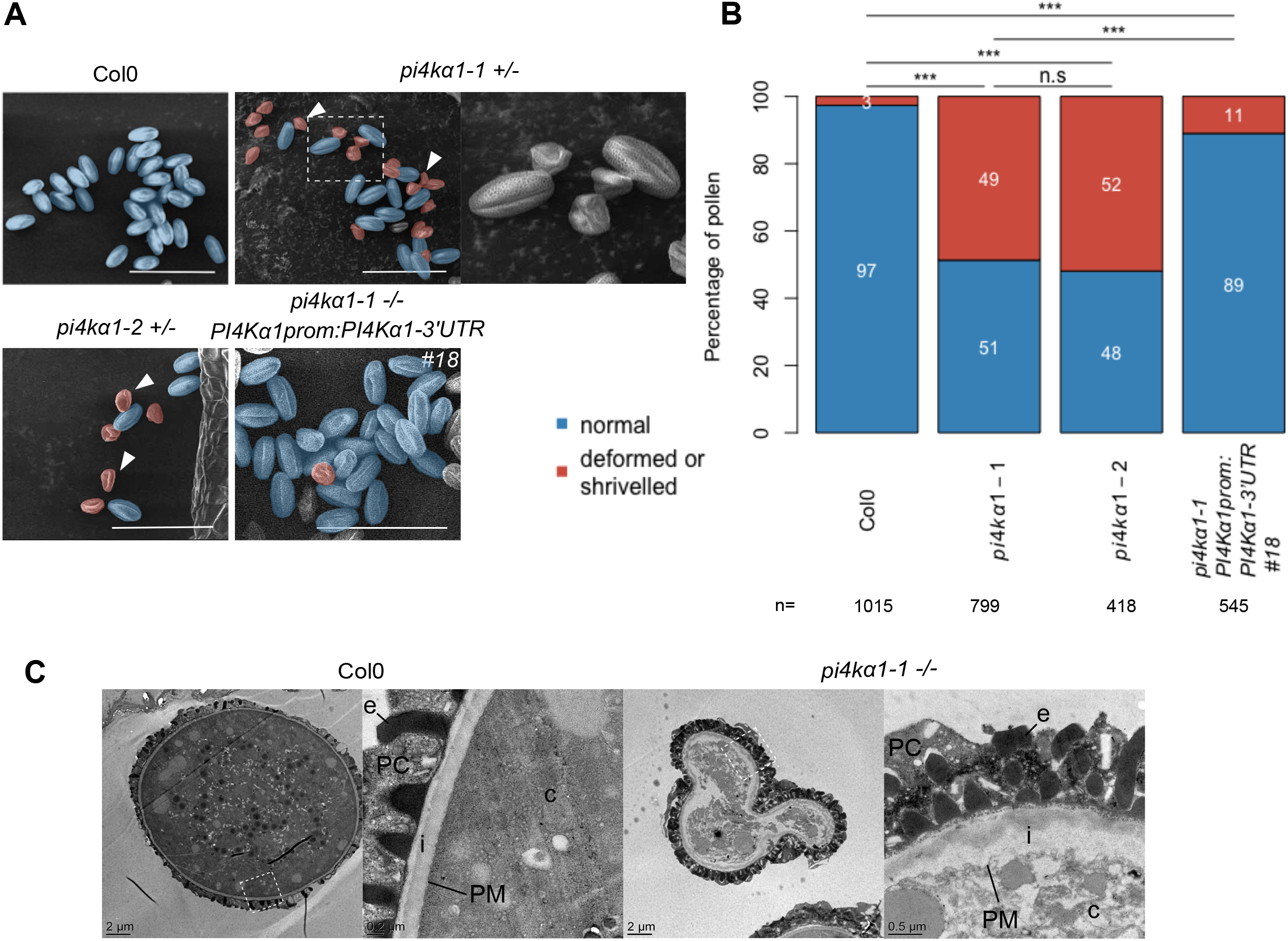
PI4Kα1 loss-of-function leads to pollen lethality. (A) Scanning electron microscope micrograph of pollen grains from Col-0, self-fertilized *pi4kα1-1* heterozygous plants, self-fertilized *pi4kα1-2* heterozygous plants, and self-fertilized *pi4kα1-1* homozygous plants expressing *PI4Kα1prom:PI4Kα1-3’UTR* (insertion n°18). Shrivelled pollen grains are colored in red and normal pollen grains are colored in blue. Close-up is shown for pi4kα1-1 pollen on the right. Scale bar: 50 μm (B) Quantification of the percentage of normal (blue) versus deformed/shrivelled (red) pollen grains from Col-0, self-fertilized pi4kα1-1 heterozygous plants, self-fertilized *pi4kα1-2* heterozygous plants, and self-fertilized *pi4kα1-1* homozygous plants expressing *PI4Kα1prom:PI4Kα1-3’UTR* (insertion n°18). n indicates the number of pollens counted for each genotype. Statistics used chi-square test. n.s, non-significant; ***, p<0.001. (C) Observation of Col-0 and *pi4kα1-1* shrivelled pollen grains by transmission electron microscopy. Right panels show close–up of the region indicated on the left panel. c, cytosol; PM, plasma membrane; i, intine; e, exine; PC, pollen coat.

The reintroduction of a wild-type copy of PI4Kα1 under the control of its own promoter in *pi4kα1-1* background fully complemented the *pi4kα1-1* lethality as shown by the possibility to obtain homozygous mutant plants (three independent complemented lines, see Supplemental Figure 3A). In addition, self-fertilized *pi4kα1-1−/−; PI4Kα 1prom:PI4Kα1* plants showed a low number of shrivelled pollen grains, comparable to control plants, indicating that a wild-type copy of *PI4Kα1* is required for transmission through the male gametophyte and normal pollen morphology (Figure 3A-B). Together, these results show that *PI4Kα 1* is an essential gene to regulate pollen development, and in particular its cell wall deposition.

### Disturbing subunits of the PI4Kα1 complex mimics pi4k α1 pollen phenotypes

Next, we isolated single mutants for all the genes encoding for *NPG*, *HYC* and *EFOP* subunits in order to ask whether they would recapitulate *pi4kα1* loss-of-function phenotype (Table S2).

The *npg1* mutant was previously published as not being able to germinate giving its name NO POLLEN GERMINA-GERMINATION to the family (Golovkin and Reddy, 2003). We reproduced this result by characterizing two new T-DNA mutant alleles of *NPG1*. The self-progeny of *npg1-1+/−* and *npg1-2+/−* had segregation rate of 50,3% and 32% resistant seedlings, respectively, indicating gamete lethality (Table 1). Reciprocal crosses confirmed their male sterility phenotype, with 0% of transmission of the mutation through the pollen, while the female gametophyte might be affected only for the second allele with a weak distortion of the segregation rate (Table 2). However, the observation of *npg1-1* and *npg1-2* pollen grains by SEM did not show any morphological defect, unlike *pi4kα1* pollens (Figure 4A-B; Supplemental Figure 4A-B). The reintroduction of NPG1 fused with mCITRINE under the control of its own promoter complemented the male sterility in *npg1-2* background leading to *npg1-2* homozygous plants (Supplemental Figure 4C). Similarly, the expression of NPGR2 fused to the mCITRINE under the control of the NPG1 promoter also complemented *npg1-2* male sterility. These experiments indicate that NPGR2 can substitute for NPG1 function in pollen and that both NPG1-mCITRINE and NPGR2-mCITRINE fusion are fully functional (Supplemental Figure 4C).

**Figure 4.**
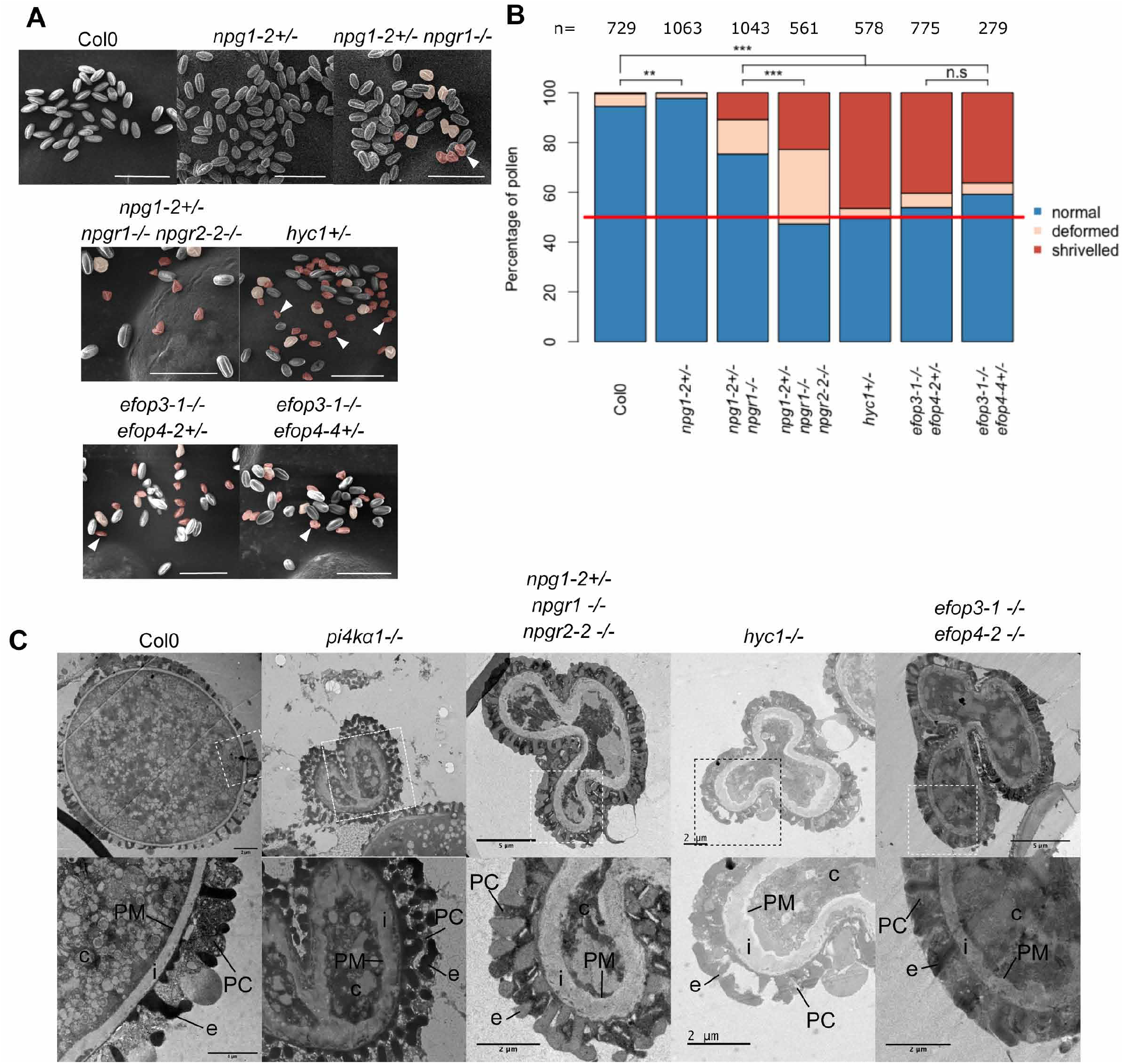
NPG, HYC and EFOP mutations recapitulate the pi4kα1 gametophytic phenotype including no male transmission and shrivelled pollen grain with a thick cell wall. (A) Pollen grains observed by scanning electron microscopy of self-fertilized Col-0, *npg1-2 +/−, npg1-2+/− npgr1−/−, npg1-2+/− npgr1−/− npgr2-2−/−, hyc1+/−, efop3-1−/−efop4-2+/−* and *efop3-1−/−efop4-4+/−* plants. Deformed pollens and shrivelled pollens are colored in red and orange, respectively. Scale bar: 50 *μ*m (B) Percentage of normal (blue), deformed (orange) and shrivelled (red) pollen grains from self-fertilized Col-0, *npg1-2 +/−, npg1-2+/− npgr1−/−, npg1-2+/− npgr1−/− npgr2-2−/−, hyc1+/−, efop3-1−/−efop4-2+/−* and *efop3-1−/−efop4-2+/−* plants. n indicates number of pollens counted for each genotype. Statistics used chi-square test. n.s, non-significant; ***, p<0.001. (C) Observation of pollen grains from self-fertilized Col-0, *pi4kα1-1, npg1-2+/− npgr1−/− npgr2-2−/−, hyc1* and *efop3-1−/−efop4-2−/−* by transmission electron microscopy. Lower panel shows close–up of region indicated on upper panel. c, cytosol; PM, plasma membrane; i, intine; e, exine; PC, pollen coat.

Because, NPGR2 can substitute for NPG1 in pollen, we speculated that a certain degree of functional redundancy between NPG1, NPGR1 and NPGR2 or compensatory effects during pollen development could lead to the weaker phenotype of *npg1* pollen compared to *pi4kα 1* pollen and thus explain why npg1 pollen did not present morphological defect by SEM. To test this hypothesis, we generated higher order mutant combination within the NPG family (Table S2). The *npg1-2+/− npgr1−/−* mutant combination presented about 10% of *pi4kα1*-like shrivelled pollen grains while npgr1npgr2-1 and npgr1npgr2-2 double homozygous mutants displayed about 25% of deformed (but not shrivelled) pollen grains (Figure 4A-B; Supplemental Figure 4A-B). Finally, the *npg1-2+/−npgr1−/−npgr2-1−/−* and *npg1-2+/−npgr1−/−npgr2-2−/−* displayed about 35 and 50% of deformed and shrivelled pollen grains, respectively. Furthermore, images in transmission electron microscopy showed similar thickening of the cell wall for *pi4kα1* and *npg1-2npgr1npgr2-2* shrivelled pollen grains (Figure 4C). These data indicate that *npg* single/multiple mutants partially or fully mimic *pi4kα1* pollen phenotype depending on the allelic combinations.

Next, we addressed the loss-of-function phenotypes of the HYCCIN-CONTAINING family members. The self-progeny of a *hyc1+/−* single mutant presented a segregation rate around 50% indicating a gametophytic lethality (Table 1). As HYC1 expression is restricted to pollen, we were expecting that the segregation bias was caused by defects of the male gametophyte. As anticipated, reciprocal crosses showed complete male sterility while the T-DNA transmission through the female gametophyte was not affected (Table 2). Observation of hyc1+/− pollen grains by SEM revealed that half of the pollen grains were shrivelled (Figure 4A and B). In addition, transmission electron microscopy also showed a thickening of the cell wall of the *hyc1* mutant pollen grains, which was similar to the phenotype observed for *pi4kα1-1* (Figure 4C). Finally, the male sterility, as well as the pollen morphological defects, were complemented by the reintroduction of *HYC1prom:HYC1-mCITRINE* (Supplemental Figure 4A-B and D). All together, these data show that hyc1 knockout phenotype fully mimic pi4kα1 knockout regarding pollen development.

None of the efop single mutant presented any pollen morphological defects or distortion of segregation likely because of redundancy between the four members of this family (Supplemental Figure 4B). We thus generated all the possible combinations of double mutants (Supplemental Figure 4B; Table 2). We were able to obtain *efop2efop3* double homozygous mutants, suggesting no strong synthetic lethality. However, they presented from 19% to 25% of shrivelled pollen grains, resembling those of the *pi4kα1* and *hyc1* mutants, and from 43% to 65% of deformed pollens (resembling those of *npgr1npgr2* double mutants). Thus, depending on the alleles, these double mutant combinations presented from 70% to 90% of abnormal pollens (Supplemental Figure 4A and B). In addition, it was not possible to generate *efop3efop4* double homozygous mutant no matter the alleles used. Indeed, reciprocal crosses indicated 0% of transmission of *efop3* mutant allele when *efop3+/−efop4−/−* plants were used as male, revealing that efop3efop4 pollens were lethal (Table 2). SEM showed that about 45% of the pollen grains present abnormal morphology (Figure 4A and B; Supplemental Figure 4A and B). Finally, observation of efop3-1efop4-2 shrivelled pollen grain by electron transmission microscopy revealed a thick cell wall similar to the phenotype observed for *pi4kα1-1* and *hyc1*. *efop2efop3* and *efop3efop4* double mutants mimic partially and fully *pi4kα1* and *hyc1* pollen phenotypes, respectively. Altogether, our genetic analyses indicate that all the protein classes in the putative PI4Kα1 complex are essential for the male gametophyte in Arabidopsis and that certain mutant combinations in the NPG, HYC or EFOP families either fully (*hyc1+/−, efop3+/−efop4−/−*) or partially (*npg1+/−, npg1+/−npgr1−/−, npgr1−/−npgr2−/−, npg1-2+/− npgr1−/−npgr2-2−/−, efop2efop3−/−*) mimic the *pi4kα1* phenotype. Thus, these proteins likely act together in plants, potentially as a single protein complex.

### Disturbing the PI4Kα1 complex results in various sporophytic phenotypes

While HYC1 is specifically expressed in pollen and is male sterile, HYC2 is predicted to be expressed in the sporophyte which suggests that HYC2 loss-of-function could lead to sporophytic phenotypes. We characterized two T-DNA alleles corresponding to two putative *hyc2* loss-of-function mutants. The segregation rate of *hyc2-2* heterozygous plants was of 60% (Table 1). Moreover, it was not possible to retrieve homozygous plants in the self-progeny of both *hyc2-2* and *hyc2-3*. Reciprocal crosses indicated a transmission of the allele through the male and the female gametophytes even if a weak distortion could be observed in both cases (Table 2). Siliques from *hyc2-2* and *hyc2-3* heterozygous plants presented around 25% to 30% of aborted seeds (Figure 5A-B; Supplemental Figure 5A). Observations of the embryos after clearing showed that in those siliques, some embryos stopped their development at the globular stage before degenerating (likely corresponding to homozygous hyc2 mutant embryos) while the rest of the embryos pursued their development normally (likely corresponding to wild-type and *hyc2+/−* embryos) (Figure 5C). This phenotype was lost and homozygous mutant plants were obtained when HYC2-mCITRINE or HYC2-2xmCHERRY were reintroduced under the control of the HYC2 promoter (Figure 5B; Supplemental Figure 5A-B). Thus, the loss of *HYC2* leads to embryo lethality at the globular stage, suggesting that *HYC1* is essential for the male gametophyte, while *HYC2* is essential for embryogenesis. These results are consistent with the idea that the four-subunit PI4Kα1 complex is essential in plants beyond the development of the male gametophyte.

**Figure 5.**
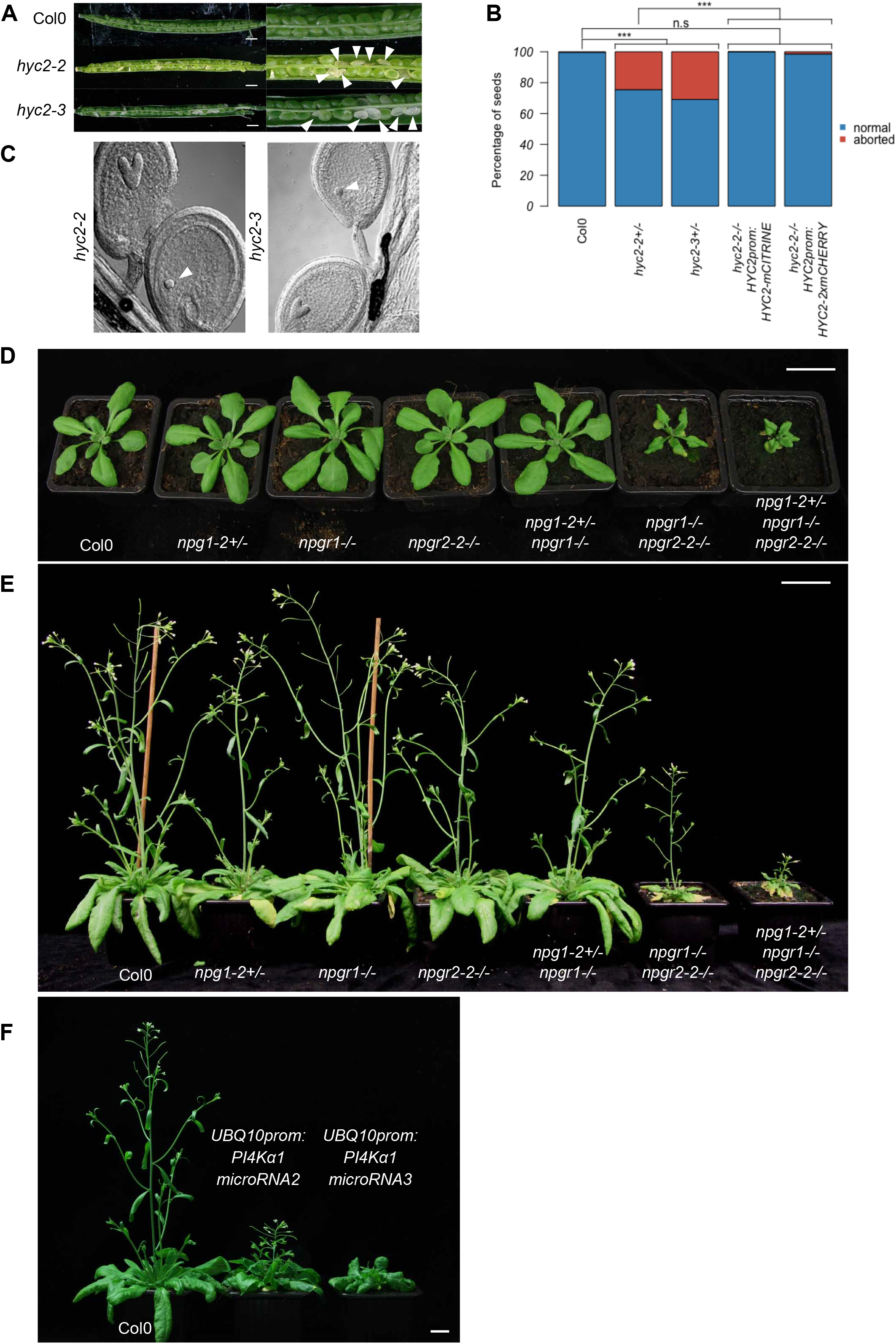
Mutations of the PI4Kα1 complex induce sporophytic phenotypes. (A) Opened siliques of self-fertilized Col-0, *hyc2-2* and *hyc2-3* heterozygous mutant plants. White arrowheads indicate aborted seeds. (B) Percentage of aborted seeds in Col-0, *hyc2-2+/−, hyc2-3+/−, hyc2-2−/− HYC2prom:HYC2-mCITRINE* (insertion n°10), and *hyc2-2−/− HYC2prom:HYC2-2xmCHERRY* (insertion n°11) siliques. The number of seeds counted is superior at 250 for each genotype. Statistics used chi-square test. n.s, non-significant; ***, p<0.001. (C) Cleared seeds from hyc2-2 and hyc2-3 heterozygous mutant plants. White arrowheads indicate globular embryos that have stopped development. (D) Twenty-seven-days-old Col-0, *npg1-2+/−, npgr1−/−, npgr2-2−/−, npg1-2+/− npgr1−/−, npgr1−/− npgr2-2−/−* and *npg1-2+/− npgr1−/− npgr2-2−/−* plants. Scale bar: 2 cm (E) Forty-one-days-old Col-0, *npg1-2+/−, npgr1−/−, npgr2-2−/−, npg1-2+/− npgr1−/−, npgr1−/− npgr2-2−/−* and *npg1-2+/− npgr1−/− npgr2-2−/−* plants. Scale bar: 2cm (F) Thirty-eight-days-old Col-0 and plants expressing microRNA2 and microRNA3 against *PI4Kα1*. Scale bar: 2cm

The lethality of *pi4kα1* or hyc knockouts did not allow us to further study the role of *PI4Kα1* at the cellular and developmental levels in plants. However, some combinations of npg mutants presented growth defect phenotypes. Indeed, *npgr1npgr2-2* double mutant and *npg1+/−npgr1−/−npgr2-1−/−* presented a mild growth phenotype while *npg1+/−npgr1−/− npgr2-2−/−* was dwarf (Figure 5D-E; Supplemental Figure 5C-D). Reintroduction of *NPGR1prom:NPGR1-mCITRINE* was able to rescue the growth phenotype of *npgr1npgr2-2* double mutant (Supplemental Figure 5E-F). This suggests that the PI4Kα1 complex is essential not only for pollen and embryo development but also for later developmental and growth processes. To confirm this hypothesis, we developed a knockdown strategy using a total of 4 independent artificial microRNAs targeted against *PI4Kα1*. Primary transformants expressing ubiquitously the artificial microRNAs number 2 and 3 showed strong growth defects (Figure 5F). The ubiquitous expression of the PI4Kα1 microRNA lines 2 and 3 lead to phenotypes with variable strength. However, most of the plants died or were sterile, which confirmed the essential role of the PI4Kα1 complex not only in gametophytes but also for sporophytic development. To avoid the problem caused by lethality, we next put the artificial microRNA2, which had the strongest phenotype when constitutively expressed, under the control of a β-estradiol-inducible ubiquitous promoter (*UBQ10prom-XVE*) (Siligato et al., 2016). Induction of this artificial microRNA2 led to a decrease in the amount of the *PI4Kα1* protein and the corresponding seedlings exhibited shorter primary roots (Figure 6A).

**Figure 6.**
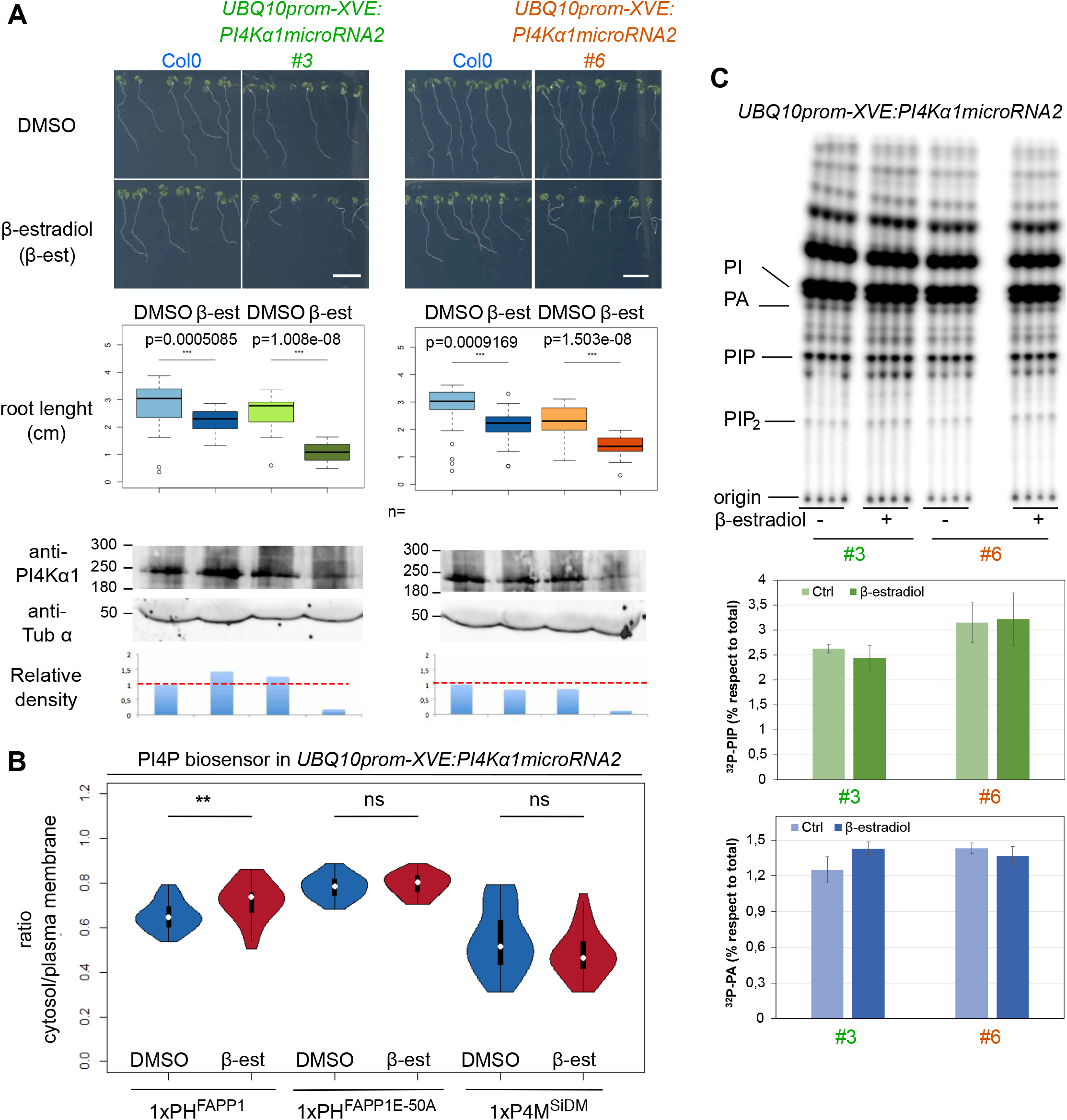
Inducible *PI4Kα1* knockdown impacts sporophytic development and has subtle effects on PI4P accumulation at the plasma membrane. (A) Pictures and measures of the primary root of 9-day-old seedlings of Col-0 and seedlings expressing the inducible microRNA2 against *PI4K α1* on non-inducible (DMSO) and inducible medium (5μM β-estradiol, β-est) supplemented with sucrose for two independent insertions (#3 in green and #6 in orange). Scale bars: 1cm. n indicates the number of seedlings measured. Statistics were done using Wilcoxon test. Bottom panel, PI4Kα1 protein levels of 9-day-old Col-0 seedlings and seedlings expressing the inducible microRNA2 against PI4Kα1 on non-inducible (DMSO) and inducible medium (5μM β-estradiol) for two independent insertions (#3 and #6). Western blot used an anti-PI4Kα1 antibody and an anti-tubuline α (anti-Tub α) as control. The relative density of signal adjusted to the Col-0 DMSO condition is indicated. (B) Cytosol/Plasma membrane signal intensity ratio of PI4P biosensors mCITRINE-1xPHFAPP1 (P5Y), mCITRINE-1x PHFAPP1-E50A and mCITRINEP4MSidM on epidermal root cells of seedlings expressing the inducible microRNA2 against *PI4Kα1* on non-inducible (DMSO) and inducible medium (5μ M β-estradiol) without sucrose. n indicates the number of seedlings measured. Three cells per seedling were measured. Statistics were done using Wilcoxon test. (C) TLC analysis of lipid extract from 9 days-old 32Pi-prelabeled seedlings expressing the inducible microRNA2 against *PI4Kα1* on non-inducible (DMSO) and inducible medium (5μM β-estradiol) supplemented with sucrose for two independent insertions (3 and 6). 32Pi labeling was O/N (16-20 hrs), and quantification of 32P-levels in PIP and PA (as control) by phosphoimaging and calculated as percentage of total 32P-lipids. Each sample represents the extract of three seedlings and each treatment was performed in quadruplicate of which averages ± SD are shown.

With this tool in hand, we next evaluated the effect of PI4K α1 knockdown on the PI4P pool using several approaches. First, we studied the impact of PI4Kα1 knockdown on the localization of PI4P biosensors by calculating the ratio of fluorescence intensity between the cytosol and plasma membrane (Figure 6B). Upon induction, the cytosol/plasma membrane ratio of the PI4P sensor 1xPHFAPP1 (also known as P5Y, Simon et al., 2016) increased, although a significant amount of the sensor was still localized at the plasma membrane (Figure 6B). This result suggests that the amount of PI4P at the plasma membrane was reduced upon inducible PI4Kα1 knockdown (Figure 6B). However, the cytosol/plasma membrane ratio remained unchanged for two additional PI4P biosensors, 1xPHFAPP1-E50A and P4M (Simon et al., 2016). These two biosensors have a high affinity for the plasma membrane pool of PI4P, while 1xPHFAPP1 interacts with PI4P as well as with the TGN/EE-localized protein ARF1 (Simon et al., 2014, 2016). These results thus suggest that the decrease of PI4P at the plasma membrane is relatively modest. Indeed, it is not sufficient to perturb the subcellular targeting of pure PI4P-binding proteins to the plasma membrane (such as 1xPHFAPP1-E50A and P4M), but it can impact the localization of coincident detectors such as 1xPHFAPP1 (which interacts both with PI4P and ARF1). Consistently, the amount of 32P-PIP in lipid extracts of 32Pi-prelabeled seedlings grown on β-estradiol was unchanged compared to the non-treated control (Figure 6C).

Together, these data confirm that PI4Kα1 is acting at the plasma membrane in the production of PI4P. They also suggest that our PI4Kα1 knockdown strategy is not sufficient to strongly impact the pool of PI4P at the plasma membrane. We reason that the remaining PI4Kα1 may sustain PI4P production in this compartment, although we cannot exclude the possibility that other enzymes participate in the PI4P production upon PI4Kα1 knockdown. Importantly, relatively subtle changes in PI4P amount at the plasma membrane (Figure 6B) already lead to visible developmental phenotypes (Figure 6A). This suggest that stronger depletion of PI4P at the plasma membrane would likely lead to lethality, as observed in pollen grain and embryo of various PI4Kα1 complex mutants. Together, our results indicate that the PI4Kα1-dependent pool of PI4P at the plasma membrane is likely required for cellular life in plants, including during gametophyte, embryonic and post-embryonic development.

### The PI4Kα1 complex is associated with the plasma membrane

To confirm the localization of PI4Kα1 at the plasma membrane, we first raised stable transgenic lines expressing PI4Kα1 tagged with mCITRINE at either its N- or C-terminal ends and the red fluorescent protein 2xmCHERRY at the C-terminal end under the control of either its own promoter or the UBQ10 promoter. Consistent with the hypothesis that PI4Kα1 acts at the plasma membrane, the three constructs mCITRINE-PI4Kα1, PI4Kα1-mCITRINE, and PI4Kα 1-2xmCHERRY localized at the plasma membrane and in the cytosol in root epidermal cells and in pollen grains (Figure 7A, Supplemental Figure 6). However, the introduction of *PI4Kα1:PI4Kα1-mCITRINE, PI4Kα1:mCITRINE-PI4Kα1* or *PI4Kα1:PI4Kα1-2xmCHERRY* constructs in the *pi4kα1-1+/−* mutant background failed to complement the pollen lethality and we never recovered pi4kα1-1−/− plants (Table S3). We used the same *PI4Kα1* promoter used for the rescue experiment with the untagged PI4Kα1 (Figure 3B-C; Supplemental Figure 3A), suggesting that PI4Kα1 fused with a fluorescent protein is non-functional. Expression of PI4Kα1 fused to smaller tags (i.e. PI4Kα1-6xHA or Flag-PI4Kα1) also failed to complement *pi4kα1-1* (Table S3).

**Figure 7.**
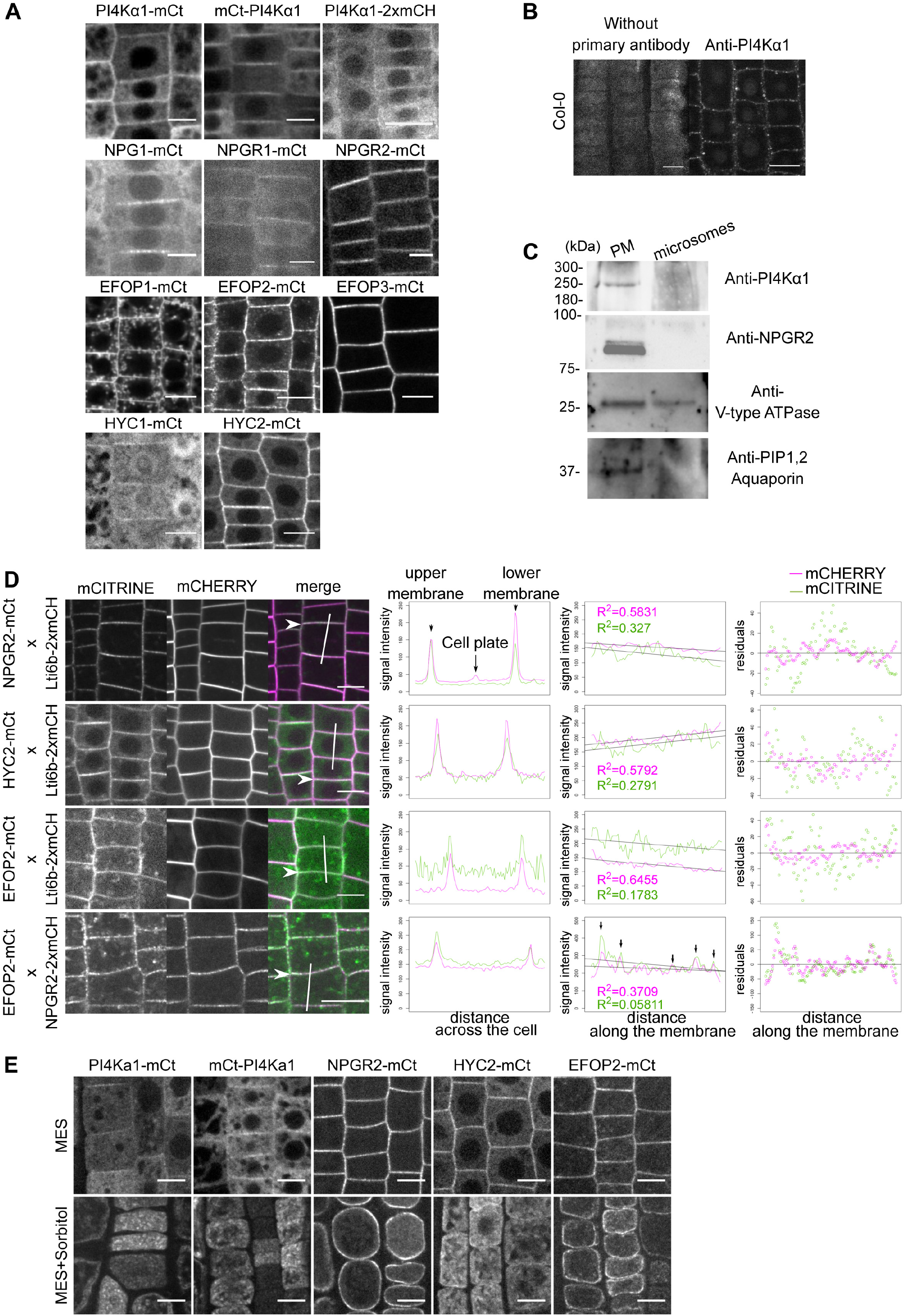
The PI4Kα1 complex localizes at the plasma membrane. (A) Confocal images of PI4Kα1, NPG1, NPGR1, NPGR2, HYC1, HYC2, EFOP1, EFOP2 and EFOP3 fused to mCITRINE (mCt) or mCHERRY under the control of the UBQ10 promoter in root epidermal cells. Scale bars: 10μm. (B) Confocal images of PI4Kα1 using an anti-PI4Kα1 antibody in epidermal root cells on WT seedlings. Control background without primary antibody is shown. Scale bar: 10μm. (C) Western blot using anti-PI4Kα1, anti-NPGR2, Anti-V-type ATPase, and anti-PIP1,2 aquaporin antibodies on plasma membrane and microsomal fractions from WT seedlings. (D) Confocal images of seedlings co-expressing Lti6b-2xmCHERRY (under the control of the 2x35S promoter), NPGR2-mCITRINE, HYC2-mCITRINE or EFOP2-mCITRINE (under the control of the UBQ10 promoter). Graphics on the left represent intensity of each signal across the cell along the white line. Graphics in the middle represent intensity of each signal along the membrane indicated by the white arrow. Matching pic intensity are indicated by black arrows. Linear regression and adjusted R square for each signal are indicated. Graphics on the right represent residuals in y between signal along the membrane and linear regression. (E) Confocal images of PI4Kα1-mCt, mCt-PI4Kα1, NPGR2-mCt, HYC2-mCt, EFOP2-mCt under the control of the UBQ10 promoter in root epidermal cells in control condition (MES) and during plasmolysis (MES+Sorbitol). Scale bars: 10μm.

To confirm the localization obtained with mCITRINE fusion, we used the antibodies against the native PI4Kα1 and performed whole mount immunolocalization in roots. Similar to the mCITRINE-PI4Kα1 and PI4Kα1-mCITRINE fusions, we observed again a signal at the plasma membrane (Figure 7B). To further confirm the preferential association of PI4Kα1 with the plasma membrane, we used cellular fractionation and PEG/Dextran phase partition of whole seedlings and compared the signal obtained on a purified plasma membrane or whole microsomal fractions. We confirmed the presence of proteins in the two fractions using an antibody against V-type ATPase. The purity of the plasma membrane fraction was evaluated with antibodies against the PIP1,2 aquaporin, a known plasma membrane resident protein (Figure 7C). When loading the same amount of protein in each fraction, this experiment revealed the presence of a band around 225kDa, corresponding to PI4Kα1 in the plasma membrane fraction and only a very faint signal in the total microsomal fraction, showing that PI4Kα1 is enriched in the plasma membrane fraction (Figure 7C). Together, fluorescent fusion, immunolocalization and cellular fractionation showed that PI4Kα1 is associated with the plasma membrane.

We next addressed the subcellular localization of NPG, HYC and EFOP proteins at the plasma membrane. NPG1 was previously found to be an extracellular protein in pollen grains (Shin et al., 2014). In our hand, NPG1-mCITRINE, NPGR1-mCITRINE and NPGR2-mCITRINE localized at the periphery of the cell in root meristem (Figure 7A). In addition, NPGR1-mCITRINE and NPGR2-mCITRINE were found at the plasma membrane in pollen grains (Supplemental Figure 6). To confirm this localization and make sure to distinguish between the plasma membrane and cell wall, we co-expressed NPGR2-mCITRINE with Lti6b-mCHERRY. We observed that the two signals perfectly colocalized indicating that NPGR2 is present at the plasma membrane (Figure 7D). Furthermore, we performed plasmolysis of the epidermal root cell by addition of sorbitol. In this context, the plasma membrane detaches from the cell wall. We observed that the signal of NPGR2-mCITRINE remains at the plasma membrane and is not present in the cell wall (Figure 7E). Moreover, in western blot using an anti-NPGR2 antibody, NPGR2 was found enriched in the plasma membrane fraction compared to the microsomal fractions (Figure 7C).

Similarly, HYC1-mCITRINE, HYC2-mCITRINE, EFOP1-mCITRINE, EFOP2-mCITRINE and EFOP3-mCITRINE expressed under the control of the *UBQ10* promoter were found at the plasma membrane in root epidermal cells (Figure 7A) and in pollen grains at the exception of HYC2 and EFOP2 that where highly cytosolic in pollen grains (Supplemental Figure 6). In addition to the plasma membrane localization, we noticed that EFOP1 and EFOP2 were associated with intracellular compartments in epidermal root cells, which were more prominently labelled for EFOP1 than EFOP2.

Upon plasmolysis, PI4Kα1-mCITRINE, mCITRINE-PI4Kα1, HYC2-mCITRINE and EFOP2-mCITRINE signals are found inside the cell and absent from the cell wall. EFOP2 remained associated with the plasma membrane while PI4Kα1 and HYC2 was delocalize to internal compartments (Figure 7E). In any case, all the protein classes in the putative PI4Kα1 complex were associated to some extent with the plasma membrane in both the root meristem and mature pollen grain.

### The PI4Kα1 complex is present in plasma membrane nanodomains

Using confocal microscopy, we noticed that for several of the translational reporters of the PI4Kα1 complex, the signal at the plasma membrane was not continuous, raising the question of a possible subcompartimentalization of the proteins. This is notably the case for PI4Kα1-mCITRINE, NPG1-mCITRINE, NPGR2-mCITRINE, HYC2-mCITRINE, EFOP2-mCITRINE, and EFOP3-mCITRINE (Figure 7A and D and Figure 8A). A similar discontinuous pattern at the plasma membrane was also evident from PI4Kα1 immunolocalization (Figure 7A). Using plant co-expressing Lti6b-2xmCHERRY and NPGR2-mCITRINE, HYC2-mCITRINE or EFOP2-mCITRINE, we observed that NPGR2-mCITRINE, HYC2-mCITRINE and EFOP2-mCITRINE signals along the plasma membrane is less homogeneous than Lti6b, and accumulated in patches of high intensity (Figure 7D). We calculated the linear regression of each signal along the membrane and observed that the R square of NPGR2-mCITRINE, HYC2-mCITRINE and EFOP2-mCITRINE are always smaller than the one of Lti6b-2cmCHERRY indicating a higher dispersion of the intensity. In addition, plants co-expressing NPGR2-2xmCHERRY and EFOP2-mCITRINE show similar intensity pattern with signals partially localized along the plasma membrane (Figure 7D). Similarly, when we observed the plasma membrane in tangential sections, NPGR2-2xmCHERRY and EFOP2-mCITRINE subdomains were partially colocalized (Figure 8A). As control, EFOP2-mCITRINE containing plasma membrane domains did not colocalize with the mostly uniformed localization of Lti6b-2xmCHERRY (Figure 7D and Figure 8A). In order to get a better axial resolution, we used Total Internal Reflection Fluorescence (TIRF) microscopy and confirmed that NPGR2-mCITRINE, HYC2-mCITRINE and EFOP2-mCITRINE were present in nanodomains of the plasma membrane (Figure 8B).

**Figure 8.**
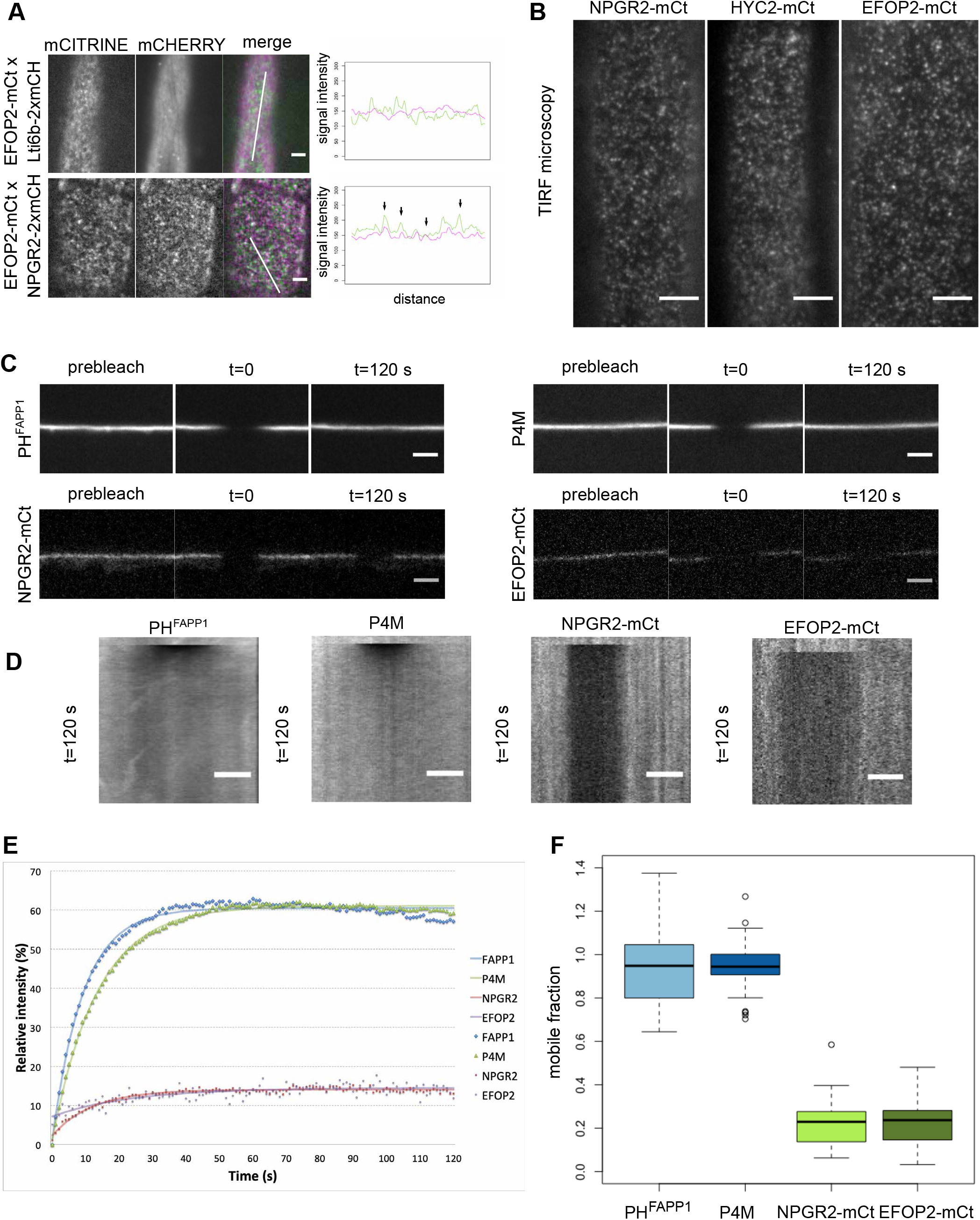
The PI4Kα1 complex localizes in highly static nanodomains at the plasma membrane. (A) Confocal images of seedlings co-expressing Lti6b-2xmCHERRY (under the control of the 2x35S promoter), NPGR2-mCITRINE or EFOP2-mCITRINE (under the control of the UBQ10 promoter). Graphics represent intensity of each signal across the cell along the white line. Black arrows indicate matching signals. Scale bars: 5 μm. (B) Confocal images of TIRF microscopy of NPGR2-mCt, HYC2-mCt and EFOP2-mCt. Scale bars: 5 μm. (C) Confocal images of P4M, PHFAPP1, NPGR2-mCt and EFOP2-mCt before photobleaching (prebleach) and 120s after photobleaching. Scale bars: 5 μm. (D) Kymographs along the membrane for the time laps in (C). Scale bars: 5 μm. (E) Graphic presenting the recovery of the signal intensity over time after bleaching. The number of zones measured is 37, 32, 30 and 13 for P4M, PHFAPP1, NPGR2-mCt and EFOP2-mCt, respectively. The fitting curves are represented. (F) Graphic presenting the mobile fraction of P4M, PHFAPP1, NPGR2-mCt and EFOP2-mCt after 2min postbleaching taking in account the inherent bleaching due to imaging.

### PI4Kα1 complex-containing nanodomains are static at the plasma membrane

Next, we investigated the lateral dynamics of the PI4Kα1 complex at the plasma membrane. To do so, we used fluorescence recovery after photobleaching (FRAP). After bleaching, the signal of NPGR2-mCITRINE, HYC2-mCITRINE and EFOP2-mCITRINE did not recover after 2 min of acquisition (Figure 8C-E; Supplemental Figure 7). In comparison, PI4P sensors (P4M and PHFAPP1) fluorescence recovered in less than a minute after bleaching. Accordingly, the mobile fraction calculated of NPGR2-mCITRINE and EFOP2-mCITRINE was low (around 20%) while the mobile fraction of the sensor reached 100% (Figure 8F). This indicates that if PI4P sensors are rapidly diffusing at the membrane, the PI4Kα1 complex is relatively static. Furthermore, the identical dynamics of NPGR2-mCITRINE, HYC2-mCITRINE and EFOP2-mCITRINE further reinforce the notion that these subunits are part of a single protein complex in vivo.

### EFOPs localize at the plasma membrane via S-acylation lipid anchoring

We next decided to investigate the mechanism by which the PI4Kα1 complex is targeted at the plasma membrane. The four subunits of the PI4Kα1 complex are soluble proteins, without known lipid binding domains. The *efop3-1efop4-4* and *efop3-2efop4-4* mutants showed the same pollen lethality phenotype as the *efop3-1efop4-2* and *efop3-2efop4-2* mutant (Figure 4A-B, Supplemental Figure 4A-B). While *efop4-2* led to a very small-truncated protein (42 aa), the *efop4-4* allele led to a near full-length protein with only a small in frame deletion of 19 residues close to EFOP4 N-terminus. This suggested that this N-terminal region is crucial for EFOP4 function (Figure 9A). Accordingly, the structural model of EFOP1 reveals that the N-terminal part of the protein forms a surface that is highly conserved among plants (Figure 9B). Moreover, this surface is enriched in positively charged amino acids (i.e. high electrostatic potential) suggesting that it could interact with the highly anionic plasma membrane.

**Figure 9.**
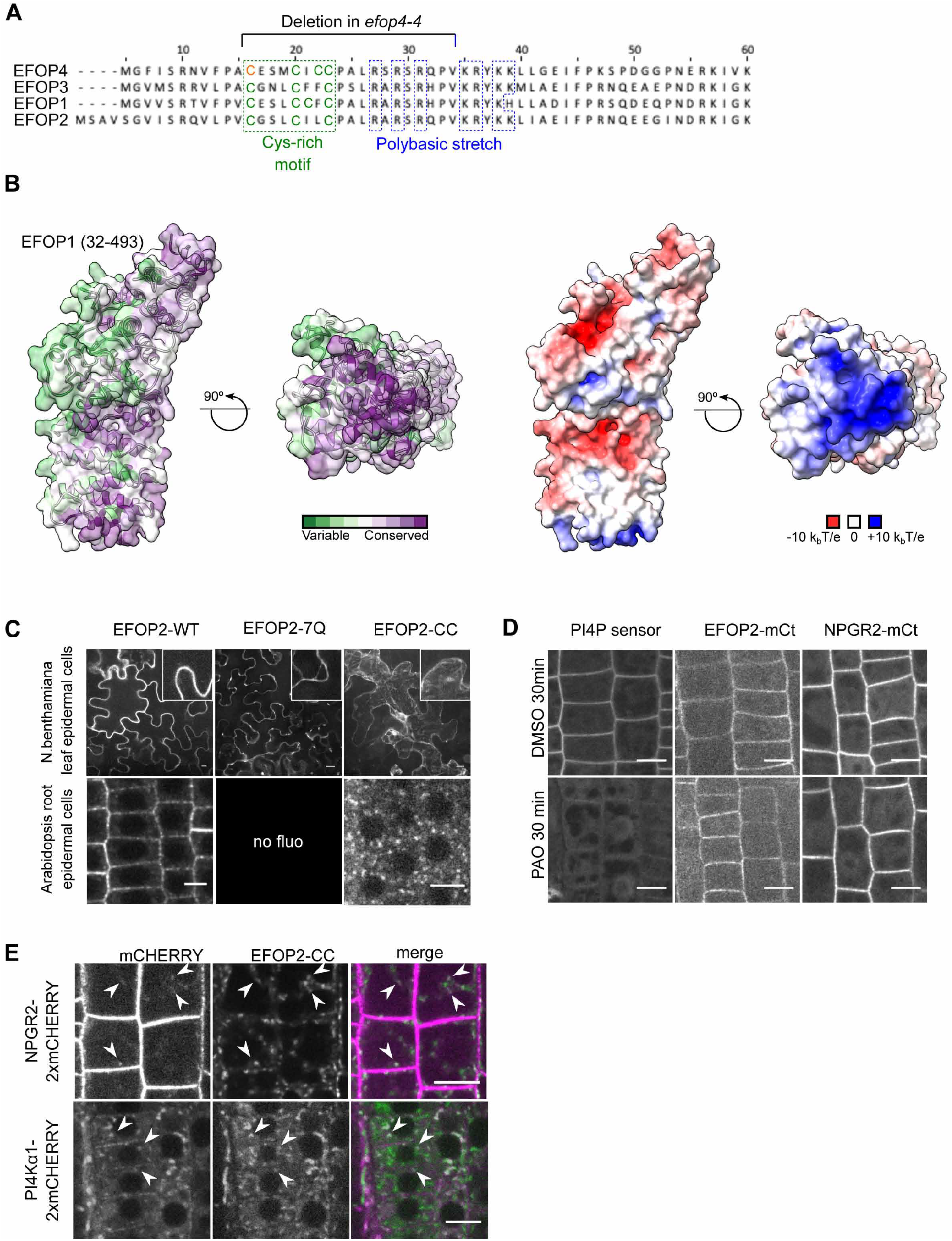
The EFOP subunit determines PI4Kα1 localization by lipid anchoring. (A) N-terminal sequence alignment of EFOPs proteins. Conserved cys-rich motif (green) and polybasic patch (blue) are indicated, as well as, the deletion in efop4-4 CrisPr allele. Bold cysteines are predicted as S-acetylated on the SwissPalm database with high (green) and medium (orange) confidence level. (B) Template-based model of the EFOP1 structure (amino acid range 32-493). The left side shows an analysis of conserved amino acid residues mapped on the solvent-excluded surface of EFOP1. The right side shows electrostatic potential mapped on the solvent-excluded surface of EFOP1. The figure shows that the N-terminally located basic patch is highly conserved. (C) Confocal images of EFOP2-mCITRINE (wild type), EFOP2-7Q-mCITRINE and EFOP2-CC-mCITRINE in N. benthamiana leaf epidermal cells and Arabidopsis root epidermal cells. Scale bar: 10 μm. (D) Confocal images of PI4P sensor (PH domain of FAPP1 carrying the mutation E50A), EFOP2-mCITRINE (EFOP2-mCt) and NPGR2-mCITRINE (NPGR2-mCt) treated for 30 min with 30 μM PAO or the equivalent volume of DMSO. Scale bars: 10μm. (E) Confocal images of Arabidopsis root epidermal cells co-expressing EFOP2-CC-mCITRINE and NPGR2-2xmCHERRY or PI4Kα1-2xmCHERRY. White arrows indicate internal structures where the two signals colocalize. Scale bar: 10μm.

The residues corresponding to the efop4-4 deletion stand astride a non-structured region (1-31 aa) and the positively charged surface visualized in the modelled EFOP structure (Figure 9A and B). These residues are well conserved among the four EFOPs and include both a cysteine-rich motif, which could be S-acylated, and a polybasic region, which could contact anionic lipids at the plasma membrane (Figure 9A). We thus tested the potential role of those two elements in the regulation of EFOP localization and potentially the recruitment of the PI4Kα1 complex at the plasma membrane.

First, we evaluated the role of the polybasic patch in the N-terminus of EFOP proteins. Indeed, this region could be involved in targeting the entire PI4Kα1 complex to the plasma membrane through electrostatic interactions with anionic lipids, notably PI4P. In EFOP2, this region goes from the aa 27 to the aa 39 and contains 7 positively charged residues (Figure 9A). We mutated all lysines/arginines into neutral glutamines and generated a UBQ10prom:EFOP2-7Q-mCITRINE construct. We observed that EFOP2-7Q-mCITRINE was soluble when transiently expressed in Nicotiana benthamiana leaf cells while wild-type EFOP2-mCITRINE was localized to the plasma membrane indicating that polybasic patch in EFOP2 could be essential for plasma membrane targeting (Figure 9C). We next introduced the UBQ10prom:EFOP2-7Q-mCITRINE construct in Arabidopsis epidermal root cells. However, we did not retrieve any lines with a detectable fluorescent signal. It is likely that EFOP2-7Q-mCITRINE is unstable either because of miss folding or because EFOP2 needs to be associated with membrane to remain stable when expressed in Arabidopsis. Finally, we directly investigated the role of PI4P in the recruitment of the PI4Kα1 complex, by using PAO, a PI4K inhibitor. In this condition, the PI4P sensor is detached from the plasma membrane and relocalized in the cytosol (Figure 9D). However, neither NPGR2-mCITRINE nor EFOP2-mCITRINE were mislocalized upon PAO treatment. Thus, PI4P might not be involved in the targeting of the PI4Kα1 complex at the plasma membrane or the depletion of PI4P is not sufficient to delocalize the PI4Kα 1 complex. In any case, this indicates that the presence of the PI4Kα1 complex at the plasma membrane relies, at least in part, on another mechanism.

We then investigated the role of the Cys-rich motif, which was deleted in the efop4-4 allele. Such motif could be a site of S-Acylation; a lipid posttranslational modification that can anchor protein to the plasma membrane (Zaballa and Goot, 2018). Indeed, according to the SwissPalm prediction software, this motif is predicted as S-acetylated with a high (in green) or medium level of confidence (in orange) (Figure 9A). Confirming this hypothesis, all four Arabidopsis EFOP proteins were found to be S-Acylated in a recent proteomic study (Kumar et al., 2020). Notably, all Cys-residues (boxed in Figure 9A) within the Cys-rich region of EFOP3 and EFOP4 were found to be S-acylated with high confidence in planta (Kumar et al., 2020). To experimentally test the importance of such lipid modification in EFOP localization, we mutated the two conserved cysteines (C20 and C23) into serine and generated a UBQ10prom:EFOP2-CC-mCITRINE construct. Similar to EFOP2-7Q-mCITRINE, we observed that EFOP2-CC-mCITRINE was soluble when transiently expressed in Nicotiana benthamiana leaf cells (Figure 9C). Next, we transformed the UBQ10prom:EFOP2-CC-mCITRINE construct into Arabidopsis and found that EFOP2-CC-mCITRINE was not localized at the plasma membrane of root meristematic cells and instead accumulated in intra-cellular structures (Figure 9C). All together, these data suggest that EFOP2 is likely targeted to the plant plasma membrane using lipid acylation anchoring.

### The EFOP/NPG/HYC complex targets PI4Kα1 to the plasma membrane

We then asked if EFOP proteins were sufficient to determine the localization of PI4Kα1 in the cell. Taking advantage of the EFOP2-CC construct localized in intracellular structures, we introgressed NPGR2-2xmCHERRY or PI4Kα 1-2xmCHERRY in Arabidopsis expressing EFOP2-CC-mCITRINE and analysed if EFOP2-CC was able to recruit NPGR2 or PI4Kα1 in those intracellular structures. We observed a weak signal of NPGR2 labelling intracellular compartments that partially colocalized with EFOP2-CC-containing structures. Similarly, PI4Kα1 was not only at the plasma membrane and soluble in the cytosol but also associated with the EFOP2-CC-containing structures (Figure 9E). This showed that EFOP is able to recruit NPGR2 and PI4Kα1 in different compartments of the cell.

Because NPG proteins likely bridges PI4Kα1 to the membrane-binding EFOP subunits, we reasoned that they should contribute to the targeting of PI4Kα1 at the plasma membrane. To test this hypothesis, we monitored the localization of PI4Kα1 in the NPG loss-of-function background. We performed PI4Kα1 immunolocalization on the *npg1-2+/− npgr1−/− npgr2-1−/−* and *npg1-2+/− npgr1−/− npgr2-2−/−* triple mutants in which one allele on NPG1 is still expressed. In the wild type, we observed 38 roots over fours independent experiments and found 33 roots that showed a clear plasma membrane labelling and 5 roots without any labelling. By contrast in the npg triple sesquimutants, we observed that PI4Kα1 was highly soluble and aggregated within the cytoplasm and showed only a faint labelling at the plasma membrane, likely due to the remaining expression of NPG1 (Figure 10A). Such cytosolic/aggregated labelling was observed for 13 roots out of 17 for the *npg1-2+/− npgr1−/− npgr2-1−/−* allelic combination (observed over three independent experiments, three roots with no cytosolic aggregates and one root without any signal) and 26 roots out of 44 for the *npg1-2+/− npgr1−/− npgr2-2−/−* allelic combination (observed over four independent experiments, nine roots with no cytosolic aggregates and nine roots without any signal). This result indicates that PI4Kα1 plasma membrane targeting requires NPG proteins.

**Figure 10.**
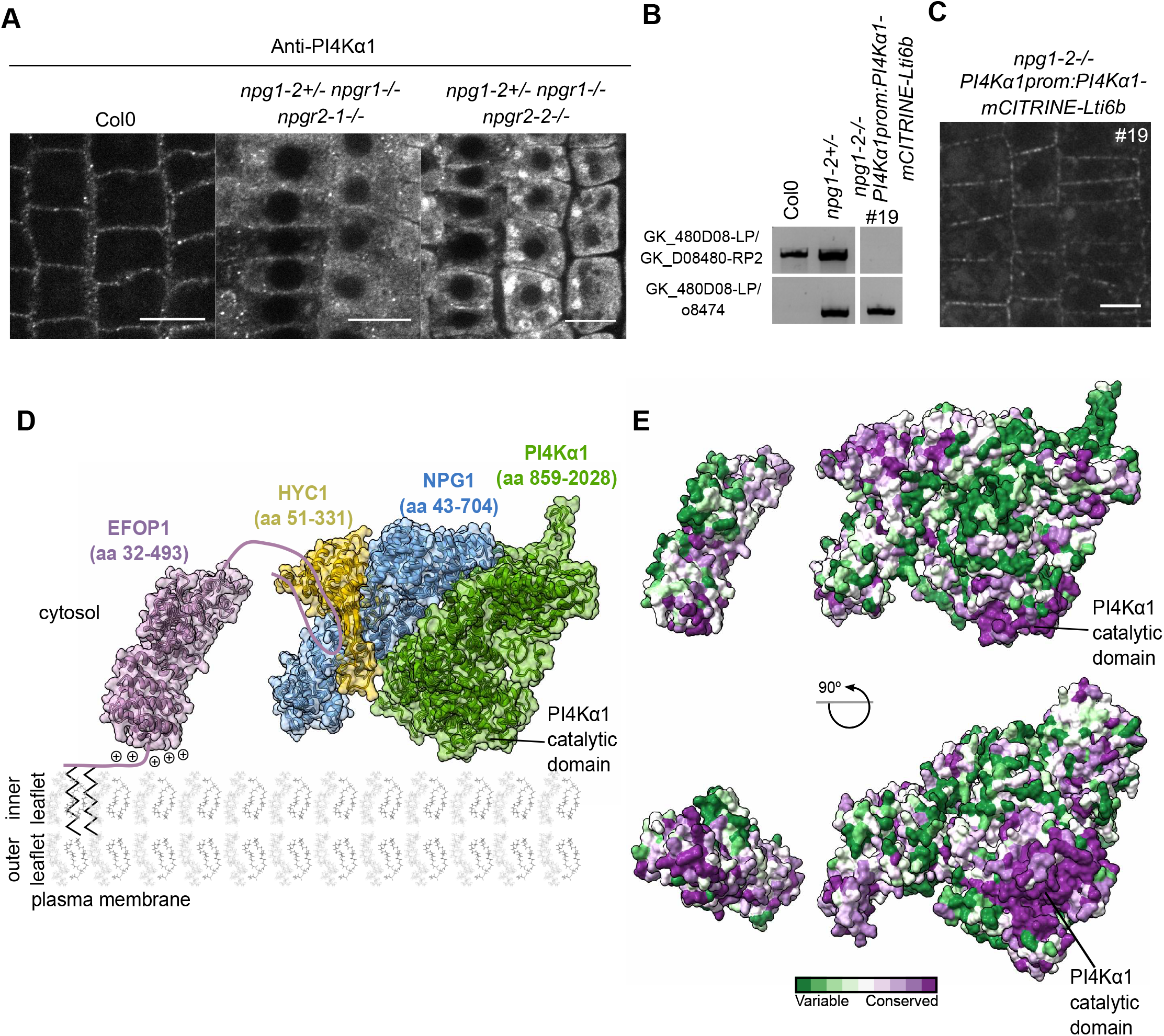
The NPG subunit acts as a scaffold around which HYC and PI4Kα1 structure themselves. (A) Confocal images of PI4Kα1 using an anti-PI4Kα1 antibody on epidermal root cells of WT, n*pg1-2+/−npgr1−/−npgr2-1−/−* and *npg1-2+/−npgr1−/−npgr2-2−/−* seedlings. Scale bar: 10μm. (B) Genotyping of Col0, *npg1-2* heterozygous plants and *npg1-2* homozygous plants complemented with PI4Kα1prom:PI4Kα1-mCITRINE-Lti6b (insertion n°19). Upper panel shows amplification of the gene sequence. Lower panel shows amplification of T-DNA border. (C) Confocal images of PI4Kα1::PI4Kα1-mCITRINE-Lti6b in *npg1-2−/−* background (insertion n°19). (D) Schematic depiction of the plasma membrane targeting of the plant PI4Kα1 complex. The structure of the heterotrimeric complex composed of PI4Kα 1 (green, amino acid region 859-2028), NPG1 (blue, amino acid region 43-704) and HYC1 (yellow, amino acid region 51-331), was obtained using template-based modelling and protein-protein docking. The EFOP1 N-terminal part (purple, amino acid region 32-493) was prepared by template-based modelling. Parts of EFOP1 with no homologous structure or predicted as intrinsically disordered are depicted as continuous lines. The S-acylated N-terminus of EFOP1 is shown together with plus signs depicting the polybasic patch of EFOP1. (E) Analysis of conserved amino acid residues mapped on the solvent-excluded surface of the heterotrimeric complex (PI4Kα1-NPG1-HYC1) and EFOP1.

Using a different approach, we also generated a fusion between PI4Kα1-mCITRINE and the transmembrane protein Lti6b in order to artificially target PI4Kα1 at the plasma membrane and thus bypass the role of the NPG/HYC/EFOP complex. The *PI4Kα1prom:PI4Kα1-mCITRINE-Lti6b* construct was able to rescue the npg1-2 as we were able to recover homozygous *npg1-2* mutants upon expression of this particular transgene (Figure 10B). This indicates that npg1 pollen lethality is likely due to the absence of PI4Kα1 at the plasma membrane during pollen development, thereby confirming 1) the functional link between NPG proteins and PI4Kα1, and 2) that the function of NPG proteins is to target PI4Kα1 to the plasma membrane. As discussed above, tagged version of PI4Kα1, including PI4Kα1-mCITRINE, were not functional as they did not complement the pi4kα1 mutant (Table S3). Similarly, we did not retrieve any complemented pi4kα1 mutant line expressing *PI4Kα1prom:PI4Kα1-mCITRINE-Lti6b* (Table S3). This raised the question on how the non-functional PI4K α1-mCITRINE-Lti6b chimeric construct was able to rescue the npg1-2 mutant. One possibility is that PI4Kα1 naturally forms homodimer. In that scenario, the non-functional PI4Kα 1-mCITRINE-Lti6b would recruit the endogenous –and functional– PI4Kα1 at the plasma membrane in the absence of npg1, and thus complement the npg1 mutant but not the pi4k α1 mutant. We currently do not know whether PI4Kα1 is able to dimerize. However, structural data from the animal field showed that the PI4KαIII complex dimerizes at the plasma membrane (Lees et al., 2017b), suggesting that it could also be the case in plants.

In these plants, PI4Kα1-mCITRINE-Lti6b is located in clusters at the plasma membrane while Lti6b is known as being rather homogenously localized at the plasma membrane (Figure 7D and 10C). The chimeric PI4Kα1-mCITRINE-Lti6b proteins might be restricted in clusters by other factors, which indicates that the subcompartimentalization of PI4Kα1 complex might not be an intrinsic property of the complex but rather come from interactions between PI4Kα1 and other lipids or proteins. Furthermore, it might indicate that the subcompartimentalization of PI4K α1 complex is an essential feature for the proper function of the complex.

Altogether, this study shows that PI4Kα1 form a heterotrimeric complex with NPG and HYC proteins. EFOP proteins target the PI4Kα1 complex to the plasma membrane by a combination of lipid anchoring and electrostatic interactions (Figure 10D). When aligned with the plasma membrane, the PI4Kα1 complex displays an evolutionarily conserved surface composed of the N-terminal basic patch of EFOP, the catalytic domain of PI4Kα1 and the N-terminal part of NPG. This highly conserved surface likely represents a membrane interacting interface of the heterotetrameric PI4Kα1 complex and provides an initial mechanistic insight into the mode of function of PI4Kα1 in plants (Figure 10E).

## DISCUSSION

### The plant PI4Kα1 complex is essential for cell survival

In this study, we showed that the loss-of-function of the PI4Kα1 leads to lethality of the male gametophyte. Similarly, knockouts of HYC1 and EFOP proteins mimic this phenotype supporting the idea that these proteins act as a complex. Surprisingly, npg1 single mutant is pollen lethal but do not present the same morphological defects that pi4k α1 or hyc1 mutants. The combination of loss of function of NPG1, NPGR1 and NPGR2 gives rise to deformed and shrivelled pollen grains indicating that during pollen development the three NPGs are expressed and partially redundant. NPG1 specifically could be needed for later steps of pollen development and germination explaining the pollen lethality of NPG1 despite the absence of morphological defect. Indeed, pollen apertures are established in distinct membrane domains enriched in PI4P and PI(4,5)P2 (Lee et al., 2018). Like npg1 mutant, loss-of-function of SEC3A, a gene coding for a subunit of the exocyst complex, is pollen lethal (Bloch et al., 2016; Li et al., 2017). *sec3a* mutant pollen grains do not present morphological defect yet they do not germinate. SEC3a is recruited at the pollen germination sites and its binding to the plasma membrane depends on positively charged amino acids. Thus, NPG1 could participate in the recruitment of PI4Kα1 at aperture sites, which could be necessary to landmark the germination site, and the subsequent recruitment of SEC3a or other proteins. However, we showed that NPGR2 complements the npg1 phenotype when expressed under the NPG1 promoter, which suggest that the difference between NPG1 and NPGR2 function is mostly at the transcriptional level.

In addition to gametophytic phenotypes, we also observed that the loss of HYC2 induces embryo lethality while npg triple sesquimutants and PI4Kα1 knockdown display severe growth phenotypes at rosette and seedling stages. This is concordant with the idea that PI4Kα1 has a critical role for the cell function, not only during gametophytic but also sporophytic development. PI4P is crucial for the plasma membrane surface charge (Platre et al., 2018; Simon et al., 2016). Thus, it is likely that the loss of the PI4Kα1 affects the membrane surface charge and lead to the mislocalization of a whole set of proteins (Barbosa et al., 2016; Noack and Jaillais, 2020; Simon et al., 2016).

The genome of Arabidopsis codes for two other PI4Kinases, PI4Kβ1 and PI4Kβ2 localized at the TGN/EE and at the cell plate (Kang et al., 2011; Lin et al., 2019; Preuss et al., 2006). As the plasma membrane and the TGN/EE are intimately linked through vesicular trafficking, the interdependence of the TGN/EE and the plasma membrane pool of PI4P remained an open question. pi4kβ1pi4kβ2 double mutant and pi4kα1 mutant display developmental phenotypes, which indicates that these lipid kinases are not fully redundant. This is coherent with the different subcellular localization of PI4Kβ1/PI4Kβ2 and PI4Kα1 in plant cells. However, only PI4Kα1 is essential in plants, while the *pi4kβ1pi4kβ2* double mutant harbours mild phenotypes. This contrasts with yeast, in which the loss of either pik1 or sst4 is lethal (Audhya et al., 2000). It is thus possible that in plants, the lack of PI4Kβs at the TGN is compensated by PI4Kα1 activity, perhaps through the endocytosis of plasma membrane PI4P. In fact, a large portion of the *pi4kβpi4k1β2* double mutant phenotype can be attributed to its function in cytokinesis (Lin et al., 2019). Thus, PI4Kα1 activity is not able to compensate for PI4Kβs function during cell division.

Using inducible PI4Kα1 knockdown, we confirmed a role for PI4Kα1 in the production of PI4P at the plasma membrane. However, using a knockdown strategy, we cannot conclude about the relative contribution of PI4Kα1 in the total PI4P production. Yet, we can speculate about this point. Indeed, we know that 1) the *pi4kβ1pi4kβ2* double mutant, in which both kinases are fully knocked-out, has no detectable diminution of total PI4P (Lin et al., 2019) and 2) the pool of PI4P at the plasma membrane is quantitatively much more abundant than in the TGN (Simon et al., 2016). Together, we can thus hypothesize that PI4Kα1 is likely responsible for most of the PI4P production in plant cells. The fact that PI4Kα1, but not PI4Kβs, is essential for cell survival argues in favour of this model, which nonetheless remains to be experimentally validated. Perhaps the use of inducible or tissue-specific knock-out using the CRISPR-cas9 technology will enable to address this point more directly in the future.

### Cell wall defect in pollen grains

From meiosis to the end of microspores development, the tapetum establishes the exine layers around the pollen grain by secreting sporopollenine, while the intine layer is synthesized by the pollen (Borg and Twell, 2011). Many male sterile mutants have been identified in Arabidopsis and many of them are involved in pollen cell wall formation. This includes defects of secretion, biosynthesis or transport in the tapetal cells or microspores or callose and cellulose deposition by the microspores (Borg and Twell, 2011). Transmission electron microscopy of pi4kα1, npg triple sesquimutants, *hyc1* and *efop3efop4* pollens reveal that they present a thicker an irregular intine layer. Thus, it is likely that the secretion of cell wall materials by the micropores is affected. The intine layer has a composition similar to a primary cell wall with cellulose, hemicellulose and pectin. Loss-of-function mutants for subunit of the cellulose synthase complex -*cesa1* and *cesa3*-are pollen sterile with shrivelled pollen grains similar to the pi4kα1 complex mutants (Persson et al., 2007). In *cesa* mutants, a nonhomogeneous deposition of intine is observed revealing the importance of cellulose microfibrilles to guide intine deposition. The TPLATE complex has also been involved in pollen cell wall deposition (Van Damme et al., 2006). tplate mutants are male sterile with shrivelled pollens due to abnormal deposition of callose in the intine layer. This may be explained by a poor regulation of callose synthase endocytosis in the tplate mutant. It is possible that callose synthase endocytosis mediated by TPLATE complex and/or cellulose is also affected in pi4kα1 complex mutants.

*efop2efop3* mutant also presents pollen morphological defects. However, it was possible to obtain double homozygous plant indicating that invalidating *efop2efop3* does not lead to gametophytic lethality. Therefore, it is possible that the defect observed on the pollen grains do not come from the microspores but from an incorrect regulation of the secretion of material by the tapetum. Another possibility is that the first steps of pollen development before meiosis are impaired in *efop2efop3* mutant. Interestingly, during microspore development, tapetal cells acquire a huge ER compartment that lay beneath the plasma membrane (Borg and Twell, 2011). Mutant involved in ER-Golgi transport such as sec31b that belongs to the COPII complex present similar phenotype that efop2efop3 mutant with a combination of normal, deformed and collapsed pollen grains (Zhao et al., 2016). We can thus speculate that the PI4Kα1-driven production of PI4P at the plasma membrane might directly or indirectly impact the ER structure, dynamics and/or interaction with the plasma membrane. However, this hypothesis will require further investigation.

### Function of the NPG-HYC-EFOP complex

Our study also showed that PI4Kα1’s plasma membrane localization is mediated by interactions with NPG and EFOP proteins rather than by a putative PH domain. At first, this PH domain was thought to localize PI4Kα1 at the plasma membrane through interaction with anionic phospholipids (Stevenson et al., 1998; Stevenson-Paulik et al., 2003; Xue et al., 1999). However, a region corresponding to the putative Arabidopsis PH domain correspond to the helical (cradle) and catalytic domains of human PI4KIIIα (Supplemental Figure 8) (Dornan et al., 2018; Lees et al., 2017). We found that PI4Kα1’s plasma membrane localization depends on the S-acylation of the EFOP proteins. Several S-acylated peptides in addition to the one in the N-terminal region have been found in EFOP proteins (Kumar et al., 2020). As S-acylation is a reversible post-translational modification, differential S-acylation could be a mechanism explaining the different localization observed for EFOP1 and EFOP2 in the cell.

NPGs are bridging PI4Kα1 and HYC with EFOP proteins. In addition, NPGs are calmodulin (Cam)-binding protein. Indeed, NPG1 can interact with several Cam isoforms in presence of Ca2+ and has been suggested to play a role in Ca2+ dependent pollen tube germination (Golovkin and Reddy, 2003). Ca2+ is also intimately connected with phosphoinositides, membrane properties and endomembrane trafficking (Himschoot et al., 2017). Ca2+ can directly bind anionic phospholipids modulating locally membrane electrostatics, preventing or promoting the recruitment of lipid binding proteins, inducing the formation of PI(4,5)P2 clusters and facilitating membrane fusion (Li et al., 2014; McLaughlin and Murray, 2005). As phosphoinositides can bind Ca2+, diffuse in the membrane and release Ca2+ somewhere else, they have been suggested to buffer and modulate Ca2+ signalling at the subcellular level. Ca2+ is also known to regulate many actors of endomembrane trafficking including regulators of the cytoskeleton, TPLATE complex, ANNEXINs and SYNAPTOTAGMINs (Bürstenbinder et al., 2013, 2017; Carroll et al., 1998; Hepler, 2016; Schapire et al., 2008; Van Damme et al., 2006). For instance, the ER-Plasma membrane tethers protein SYT1 contains C2 domains that bind Ca2+ and phosphoinositides (Giordano et al., 2013; Idevall-Hagren et al., 2015; Lee et al., 2019; Pérez-Sancho et al., 2015; Ruiz-Lopez et al., 2020; Yamazaki et al., 2008). The PI4Kα1 complex could be localized at ER-Plasma membrane contact sites and participate in the Ca2+ signalling at this junction through calmodulin binding.

If EFOPs are anchoring the complex at the membrane and NPGs are bridging PI4Kα1 and EFOPs, the role of HYC-CIN-CONTAING proteins in the complex is less clear. In mammals, FAM126A, which has a HYCCIN domain, is involved in the stability of the complex and is used as a scaffold by TTC7 to be shaped around (Dornan et al., 2018). In human, FAM126A mutations lead to severe case of hypomyelination and congenital cataract (Baskin et al., 2016; Miyamoto et al., 2014). However, even complete knockout of FAM126A is not lethal while loss-of-function of PI4KIIIα is (Baskin et al., 2016). This subunit is not present in yeast, suggesting that HYCCIN-domain containing proteins may have an accessory function in the PI4Kα1 complex, rather than an essential one. By contrast, we found that in Arabidopsis both hyc1 and hyc2 mutants are lethal as they are required for male gametophyte and embryo development, respectively. Our result thus suggests that HYC is an essential subunit for the function of the PI4Kα1 complex in plants. In human, there are additional HYCCIN-containing proteins, including FAM126B. Our results thus open the possibility that FAM126A is not the only HYCCIN-containing proteins that may contribute to PI4KIIIα function in Metazoa.

### Formation and possible function of PI4Kα1-containing nanoclusters

The PI4Kα1 complex localizes at the plasma membrane in nanodomains. In yeast, Stt4 is found in large clusters called PIK patches (Baird et al., 2008). However, the clusters of Stt4 do not correlate with clusters of PI4P that probably diffuse laterally in the membrane. Similarly in Arabidopsis, PI4P biosensors are not clustering at the plasma membrane (Simon et al., 2014, 2016; Vermeer et al., 2009). This is in accordance with in vitro data showing that PI4P inhibits the catalytic activity of PI4Kα1 (Stevenson-Paulik et al., 2003). In addition, we showed that these nanodomains are immobile in plants, despite the fluidity of the plasma membrane (Jaillais and Ott, 2020). It is possible that unknown interactions with transmembrane protein, cytoskeleton or lipid nanodomains stabilize the PI4Kα1 complex.

Do these nanodomains correspond to a functional unit? Among many possibilities, they could correspond to ER-plasma membrane contact sites. Indeed, in yeast, Stt4 reside at these contacts (Omnus et al., 2018). Another hypothesis is that they could be a zone of attachment between the plasma membrane and the actin cytoskeleton. PI4Kα1 has been purified from F-actin-rich fraction from carrots and associates with polymerized F-actin in vitro (Stevenson et al., 1998). Additionally, in yeast, Stt4p is necessary for a proper actin cytoskeleton organization (Audhya et al., 2000; Foti et al., 2001). The two hypotheses are not mutually exclusive. Indeed, the actin disorganization phenotype of stt4 mutant is rescued in yeast by the knockout of Sac1p, a PI4P phosphatase that resides in the ER membrane (Foti et al., 2001). Stt4p and Sac1p together control the PI4P gradient at membrane contact sites (Noack and Jaillais, 2020).

PI(4,5)P2 is enriched in nanodomains at the plasma membrane in pollen tubes (Fratini et al., 2020; Kost et al., 1999). These nanodomains contain PIP5K2 and are involved in actin dynamics. We speculate that the PI4Kα1 complex could also be targeted to these nanodomains in order to catalyse a phosphorylation cascade from PI to PI4P to PI(4,5)P2. In this model, PI4Kα1 inhibition by its substrate could help to coordinate the subsequent phosphorylation of PI4P by PIP5K2. In any case, given the absolute requirement on the PI4Kα1 complex for plant cell survival, deciphering the mechanisms behind the precise spatiotemporal regulation of this complex and the associated functions will be a fascinating challenge for the future.

## Acknowledgments

We thank the present and past members of the SiCE group as well as G. Vert and G. Ingram for discussions and comments, the NASC/ABRC collection for providing transgenic Arabidopsis lines, L. Kalmbach and M. Barberon for the gift of pLOK180_pFR7m34GW, A. Lacroix, J. Berger and P. Bolland for plant care. We acknowledge the contribution of SFR Biosciences (UMS3444/CNRS, US8/Inserm, ENS de Lyon, UCBL) facilities: C. Lionet, E. Chatre and J. Brocard at the LBI-PLATIM-MICROSCOPY for assistance with imaging and V. Gueguen-Chaignon, A. Page and F. Delolme at the Protein Science Facility (PSF) for assistance with mass spectrometry. Transmission Electron microscopy images were acquired at the CTμ (Centre Technologique des Microstructures) Université de Lyon1, we acknowledge Xavier Jaurand and Veronica La Padula for technical assistance. YJ. was funded by ERC no. 3363360-APPL under FP/2007-2013 and ANR caLIPSO (ANR18-CE13-0025), M-C.C. by a start-up package “font de recherche – projet émergent” from ENS de Lyon, SM by project PlayMobil ANR-19-CE20-0016-02 and by project PhosphoREM-domain ANR-19-CE13-0021-01, L.C.N. by a PhD fellowship from the French Ministry of Higher Education and R.P. by the Czech Science Foundation grant 19-21758S.

## Author contributions

L.C.N. was responsible of all the experiments described in this manuscript with the following exception: V.B. performed transmission electron microscopy experiments and helped with microscopy image acquisition and analyses; F.R. performed and imaged whole mount immunolocalization; A.M-C. and S.M. purified plasma membranes; M-C.C. initiated the yeast-two hybrid screen; L.A. helped with yeast-two hybrid experiments; R.P. performed structural modelling, F.D.S. and T.M. measured 32P PIP levels. L.C.N. prepared the figures, L.C.N. and Y.J. wrote the manuscript and all authors commented on the manuscript.

## Conflict of Interest

The authors declare that they have no conflict of interest

## Material and methods

### Plant Material

*Arabidopsis thaliana,* ecotype Columbia (Col0) l was used in this study, except for *pi4kα1-2*, which is in Ws background. The plants expressing *2xp35S:myrimCITRINE-mCITRINE* were obtained from Jaillais et al., 2011. The PI4P sensors lines, *pUBQ10::mCITRINE-1xPH^FAPP1^* and *pUBQ10:mCITRINE-P4M^SidM^* were obtained from (Simon et al., 2014, 2016) **(Table S3)**.

### In vitro culture conditions

Seeds were surface-sterilized by addition of 4ml of HCl 37% in 100mL of bleach for four hours before plating on Murashige and Skoog (MS, Duchefa Biochemie®) media supplemented with 0.8% plant agar and containing the appropriate antibiotic or herbicide. Glufosinate, Kanamycin, Hygromycin and Sulfadyacin have been used at 10mg.L^−1^, 50mg.L^−1^, 30mg.L^−1^ and 75mg.L^−1^ respectively. Plates were placed under continuous white light conditions for 7 to 10 days. Resistant and sensitive seedlings were counted for segregations.

### Culture condition on soil

Seeds were directly sown in soil or 7-day-old seedlings were transferred from *in vitro* plates to soil. Plants were grown at 21°C under long day condition (16 hrs light, LED 150μmol/m^2^/s).

### Sequence Analysis

Sequence alignments have been performed using the Muscle WS software and edited with Jalview 2.0 software. Clustal colour code has been used. Domains have been identified using the SMART (Simple Modular Architecture Research Tool) software. Predicted lipid modification sites have been found using the GPS-Lipid software.

### Structure Modeling

To model a structure of individual PI4Kα1 complex subunits, we used several different threading algorithms, namely Swiss-model (Waterhouse et al., 2018), HHPred(Zimmermann et al., 2018) and RaptorX (Källberg et al., 2012). All three threading algorithms lead to highly similar results. For subsequent protein-protein docking and analyses, we used the models calculated using the Swiss-model program as the obtained models had the best ProSA score (Wiederstein and Sippl, 2007). Next, we utilized a hybrid protein-protein docking algorithm implemented in the HDOCK program (Yan et al., 2020) to position PI4Kα1 to NPG1 and NPG1 to HYC1. To analyze amino acid conservation, we employed the Consurf server with the default parameters (Ashkenazy et al., 2016). The electrostatic potential of EFOP1 was calculated by solving the nonlinear Poisson-Boltzmann equation using the APBS-PDB2PQR software suite (Baker et al., 2001; Dolinsky et al., 2004). The structures were visualized in the ChimeraX program (Pettersen et al., 2021).

### Cloning of reporter lines

Lti6b/pDONR207, UBQ10prom/pDONR P4-P1r, 2X35S/pDONR P4-P1r, mCITRINE/pDONR 221, mCITRINE/pDONR P2R-P3, 2xmCHERRY-4xmyc/pDONR P2R-P3 entry vectors were obtain from Elsayad et al., 2016; Jaillais et al., 2011; Marquès-Bueno et al., 2016; Simon et al., 2014, 2016 **(Table S4)**. For other entry vectors, coding gene or genomic gene sequences and 3’UTR sequences were cloned using Gateway® technology (Invitrogen®). Promoter sequences and Lti6b-FRB were cloned by Gibson Assembly method (Biolab®). The coding sequences were amplified by PCR using the high fidelity Phusion Hot Start II (Thermo Fisher Scientific®) taq polymerase and the indicated primers and template **(Table S5)**. The PCR products were purified using the NucleoSpin Gel and PCR Clean kit (Macherey-Nagel®) and recombined in the indicated vector using BP Clonase™ II Enzyme Mix (Invitrogen®) or Gibson Assembly mix (Biolab®) **(Table S5)**. Thermocompetent DH5α *E. coli* were transformed with the corresponding vectors and plate on LB (Difco™ LB Broth, Lennox, #214010, 20 g/L, 15% agar, Difco™ Bacto Agar) containing the Kanamycin at 50mg.L^−1^. Plasmids were purified using the Nucleospin Plasmid kit (Macherey-Nagel®) and inserts sequenced. Expression vectors containing the promoter-gene-fluorescent tag cassette or 3’UTR were obtained by using LR clonase-based three-fragment recombination system (Invitrogen®), the pB7m34GW/pH7m34GW/pK7m34GW/pS7m43GW/ pLOK180_pR7m34g (gift from L. Kalmbach) destination vectors, and the corresponding entry vectors **(Table S3–S4)**. Only the EFOP2-7Q was introduce in the destination vector pK7FWG2 using the LR clonase-based one-fragment recombination system (Invitrogen®) (Karimi et al., 2002). Thermocompetent DH5α cells were transformed and selected on LB plate (Difco™ LB Broth, Lennox, #214010, 20 g/L, 15% agar, Difco™ Bacto Agar) containing 100mg.L^−1^ of Spectinomycin. Plasmids were purified using the Nucleospin Plasmid kit (Macherey-Nagel®) and inserts sequenced.

### Site directed mutagenesis

Plasmids were amplified by PCR using the high fidelity Phusion Hot Start II (Thermo Fisher Scientific®) taq polymerase and the indicated primers carrying mutations **(Table S6)**. PCR products were digested using and cutsmart buffer (Biolab®). Thermocompetent DH5α *E. coli* were transformed with the digested PCR product and plate on LB (Difco™ LB Broth, Lennox, #214010, 20 g/L, 15% agar, Difco™ Bacto Agar) containing the appropriate antibiotic. Plasmids were purified using the Nucleospin Plasmid kit (Macherey-Nagel®) and mutations were sequenced.

### Artificial microRNA designed

Artificial microRNAs were designed using Web MicroRNA Designer (http://wmd3.weigelworld.org/cgi-bin/webapp.cgi). DNA synthesis containing AttB1-AttB2 sites and the miR319a modified with the sequence of the artificial microRNA (Table S9) were ordered at Integrate DNA TechnologyR (IDT). The DNA were recombined in the pDONR 221 (InvitrogenR) using BP ClonaseTM II Enzyme Mix (InvitrogenR).

### Agrobacterium transformation

Electrocompetent *A. tumefaciens* (C58pmp90) were transformed by electroporation. 1μL of DNA plasmid at a concentration of 0.25-1 μg/μl was added into 50 μL of electrocompetent agrobacterium on ice. The agrobacterium were transfered into cold 1mm wide electroporation chamber (Eurogentec, #CE00150). A pulse of 2 kV, 335 Ω, 15μF, for 5 ms was performed on the electroporation chamber using the MicropulserTM (Bio-Rad, #165-2100). 1 mL of liquid LB media was added and the bacteria were placed into a new tube and incubated at 29°C for 2-3h. The agrobacterium were selected on LB (Difco™ LB Broth, Lennox, #214010, 20 g/L, 15% agar, Difco™ Bacto Agar) or YEB (0.5% beef extract, 0.1% yeast extract, 0.5% peptone, 0.5% sucrose, 1.5% bactoagar, pH7.2) plates containing the appropriate antibiotics to select the agrobacterium strain (50mg.L-1 of Rifampicin and 20mg.L-1 of Gentamycin) and the target construct (250mg.L-1 of Spectinomycin). Plates were incubated at 29°C for 48h.

### Plant transformation and selection

Agrobacterium were spun and resuspended in 5% sucrose and 0.02% Silwet L-77 detergent. Arabidopsis were transformed by floral dipping.

Transformed seedlings were selected on Murashige and Skoog (MS, Duchefa Biochemie®) media supplemented with 0.8% plant agar containing the appropriate antibiotic or herbicide. Basta, Kanamycin, Hygromycin and Sulfadiazin have been used at 10mg.L^−1^, 50mg.L^−1^, 30mg.L^−1^ and 75mg.L^−1^ respectively. Selection of the pLOK180_pR7m34g plasmid was done using the FastRed method (i.e. red fluorescent seeds).

### Protein extraction, immunoprecipitation and western blot analysis

Leaf tissue from wild-type plants and/or transgenic lines were collected, frozen in liquid nitrogen and ground to powder. The powder was resuspended in −20°C MetOH; 1% Protease inhibitor (P9599, Sigma-Aldrich®) and incubated at −20°C for 5min. Proteins were pelleted, resuspended in −20°C acetone and incubated at −20°C for 5min. Proteins were pelleted. The pellet was dried and resuspended in Protein Extraction Reagent (C0856, Sigma-Aldrich®) supplemented with 1% Protease inhibitor (P9599, Sigma-Aldrich®). Protein extraction were then kept at −20°C.

Protein samples in 1xSDG (Tris-HCL 0.25 M, 10% glycerol, 2% DTT, 2.5% SDS) supplemented with bromophenol blue were denaturated at 95°C for 5min. Samples migrated on 7.5% (for proteins over 200kDa) or 10% SDS-PAGE polyacrylamide gel at 140V with 1xTris-Glycine SDS running buffer (Euromedex®). Proteins were transferred on nitrocellulose membrane 0.45 μm in 1xTris-Glycine (Euromedex®), 20%EtOH transfer buffer at 100 V for 1h. Membrane were blocked with 5% milk, 1X PBS-T (Phosphate Buffer Saline) (Dominique Dutscher®), 2% Tween® 20 for 1h. Primary antibodies **(Table S1)** were used at 1/2000^e^ in 5% milk, 1X TBS, 0.1% Tween 20 overnight. Secondary antibodies were used at 1/5000^e^ in 1X TBS, 0.1% Tween 20 for 1h. Protein revelation was done using the Clarity or Clarity Max ECL Western Blotting Substrates (Biorad®) and the Chemidoc MP imaging system (Biorad®).

For co-immunoprecipitation, leaf tissue from transgenic lines were collected, frozen in liquid nitrogen and ground to powder. Proteins fused to mCITRINE were immunoprecipitated using the pull-down kit Miltenyi μMacs anti-GFP (Miltenyi Biotec®). Crude extracts were put in 1XSDG supplemented with bromophenol blue and denaturated at 95°C for 5 min while IP samples were directly ready for migration on polyacrylamide gels.

Each co-immunoprecipitation experiment was repeated a minimum of three times independently with similar outcomes.

### Mass spectrometry analysis

Mass spectrometry analysis had been performed by the Protein Science Facility of SFR (Structure Fédérative de Recherche) Bioscience Lyon. Trypsine-digested samples were analysed by LC-MS/MS (Orbitrap, ThermoFischer Scientific®). The peptides identified were compare to the UniProtKb database.

### Yeast-two-hybrid

The initial screen was performed by hybrigenics services (https://www.hybrigenics-services.com/contents/our-services/interaction-discovery/ultimate-y2h-2), using the ULTImate Y2H screen against their Universal Arabidopsis Normalized library obtained using oligo_dT. The residues 2 to 1479 of PI4K◻1 were used. The screen was performed on 0.5 mM 3AT, 58.6 million interactions were analyzed, and 313 positive clones were sequenced.

AD vectors (prey) and DB vectors (bait) were transformed in the Y8800 and Y8930 yeast strains, respectively. Transformed yeasts were plated on SD (0.67% yeast nitrogen base, 0.5% dextrose anhydrous, 0.01% adenine, 1.8% agar) with all amino acids except tryptophan (SD-T) for AD vectors and SD-L for DB vectors for selection.

Yeast colony transformed with AD- or DB- vectors were grown in SD-T and SD-L liquid media, 30°C, 200rpm for 48h. Mating was performed at 30°C, 200rpm, overnight using 10μl of AD and 10μl DB clones in 180μl YEPD (1% yeast extract, 2% peptone, 2% dextrose anhydrous, 0.01% adenine) liquid media. Diploid yeasts were selected by addition of 100μl of SD-LT liquid media. Yeast were grown for 48h at 30°C, 200rpm. Diploids yeasts were plates on the different selective media: SD-LT to verify the mating; SD-LTH to select the positive interactions; SD-LTH+3AT (3-Amino-1,2,4-Triazol at 1mM final) to select strong interactions only: SD-LH+CHX (cycloheximide at 10μg/mL final) to determine the autoactivated DB clones; SD-LH+CHX+3AT to determine which concentration of 3AT erase the autoactivation of DB clones. The experiment has been repeated three times independently and similar results were obtained.

### Crispr lines

20bp target sequence upstream to a 5’-NGG -3’ sequence in an exon of the gene of interest were found using the CrisPr RGEN tool (www.rgenome.net/). Primers were designed following the methods developed in (Wang et al., 2015; Xing et al., 2014) **(Table S7)**.

### T-DNA and Crispr Mutant Genotyping

Leaf tissue from wild-type plants, T-DNA insertion lines and Crispr lines were collected, frozen in liquid nitrogen and ground to powder. The powder was resuspended in the extraction buffer (200mM of Tris pH 7.5, 250mM of NaCl, 25mM of EDTA, 0.5% of SDS). DNA was precipitated with isopropanol and the DNA pellet was washed with 75% ethanol before resuspension in water. Plants were genotyped by PCR using the GoTaq® polymerase (Promega®) and the indicated primers **(Table S8)**. PCR products were migrated on 1% agarose gel or the percentage indicated **(Table S8)**. When sequencing was required, the bands were purified using the NucleoSpin Gel and PCR Clean kit (Macherey-Nagel®) and sequenced.

### Pollen observation by SEM

Pollen grains from mutant or WT flowers were placed on tape and observed using the mini SEM Hirox® 3000 at −10°C, 10kV.

### Pollen inclusion and observation by TEM

For transmission electron microscopy, anthers were placed in a fixative solution of 3.7% paraformaldehyde and 2.5% glutaraldehyde, Na_2_HPO_4_ 0.1 M, NaH_2_PO_4_ 0.1 M overnight and postfixed in 1% OsO_4_, Na2HPO4 0.1 M, NaH_2_PO_4_ 0.1 M. Anthers were dehydrated through a graded ethanol series from 30% to 100% and embedded in SPURR resin. Sections were made using an ultramicrotome Leica UC7 at 70-80nm and poststained with Acetate Uranyle 5% (in Ethanol), Lead Citrate (in NaOH). Pollen were observed using a transmission electron microscope Jeol 1400 Flash. For the following genotype: *pi4kα1-1, npg1-2+/− npgr1−/− npgr2-2−/−, hyc1 and efop3-1−/−efop4-2−/−*, shrivelled pollen grains were selected for imaging and compared with wild-type pollen.

### Pollen staining

To perform Alexander staining, flowers that were about to open were dissected and anthers were put between slide and coverslip in Alexander staining solution (25% glycerol, 10% EtOH, 4% acetic acid, 0.05% acid fuchsin, 0.01% Malachite green, 0.005% phenol, 0.005% chloral hydrate, 0.005% Orange G,) for 7h at 50°C before observation under a stereomicroscope.

For DAPI staining, flowers that were about to open were dissected and their anthers were put between slide and coverslip in DAPI solution (10% DMSO, 0.1% NP-40, 50mM PIPES, 5mM EGTA, 0.01% (4’,6-diamidino-2-phénylindole) DAPI) for 5min at RT before observation at the confocal microscope. DAPI was excited with a 405nm laser (80mW) and fluorescence emission was filtered by a 447/60 nm BrightLine® single-band bandpass filter (Semrock, http://www.semrock.com/).

### Seed clearing

Siliques were opened. The replums with the seeds attached were placed between slide and coverslip in clearing solution (87.5% chloral hydrate, 12.5% glycerol) for 1 hour before observation at the microscope with differential interference contrast.

### Artificial microRNA phenotype

Surface-sterilized seeds were plated on Murashige and Skoog (MS, Duchefa BiochemieR) media supplemented with sucrose (10g per L) containing 5μM β-estradiol or the equivalent volume of DMSO. Plates were vernalized 3 days before being placed under continuous white light conditions for 11 days. Plates were scanned. Root length was measured using ImageJ software. Similar result was obtained for five independent experiments.

### Confocal Imaging setup

7 to 10 days-old seedlings were observed with an Zeiss microscope (AxioObserver Z1, Carl Zeiss Group, http://www.zeiss.com/) with a spinning disk module (CSU-W1-T3, Yokogawa, www.yokogawa.com) and a Prime 95B camera (Photometrics, https://www.photometrics.com/) using a 63x Plan-Apochromat objective (numerical aperture 1.4, oil immersion) and the appropriated laser and bandpath filter.

GFP was excited with a 488 nm laser (150mW) and fluorescence emission was filtered by a 525/50 nm BrightLine® single-band bandpass filter (Semrock, http://www.semrock.com/). mCITRINE was excited with a 515nm laser (60mW) and fluorescence emission was filtered by a 578/105 nm BrightLine® single-band bandpass filter (Semrock, http://www.semrock.com/). mCHERRY was excited with a 561nm laser (80mW) and fluorescence emission was filtered by a 609/54 nm BrightLine® single-band bandpass filter (Semrock, http://www.semrock.com/). In the case of seedling expressing mCHERRY and mCITRINE markers, mCITRINE was excited and emission was filtered using the GFP settings.

For each line, several independent transformation events were used and observed in three independent experiment at minima. TIRF microscopy used an objective based azimuthal ilas2 TIRF microscope (Roper Scientific) with 100x Apo NA 1.46 Oil objective. Exposure time used was 1s. HYC2-mCITRINE and NPGR2-mCITRINE were excited and emission was filtered using the GFP settings while EFOP2-mCITRINE was excited and emission was filtered using the YFP settings.

### Immunolocalization

Six-days old seedlings were fixed in PFA 4%, MTBS (50mM PIPES, 5mM EGTA, 5mM MgS04, pH7) for 2h. Roots were Superfrost Plus® slides (Thermo Scientific, #10149870) and dried at RT for 1h and rehydrated using MTBS + 0.1% tritonx100. Permeabilization was done using enzymatic digestion (250μl of 0.75% pectinase, 0.25% pectolyase, 0.5% macérozyme, 0.5% cellulase stock solution was diluted into 1mL MTBS) for 40 min (Rozier et al., 2014). Roots were treated with 10% dimethylsulfoxide, 3% Igepal (Sigma, #CA-630) in MTSB for 1 h. Blocking was done using 5% normal goat serum (NGS Sigma #G9023) in MTSB. Roots were incubating with the primary antibody diluted at 1/100 (anti-PI4Kα1) and with secondary antibody diluted at 1/300 (anti Rat IgG, alexa Fluor 488 conjugate, InVitrogen-Molecular Probes, #A-21210) overnight at 4°C. Roots were placed between slide and coverslip in Vectashield® (Vectorlabs, #H-1000-10) and observed using a confocal microscope Zeiss LSM800. Similar results were obtained in six independent experiments.

### Biochemical quantification of PIP levels in inducible PI4K knockdown

Nine-days-old seedlings grown on 1/2 MS plates with 1% (w/v) sucrose, 1% (w/v) agar and 5μM β-estradiol or DMSO (control) were transferred to 2 ml Safe-lock Eppendorf tubes containing 190 μL labelling buffer (2.5 mM MES (2-[N-Morpholino]ethane sulfonic acid) (pH 5.7 with KOH), 1 mM KCl) supplemented with either 5μM β-estradiol or DMSO (control), with each tube containing three seedlings. Seedlings were metabolically labelled with radioactive phosphate by incubating them overnight for ~16-20 hrs with 10 μL (5-10 μCi) carrier-free ^32^P-PO_4_^3-^ (^32^P_i_; PerkinElmer, The Netherlands) in labelling buffer. Incubations were stopped by adding 50 μL of 50% (w/v) perchloric acid and the lipids extracted (Zarza et al., 2020). PIP was separated from the rest of the phospholipids by thin-layer chromatography (TLC) using K-oxalate-impregnated and heat-activated silica gel 60 plates (20×20 cm; Merck) and an alkaline TLC solvent, containing chloroform/methanol/25% ammonia/water (90:70:4:16) (Munnik and Zarza, 2013). Each lane contained 1/5^th^ of the extract. Radioactivity was visualized by autoradiography and quantified by phosphoimaging (Typhoon FLA 7000, GE Healthcare). PIP and PA levels were quantified as percentage of total ^32^P-labelled phospholipids. Experiments were performed in quadruplicate.

### Microsomes and plasma membrane purification

Microsomes were purified as described in (Simon-Plas et al., 2002) and resuspended in phosphate buffer (5 mM, pH 7,8) supplemented with sucrose (300 mM) and KCl (3 mM). Plasma membrane were obtained after cell fractionation by partitioning twice in a buffered polymer two-phase system with polyethylene glycol 3350/dextran T-500 (6.6% each).

For PI4Kα1, proteins were precipitated using 5 volumes of −20°C acetone for 1 volume of protein extraction and incubated for 10min at −20°C. Proteins were pelleted. This process was repeated 2 more times. Pellet was dried and resuspended in 1xSDG supplemented with bromophenol blue. All steps were performed at 4 °C.

### Plasmolysis

7 to 10 days-old seedlings were placed between side and coverslip in MES 10 mM at pH 5.8 with or without 0.75M sorbitol and observed using a confocal microscope.

### Fluorescence recovery after photobleaching (FRAP)

Fluorescence in a rectangle ROI (5×1,7μm) at the plasma membrane was bleached in the root by successive scans at full laser power (150 W) using the iLas2 FRAP module (Roper scientific, http://www.biovis.com/ilas.htm). Fluorescence recovery was measured in the ROIs and in controlled ROIs (rectangle with the same dimension in unbleached area). FRAP was recorded continuously during 120 s with a delay of 1s between frames. Fluorescence intensity data were normalized as previously described (Martinière et al., 2012). The mobile fraction was calculated at t=120s with the following formula: I(t)-Min(I)/Ictrl(t)/Min(I) where I(t) and Ictrl(t) are the intensity of the bleached and control region at a time t, repectively. Data were obtained in several independent experiments.

### Nicotiana benthamiana leaf infiltration

Transformed agrobacterium were directly taken from plate with a tip and resuspended into 2 mL of infiltration media (10 mM MES pH 5.7, 10 mM MgCl_2_, 0.15 mM acetosyringone (Sigma-Aldrich®, #D134406)) by pipetting. The OD_600_ was measured using a spectrophotometer (Biophotometer, Eppendorf) and adjusted to 1 by adding infiltration media.

The infiltration was performed on « heart shape » tobacco leaves from 2-3 weeks old plants. Using 1 mL syringe (Terumo®, #125162229), the infiltration solution with the agrobacterium was pressed onto the abaxial side of the chosen tobacco leaf. The plants were put back to the growth chamber for 2-3 days under long day conditions.

5 mm^2^ regions of the leaf that surround the place where the infiltration has been made were cut. The pieces of leaf were mounted in water between slide and coverslip with the abaxial side of the leaf facing the coverslip. Using the appropriate wavelength, an epifluorescent microscope and the smallest objective (10X), the surface of the leaf was screened to find the transformed cells. Then, the subcellular localization of the fluorescent protein was observed using a spinning confocal microscope and 63X objective.

### PAO treatment

Seedlings (7 to 10-days-old) were incubated in liquid MS with 30μM PAO (Sigma, www.sigmaaldrich.com, PAO stock solution at 60 mM in DMSO) for 30 min before observation using a confocal microscope.

### Statistical analysis

Statistical analyses were performed using R (v. 3.6.1, (R Core Team, 2019) and the R studio interface **(Table S10)**.

---

**Supplemental Figure 1.**
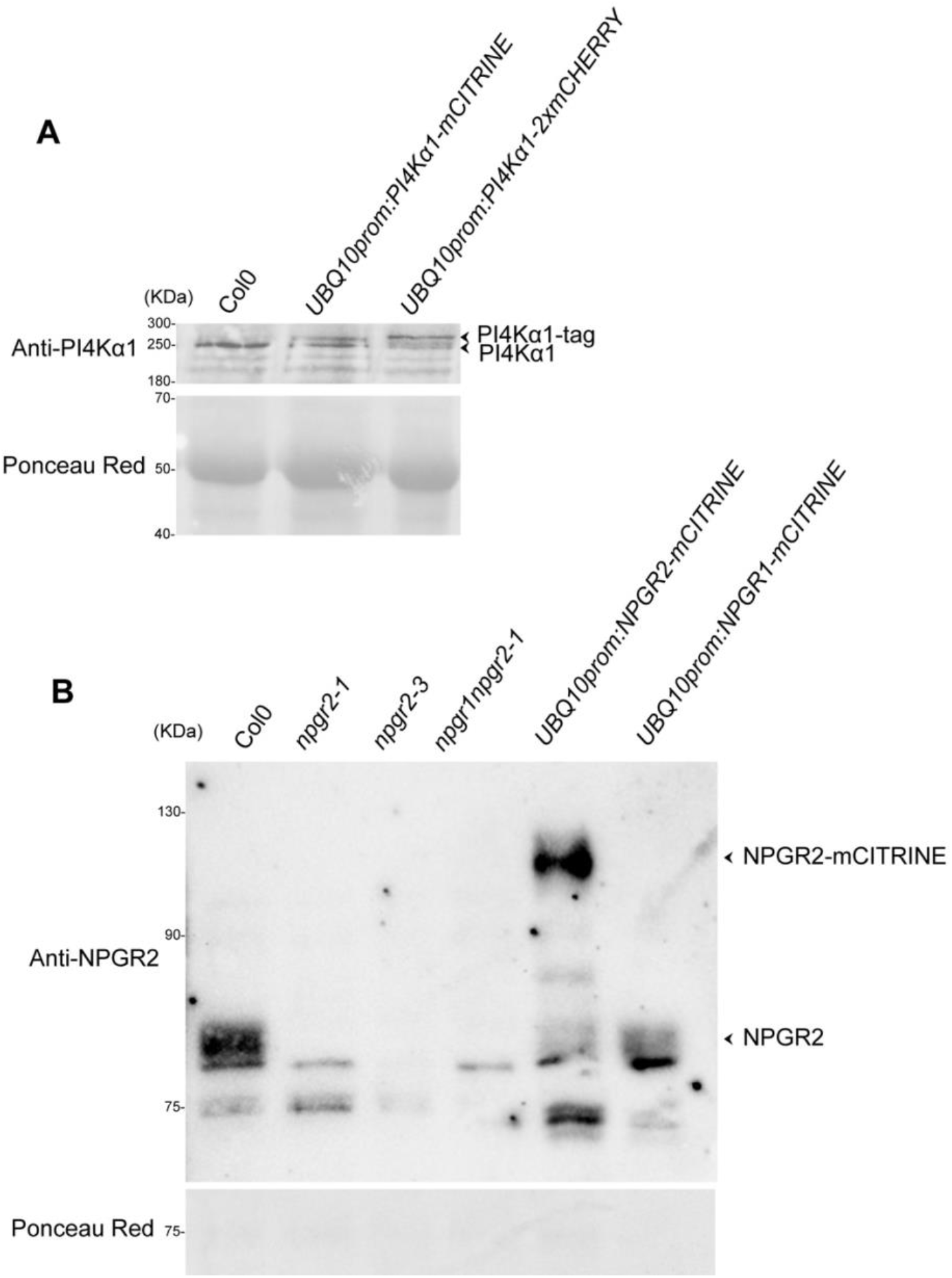
Western-blot validation of the custom-made anti-PI4Kα1 and anti-NPGR2 antibodies. **(A)** Western blot anti-PI4Kα1 on total proteins extract of Col-0 seedlings, seedling expressing *UBQ10prom:PI4Kα1-mCITRINE* and seedling expressing *UBQ10prom:PI4Kα1-2xmCHERRY*. Red ponceau shows similar protein loading in every well. **(B)** Western blot anti-NPGR2 on total proteins extract of Col0, *npgr2-1, npgr2-3, npgr1npgr2-1* seedlings, seedling expressing *UBQ10prom:NPGR2-mCITRINE* and seedling expressing *UBQ10prom:NPGR1-mCITRINE*. Ponceau red shows similar protein loading in every wells. Supports Figure 1.

**Supplemental Figure 2.**
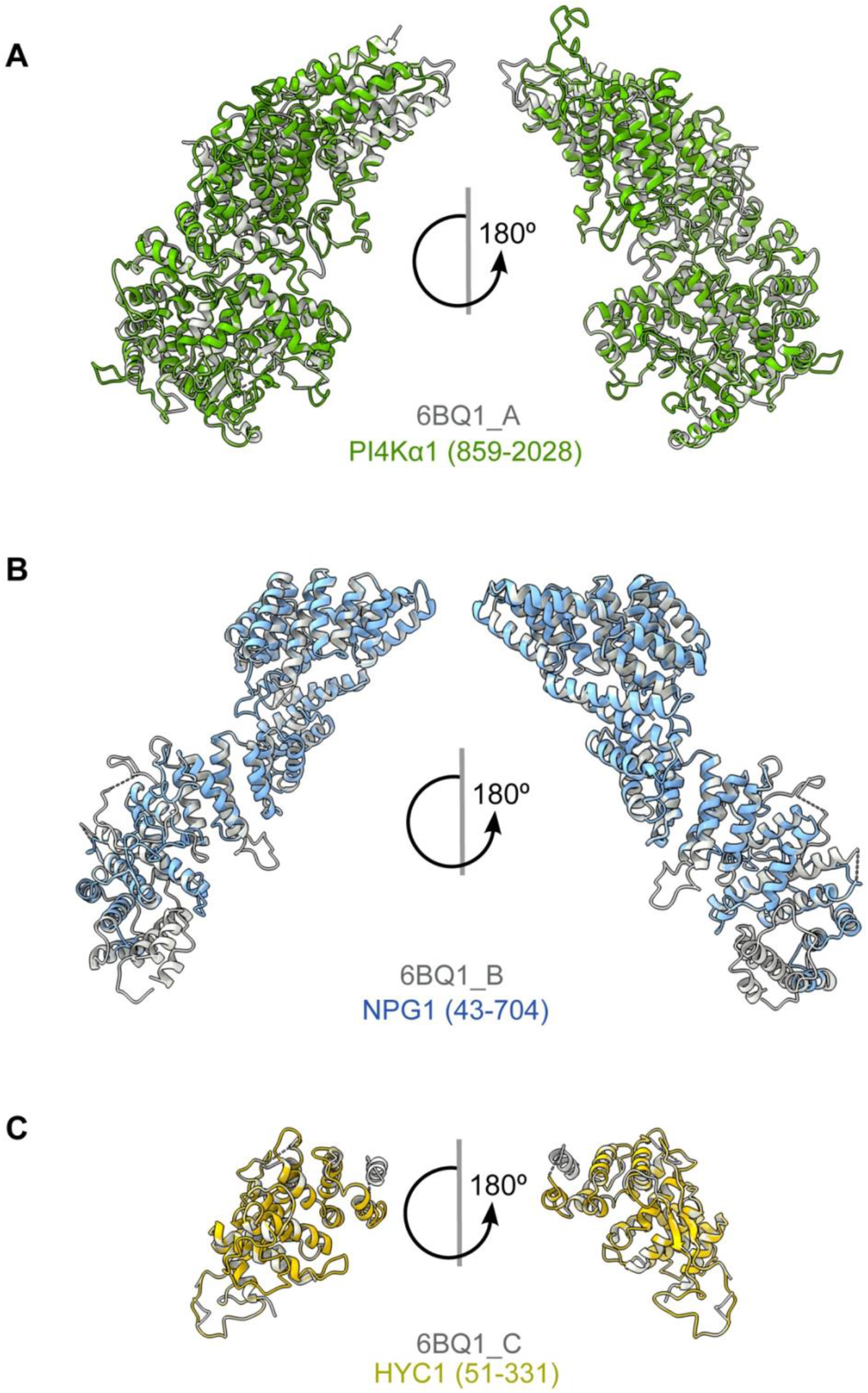
Template-based modelling of PI4Kα1, NPG1 and HYC1. **(A)** Superimposition of the template-based structure of Arabidopsis PI4Kα1 (green, amino acid region 859-2028) with the template structure of human PI4KIIIα (PDB code 6BQ1, chain A, in grey). **(B)** Superimposition of the template-based structure of Arabidopsis NPG1 (blue, amino acid region 43-704) with the template structure of human TTC7 (PDB code 6BQ1, chain B, in grey). **(C)** Superimposition of the template-based structure of Arabidopsis HYC1 (yellow, amino acid region 51-331) with the template structure of human FAM126A (PDB code 6BQ1, chain C, in grey). Supports Figure 2.

**Supplemental Figure 3.**
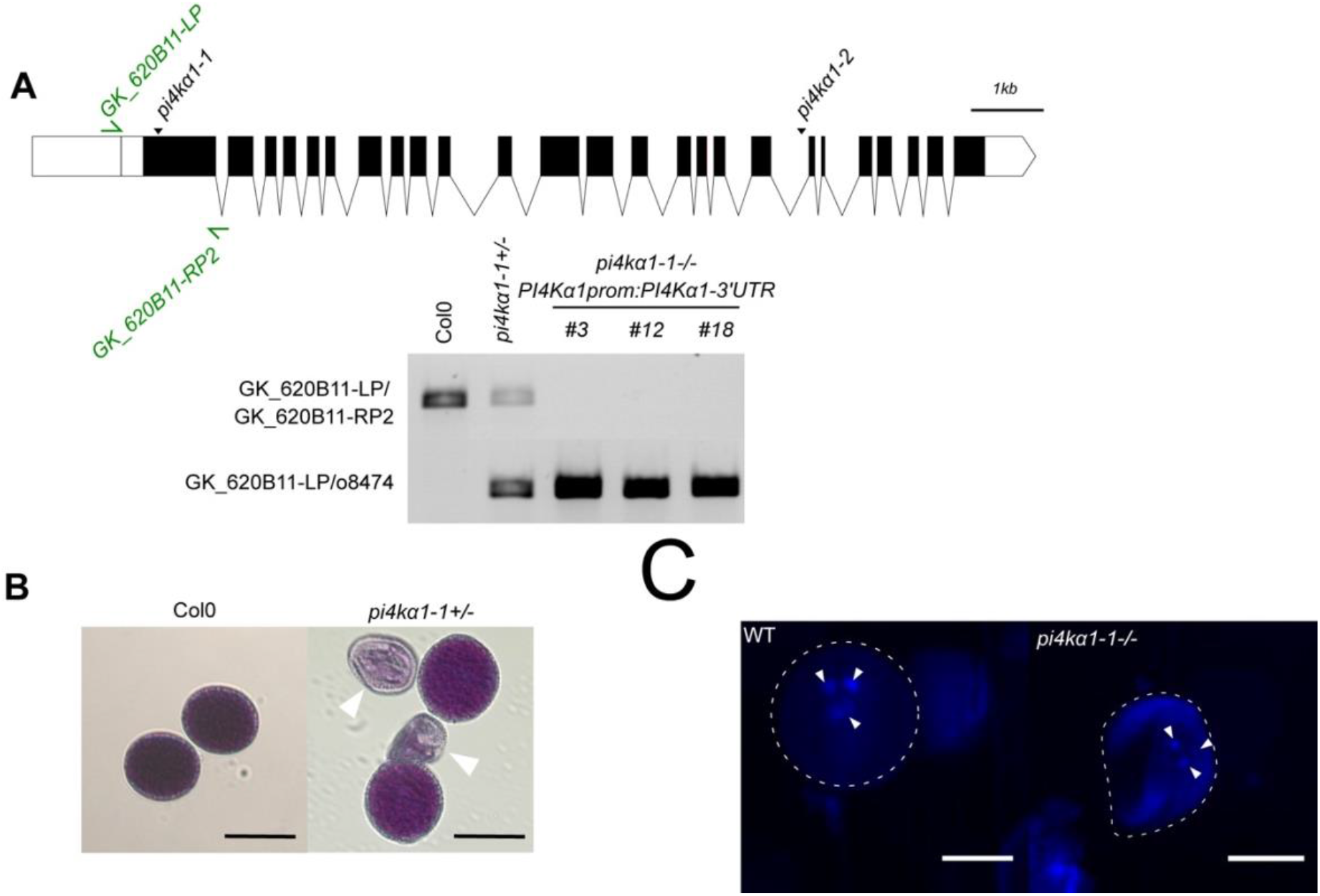
Characterization of *pi4kα1* pollen phenotype. **(A)** Genotyping of Col-0, *pi4kα1-1* heterozygous plants, and *pi4kα1-1* homozygous plants expressing *PI4Kα1prom:PI4Kα1-3′UTR* (insertion n°3, 12 and 18). Upper panel shows the amplification of gene sequence. Lower panel shows amplification of T-DNA border. **(B)** Alexander staining of pollen grains from Col-0 and self-fertilized *pi4kα1-1* heterozygous plants. Shrivelled pollen grains are indicated by white arrowheads. Scale bars: 20 μm. **(C)** DAPI staining of pollen grains from self-fertilized pi4kα1-1 heterozygous plants with normal (WT, left) and shrivelled (*pi4kα1-1 −/−*, right) shape. Nuclei are indicated with white arrowheads. Scale bars: 10 μm. Supports Figure 3.

**Supplemental Figure 4.**
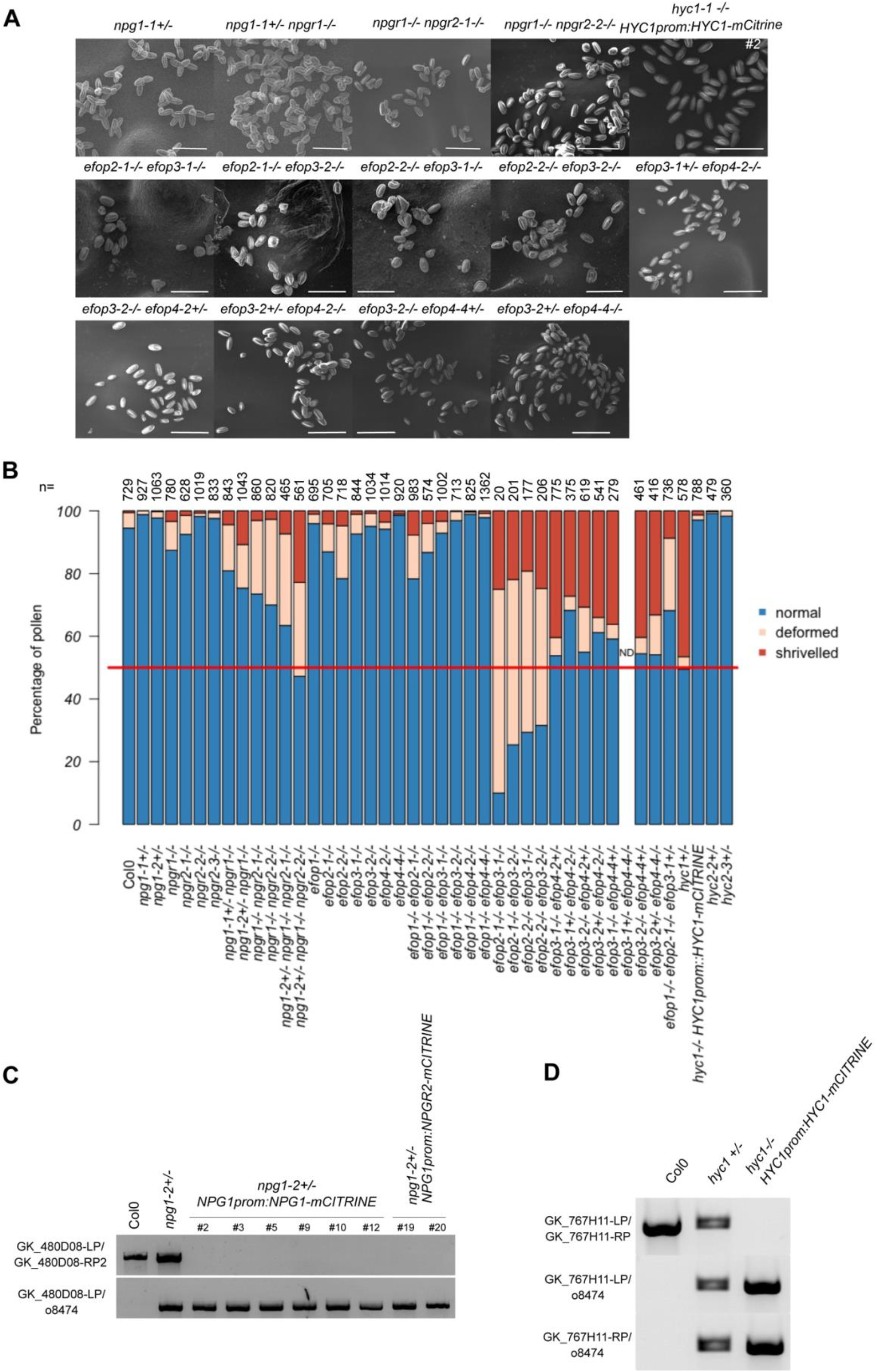
Characterization of the pollen phenotype of single and multiple *npg*, *hyc* and *efop* mutants. **(A)** Pollen grains observed at scanning electronic microscope of self-fertilized *npg1-1+/−, npg1-1+/− npgr1−/−, npgr1-1−/− npgr2-1−/−, npgr1-1−/− npgr2-2−/−, hyc1−/−* expressing *HYC1prom:HYC1-mCITRINE, efop2-1−/− efop3-1−/−, efop2-1−/− efop3-2−/−, efop2-2−/− efop3-1−/−, efop2-2−/− efop3-2−/−, efop3-1+/− efop4-2−/−, efop3-2−/− efop4-2+/−, efop3-2+/− efop4-2−/−, efop3-2−/− efop4-4+/−,* and *efop3-2+/− efop4-4−/−* plants. Scale bars: 50 μm **(B)** Quantification of the % of normal (blue), deformed (orange) and shrivelled (red) pollen grains of all indicated genotypes. ND is indicated when no quantification of this genotype is available. n indicates the number of pollen grains counted. **(C)** Genotyping of Col-0, *npg1-2* heterozygous plants and *npg1-2* homozygous plants complemented with *NPG1prom:NPG1-mCITRINE* (insertion n° 2, 3, 5, 9, 10 and 12) and *NPG1prom:NPGR2-mCITRINE* (insertion n° 19 and 20). Upper panel shows amplification of the gene sequence. Lower panel shows amplification of T-DNA border. **(D)** Genotyping of Col-0, *hyc1* heterozygous plants and *hyc1* homozygous plants complemented with *HYC1prom:HYC1-mCITRINE* (insertion n° 2). Upper panel shows amplification of gene sequence. Lower panel shows amplification of T-DNA border. Supports Figure 4.

**Supplemental Figure 5.**
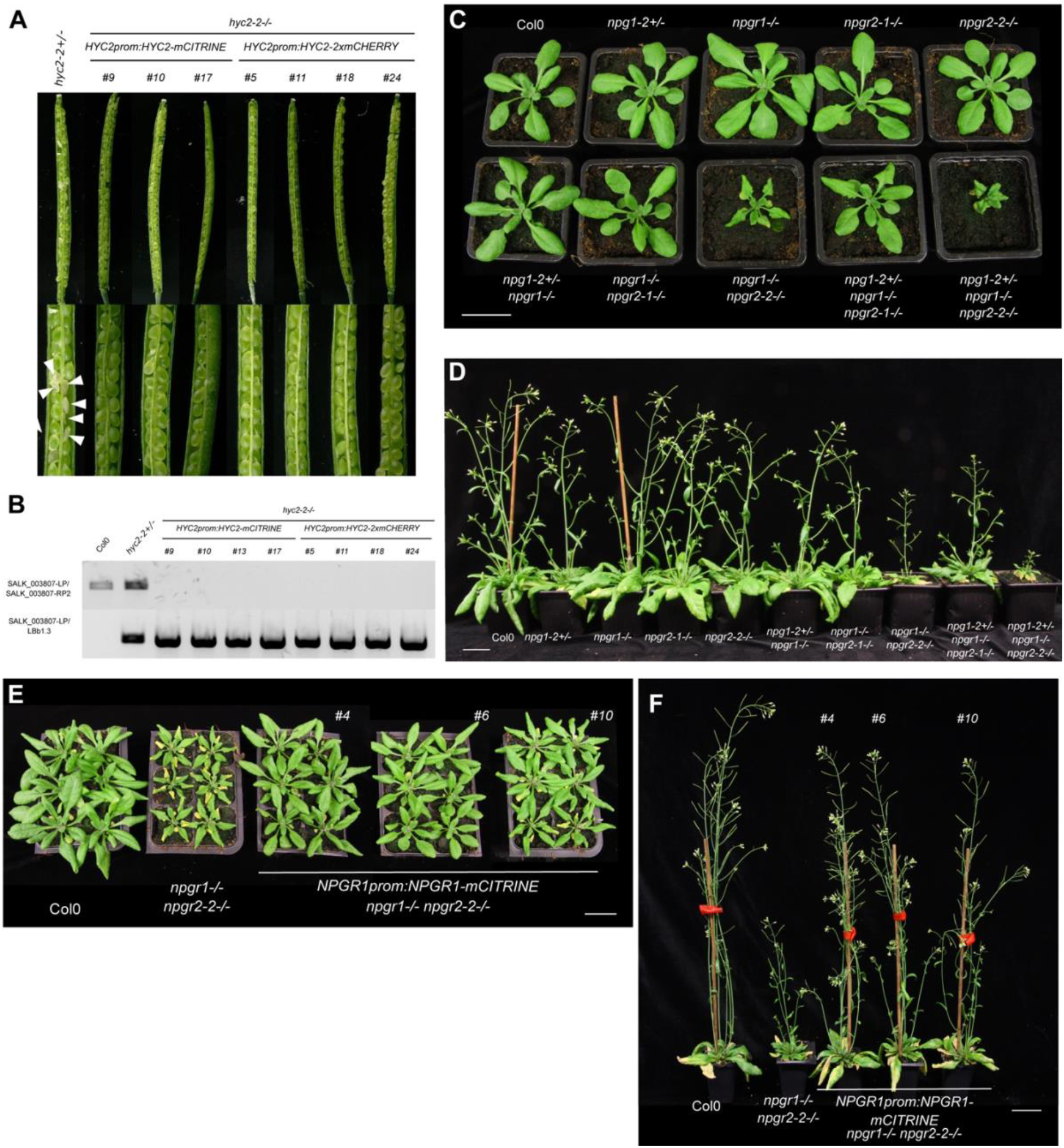
Sporophytic phenotypes and complementation of *npg* multiple mutants and *hyc2* single mutant. **(A)** Opened siliques of self-fertilized *hyc2-2* heterozygous mutant plants and self-fertilized hyc2-2 homozygous plants complemented by the expression of *HYC2prom:HYC2-mCITRINE* (insertion n°9, 10 and 17) and *HYC2prom:HYC2-2xmCHERRY* (insertion n°11, 18 and 24). White arrowheads indicate aborted seeds. **(B)** Genotyping of Col-0, *hyc2-2* heterozygous plants and *hyc2-2* homozygous plants complemented with *HYC2prom:HYC2-mCITRINE* (insertion n°9, 10, 13 and 17) and *HYC2prom:HYC2-2xmCHERRY* (insertion n° 11, 18 and 24). Upper panel shows amplification of gene sequence. Lower panel shows amplification of T-DNA border. **(C)** Twenty-seven-days-old Col-0, *npg1-2+/−, npgr1−/−, npgr2-2−/−, npg1-2+/− npgr1−/−, npgr1−/− npgr2-2−/−* and *npg1-2+/− npgr1−/− npgr2-2−/−* plants. Scale bar: 2 cm **(D)** Forty-one-days-old Col-0, *npg1-2+/−, npgr1−/−, npgr2-2−/−, npg1-2+/− npgr1−/−, npgr1−/− npgr2-2−/−* and *npg1-2+/− npgr1−/− npgr2-2−/−* plants. Scale bar: 2 cm **(E)** Twenty-seven days-old Col-0, *npgr1−/− npgr2-2−/−* and *npgr1−/− npgr2-2−/−* expressing NPGR1prom:NPGR1-mCITRINE. Several independent insertions are shown. Scale bar: 2 cm **(F)** Forty days-old Col-0, *npgr1−/− npgr2-2−/−* and *npgr1−/− npgr2-2−/−* expressing NPGR1prom:NPGR1-mCITRINE. Several independent insertions are shown. Scale bar: 2 cm. Supports Figure 5.

**Supplemental Figure 6.**
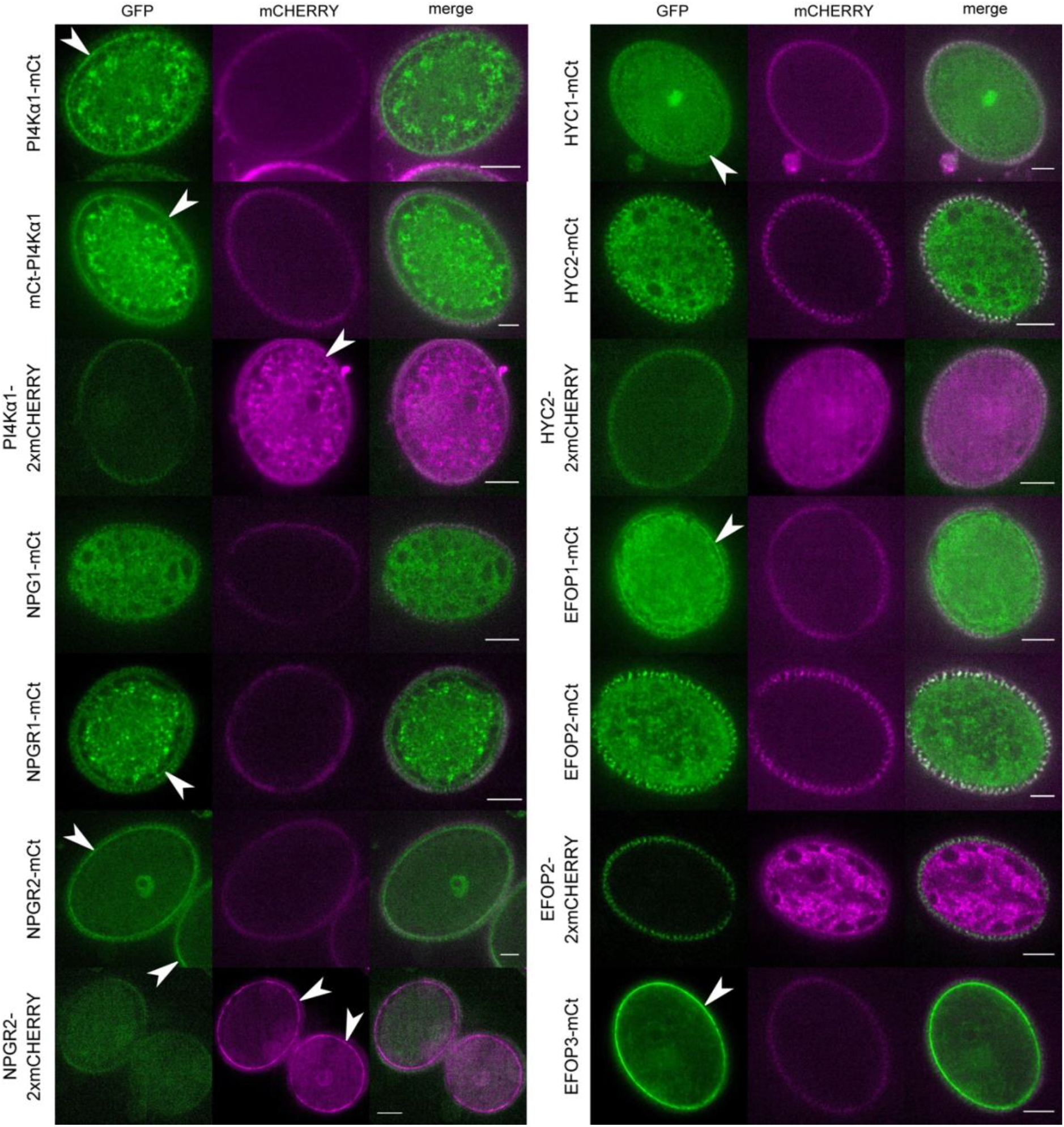
PI4Kα1, NPG, HYC and EFOP protein localization in pollen grains. Confocal images of PI4Kα1, NPG1, NPGR1, NPGR2, HYC1, HYC2, EFOP1, EFOP2 and EFOP3 fused to mCITRINE (mCt) or mCHERRY under control of *UBQ10* promoter in pollen grains. Both GFP and mCHERRY fluorescence emission channels are shown to visualise the autofluorescence from the pollen cell wall. White arrowheads indicate plasma membrane labelling. Scale bars: 10μm. Supports Figure 7.

**Supplemental Figure 7.**
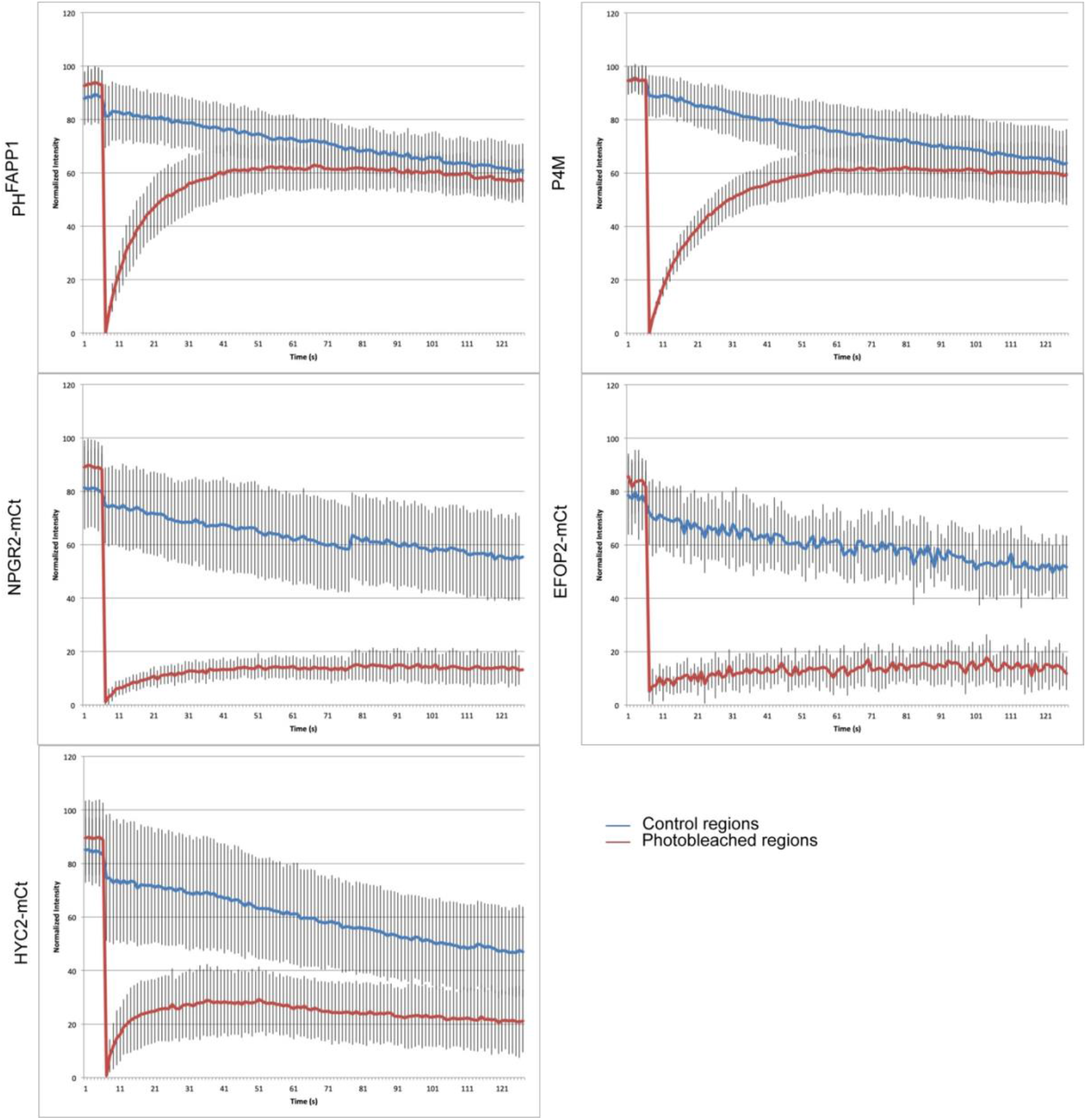
FRAP analysis of NPGR2/EFOP2/HYC2 fused with mCITRINE in Arabidopsis root. Graphics presenting signal intensity over time for photobleached (red) and control regions (blue). Standard deviations are shown. The number of zones measured is 37, 32, 30, 13 and 29 for P4M, PH^FAPP1^, NPGR2, EFOP2-mCt and HYC2-mCt, respectively. Supports Figure 8.

**Supplemental Figure 8.**
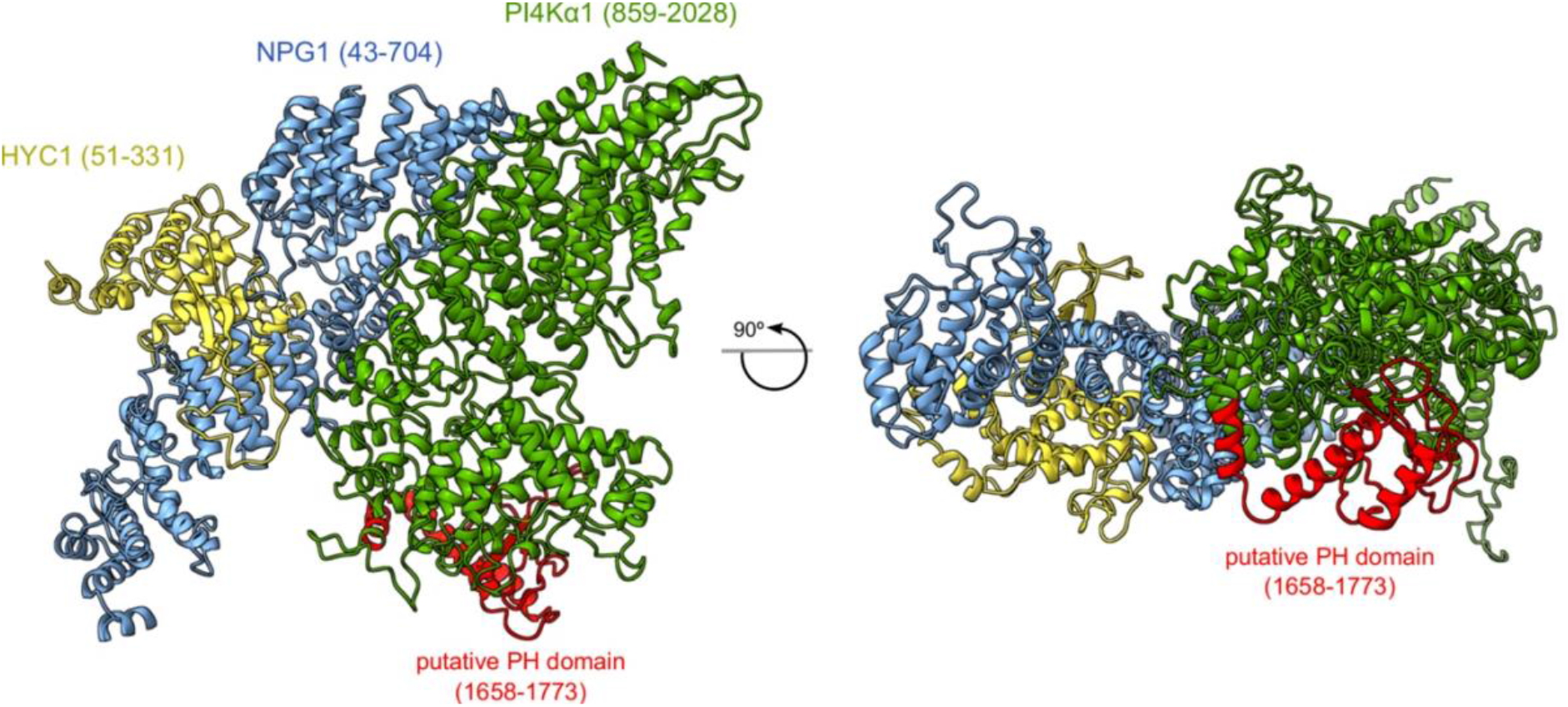
Position of the putative PH domain of PI4Kα1. Mapping of the region corresponding to the putative PH domain of PI4Kα1 described by Stevenson et al., 1998; Stevenson-Paulik et al., 2003; Xue et al., 1999. on the heterotrimeric PI4Kα1 complex structure obtained by template-based modelling and protein-protein docking. The putative PH domain overlaps with the helical (cradle) and catalytical domain of PI4Kα1. Supports Figure 8.

**Table S1:** Antibodies.

|  | **Antibody** | **References** | **Company** | **Host** | **Concentration used** |
| --- | --- | --- | --- | --- | --- |
| Primary | anti-GFP | A-6455 | ThermoFisher | Rabbit | 1/2000 |
|  | anti-mCherry | ab167453 | Abcam | Rabbit | 1/1000 |
|  | anti-PI4Ka1 | A03 rat #1 | Proteogenix | Rat | 1/1000 |
|  | anti-NPGR2 | 9094-A01 rabbit #2 | Proteogenix | Rabbit | 1/2000 |
| Secondary | anti-Rat IgG, HRP conjugate | AP136P | Merck | Goat | 1/5000 |
|  | anti-Rabbit IgG, HRP conjugate | W4011 | Promega | Goat | 1/5000 |

**Table S2:**
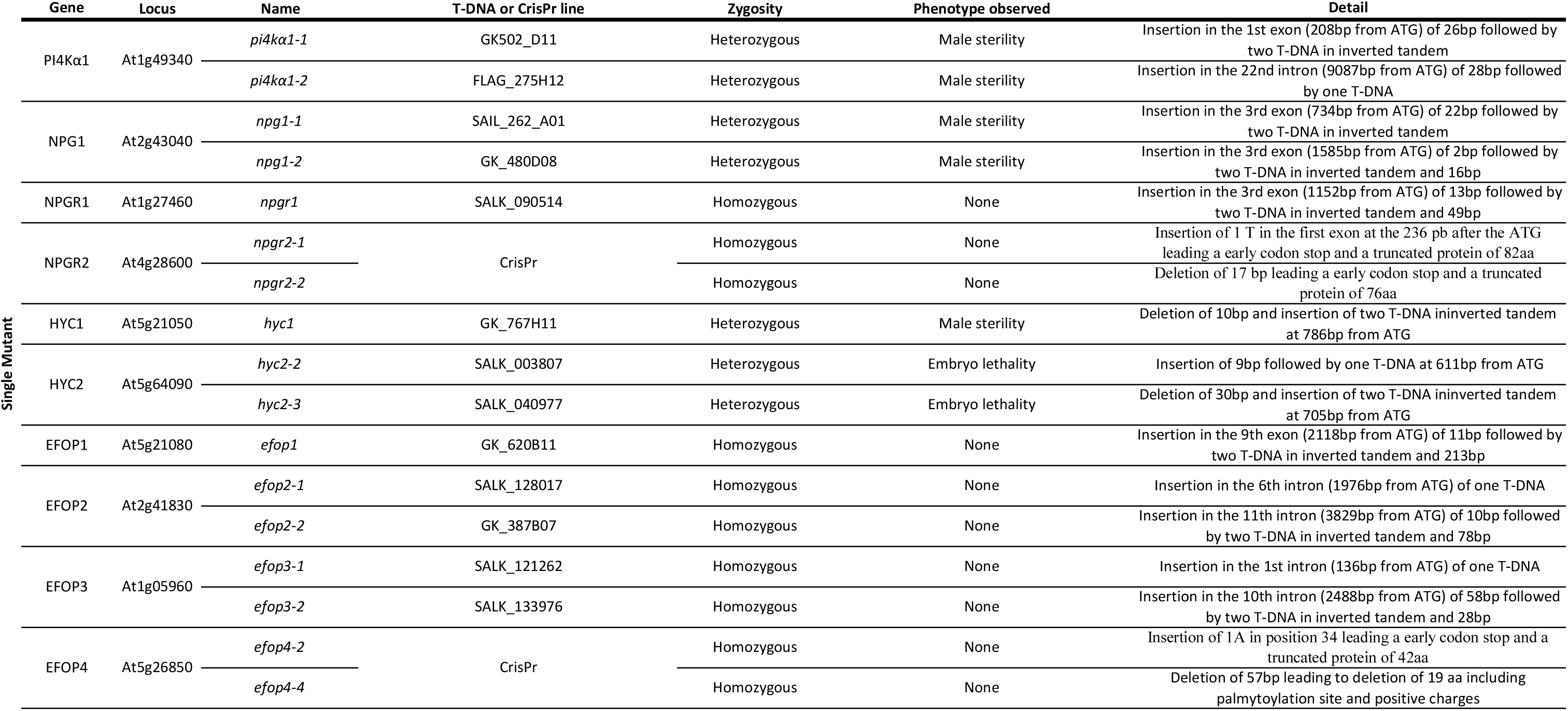
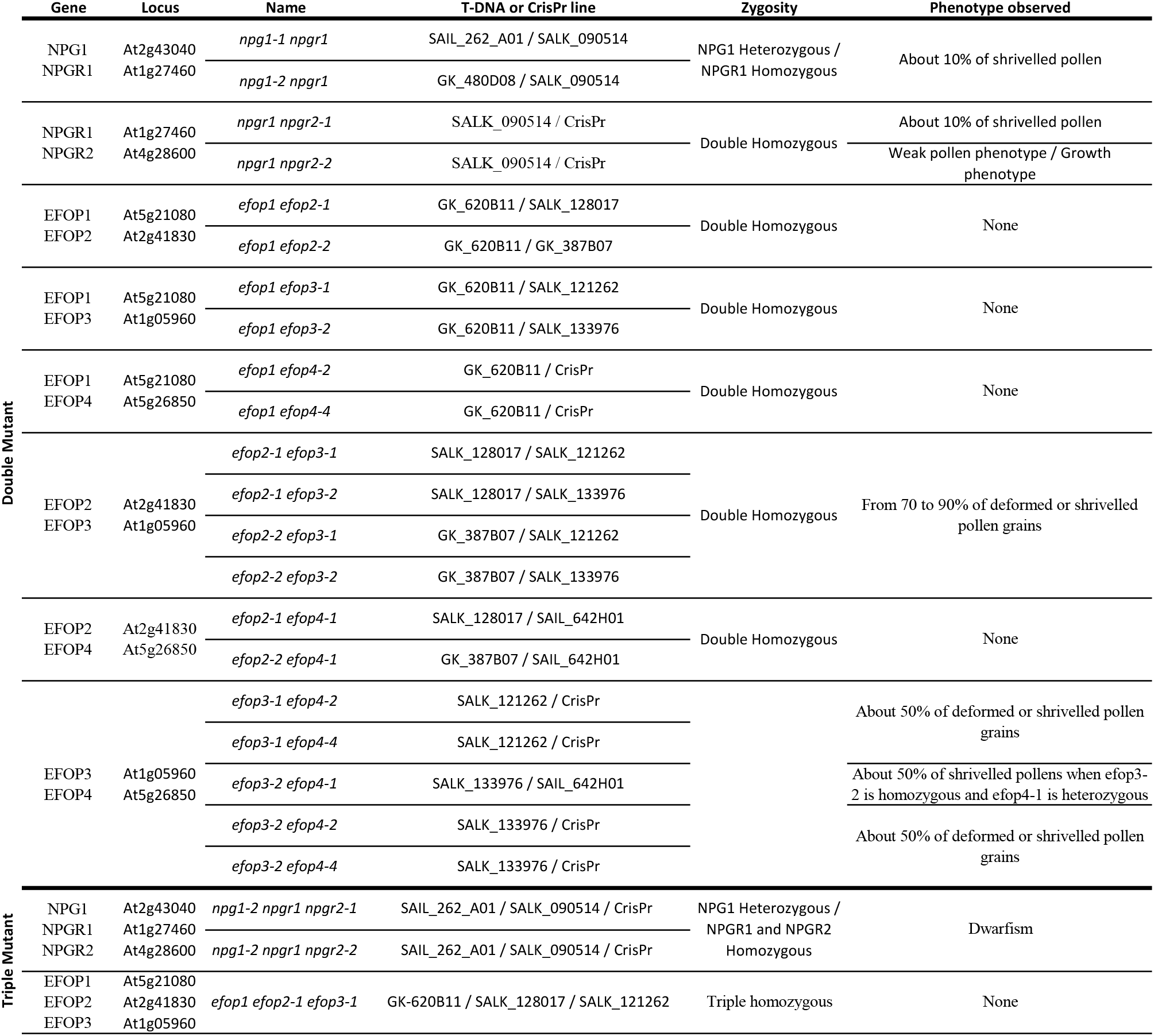
Description of the single and multiple mutants analysed in this study.

**Table S3:**
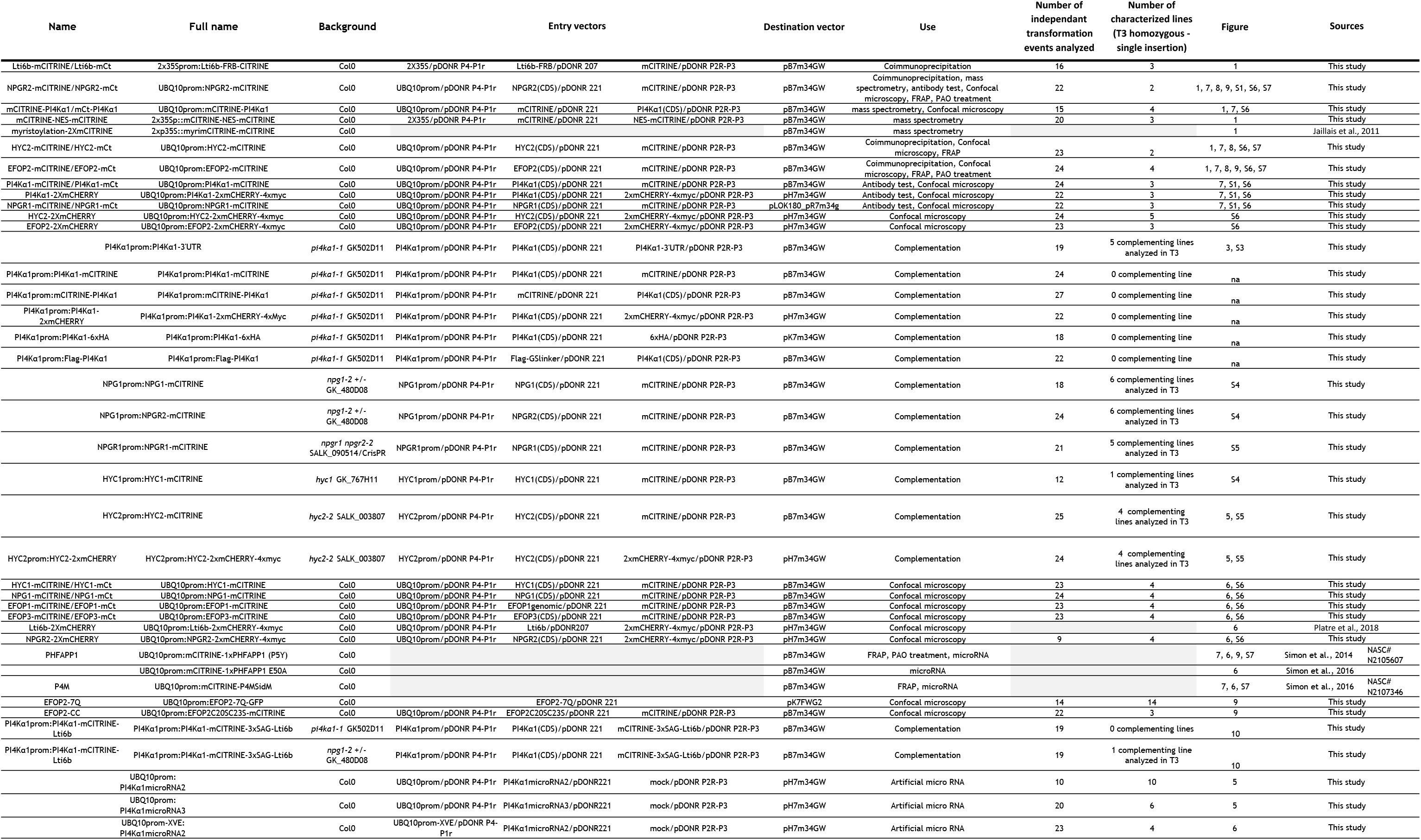
Transgenic lines.

**Table S4:** Published vectors used in this study.

| Name | Source |  |
| --- | --- | --- |
| Empty gateway entry vector:<br>pDONR221 | thermofisher | cat# 12536017 |
| Empty gateway entry vector:<br>pDONR P4P1R | thermofisher | cat# 12537023 |
| Empty gateway entry vector:<br>pDONR P2RP3 | thermofisher | cat# 12537023 |
| Empty gateway destination<br>vector: pB7m34GW | Karimi et al.,<br>2007 | N/A |
| Empty gateway destination<br>vector: pH7m34GW | Karimi et al.,<br>2007 | N/A |
| Empty gateway destination<br>vector: pLOK180_pR7m34g | L. Kalmbach and<br>M. Barberon | N/A |
| Empty gateway destination<br>vector: pK7FWG2 | Karimi et al.,<br>2002 | N/A |
| Entry vector:<br>UBQ10prom/pDONR P4-P1r | Jaillais et al.,<br>2011 | NASC N2106315 |
| Entry vector: 2X35S/pDONR<br>P4-P1r | Mar Marquès-Bueno, M. et al.<br>2016 | NASC N2106316 |
| Entry vector:<br>mCITRINE/pDONR 221 | Simon et al.,<br>2014 | NASC N2106287 |
| Entry vector:<br>mCITRINE/pDONR P2R-P3 | Jaillais et al.,<br>2011 | NASC N2106288 |
| Entry vector: 6xHA/pDONR<br>P2R-P3 | Jaillais et al.,<br>2011 | N/A |
| Entry vector: 2xmCHERRY-4xmyc/pDONR P2R-P3 | Simon et al.,<br>2014 | NASC N2106292 |
| Lti6b/pDONR207 | Elsayad et al.,<br>2016 | N/A |
| pEGFP-FRB (Gift of Klaus<br>Hahn) | Gift of Klaus<br>Hahn | plasmid addgene<br>#25919 |
| UBQ10prom-XVE/pDONR P4-P1r | Siligato et al.,<br>2016 | N/A |

**Table S5:**
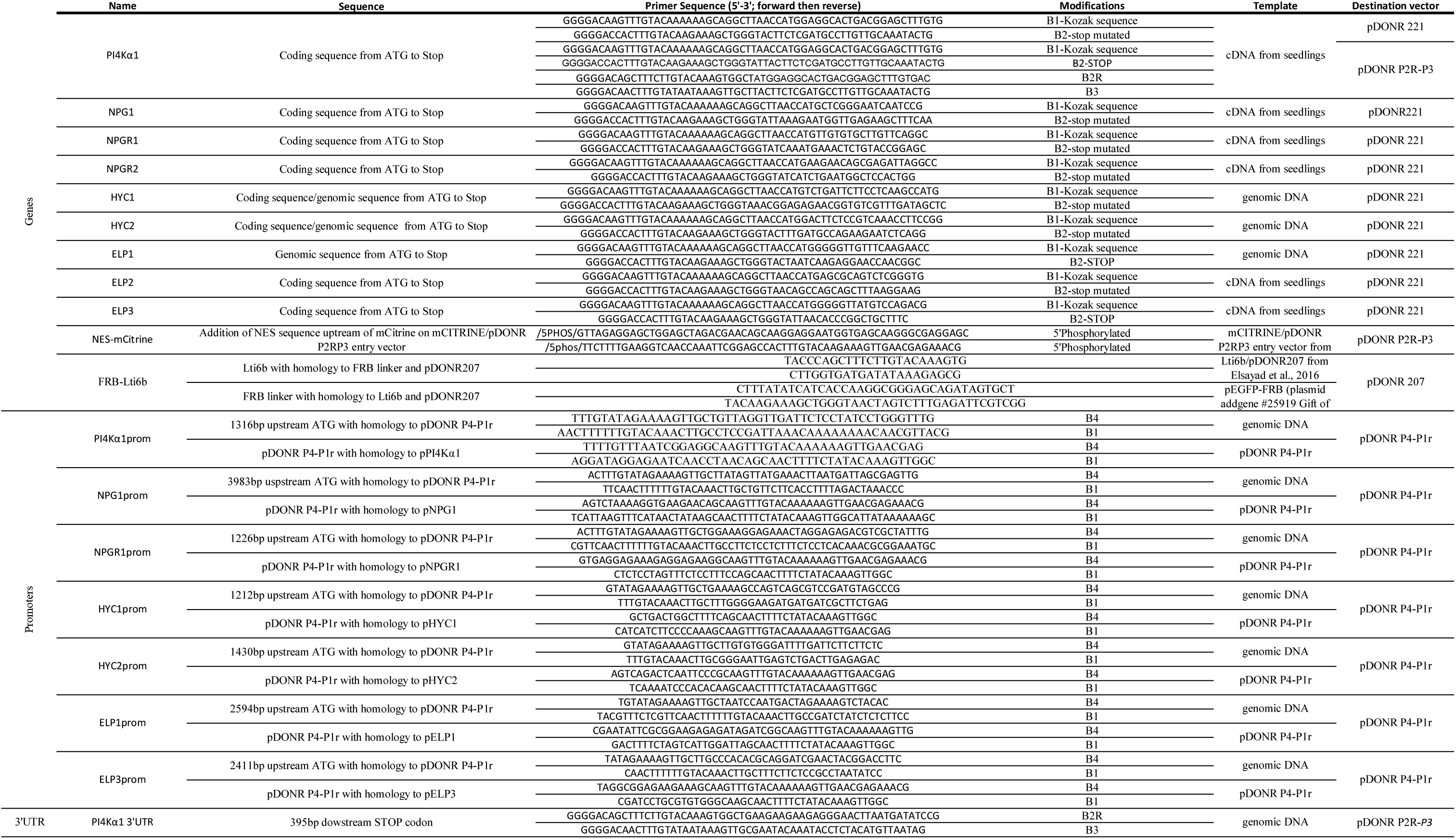
Primers used for cloning into gateway entry vectors.

**Table S6:**
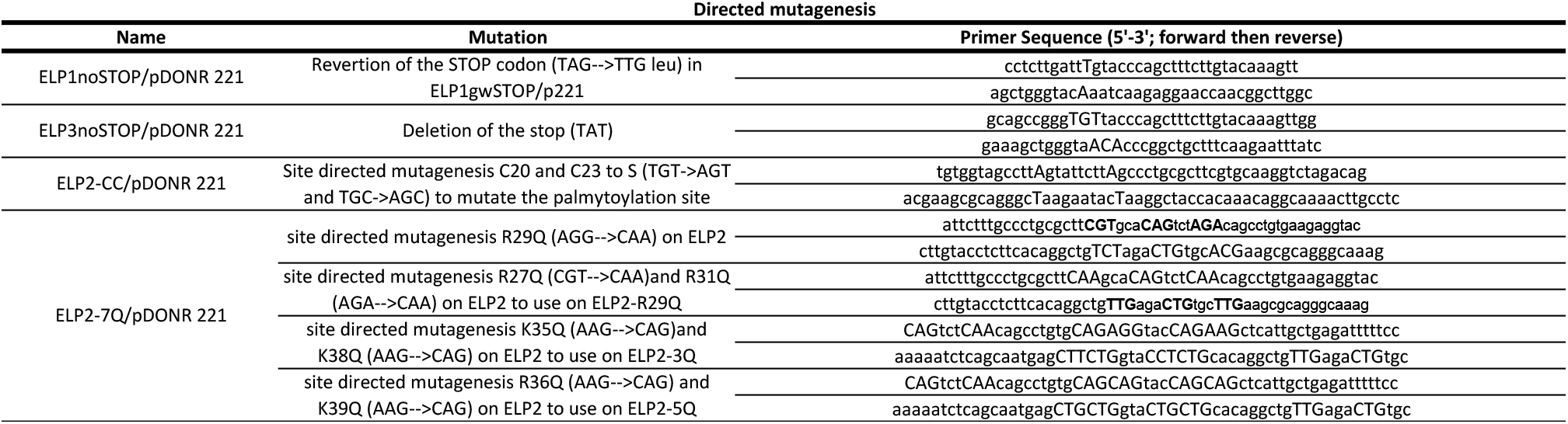
Primers used for site directed mutagenesis.

**Table S7:**
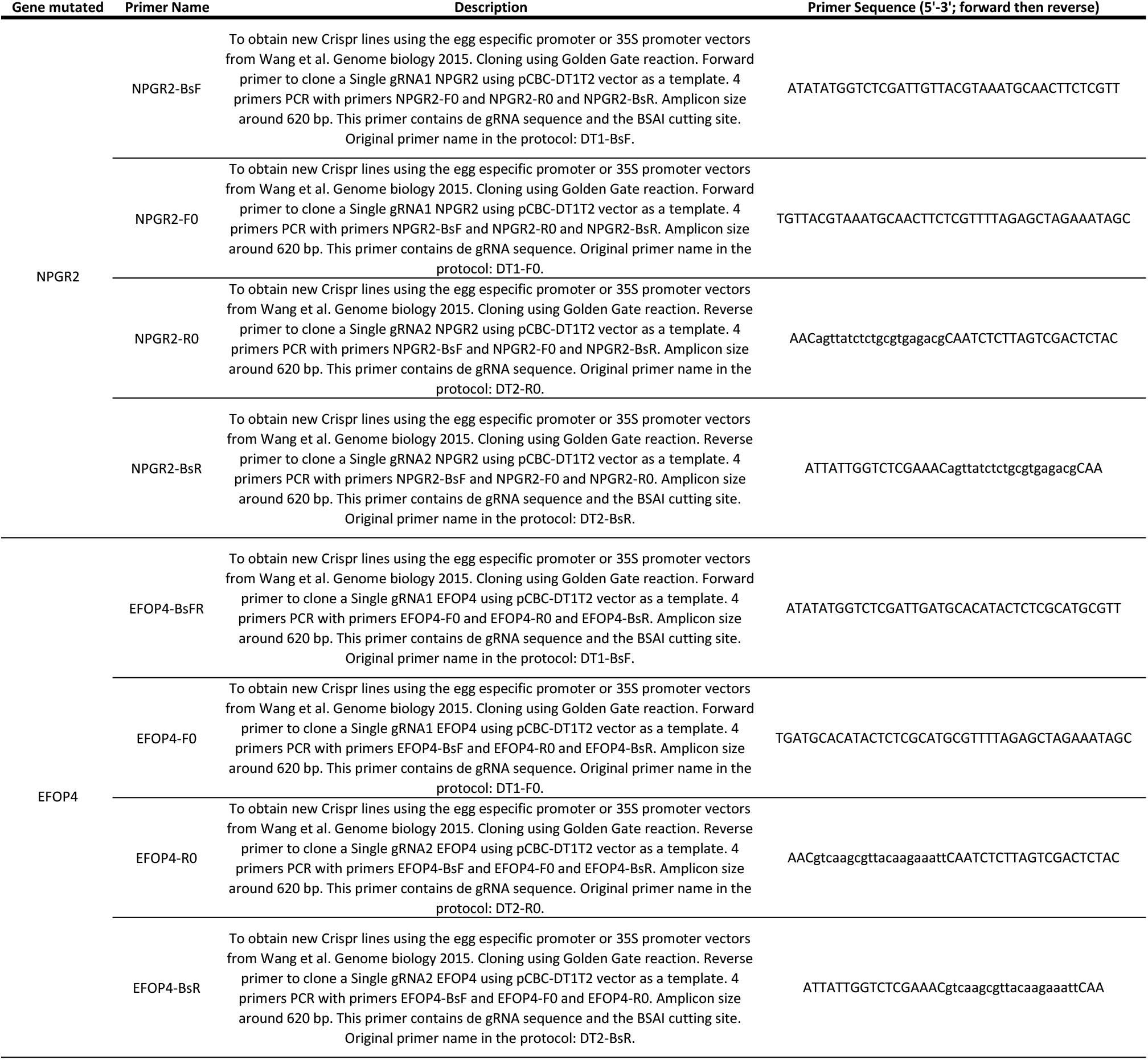
Primers used for crispr constructs.

**Table S8:**
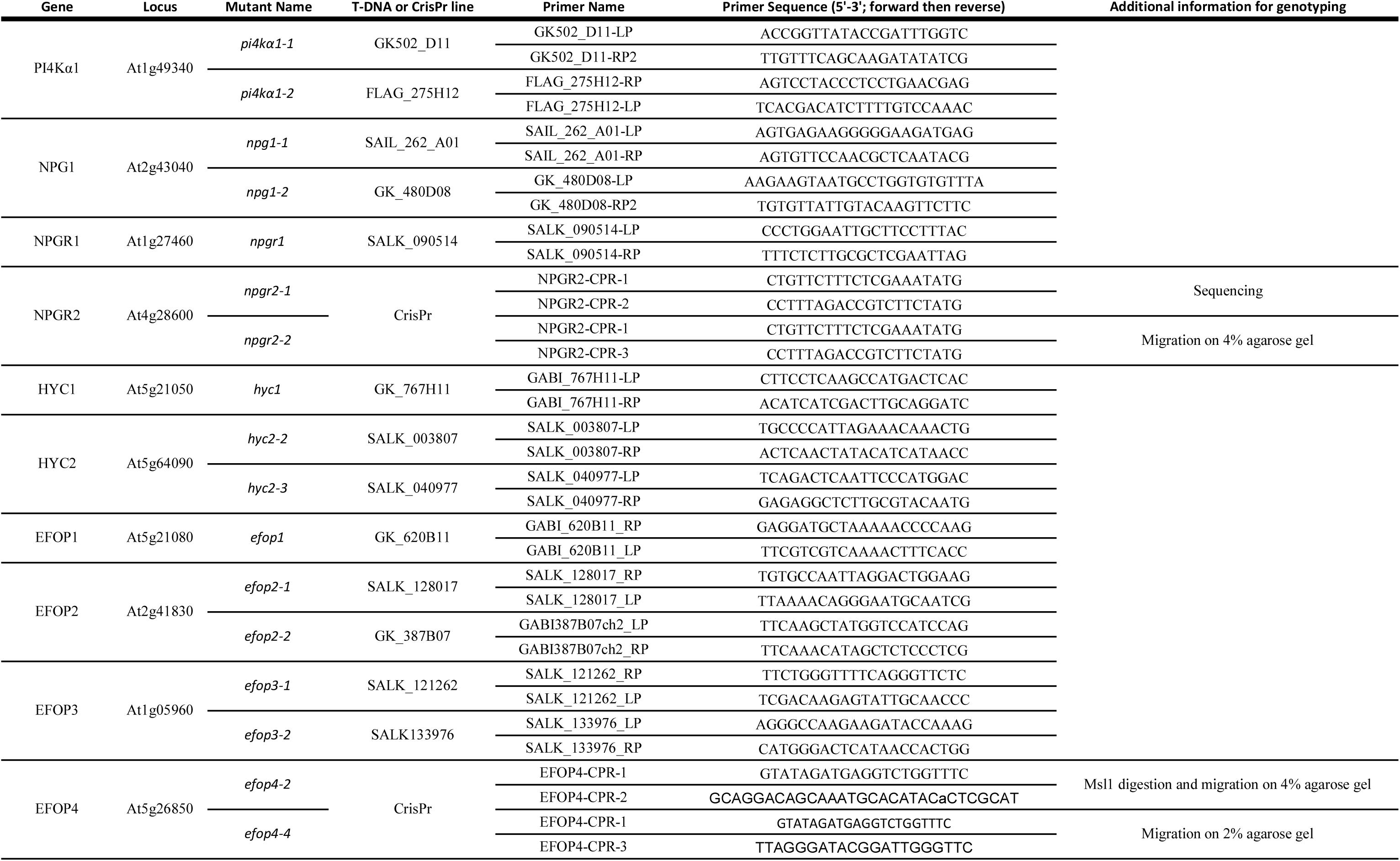
Genotyping primers.

**Table S9:**
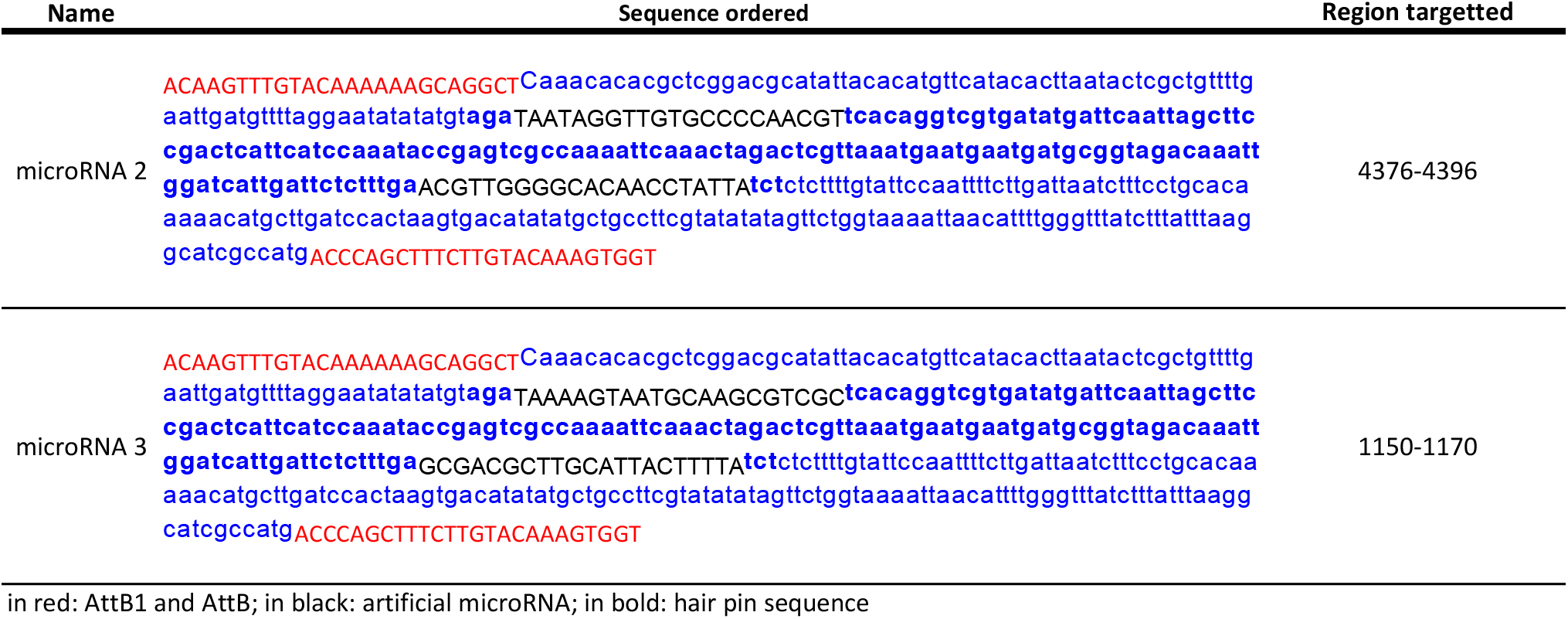
microRNA sequences.

**Table S10:**
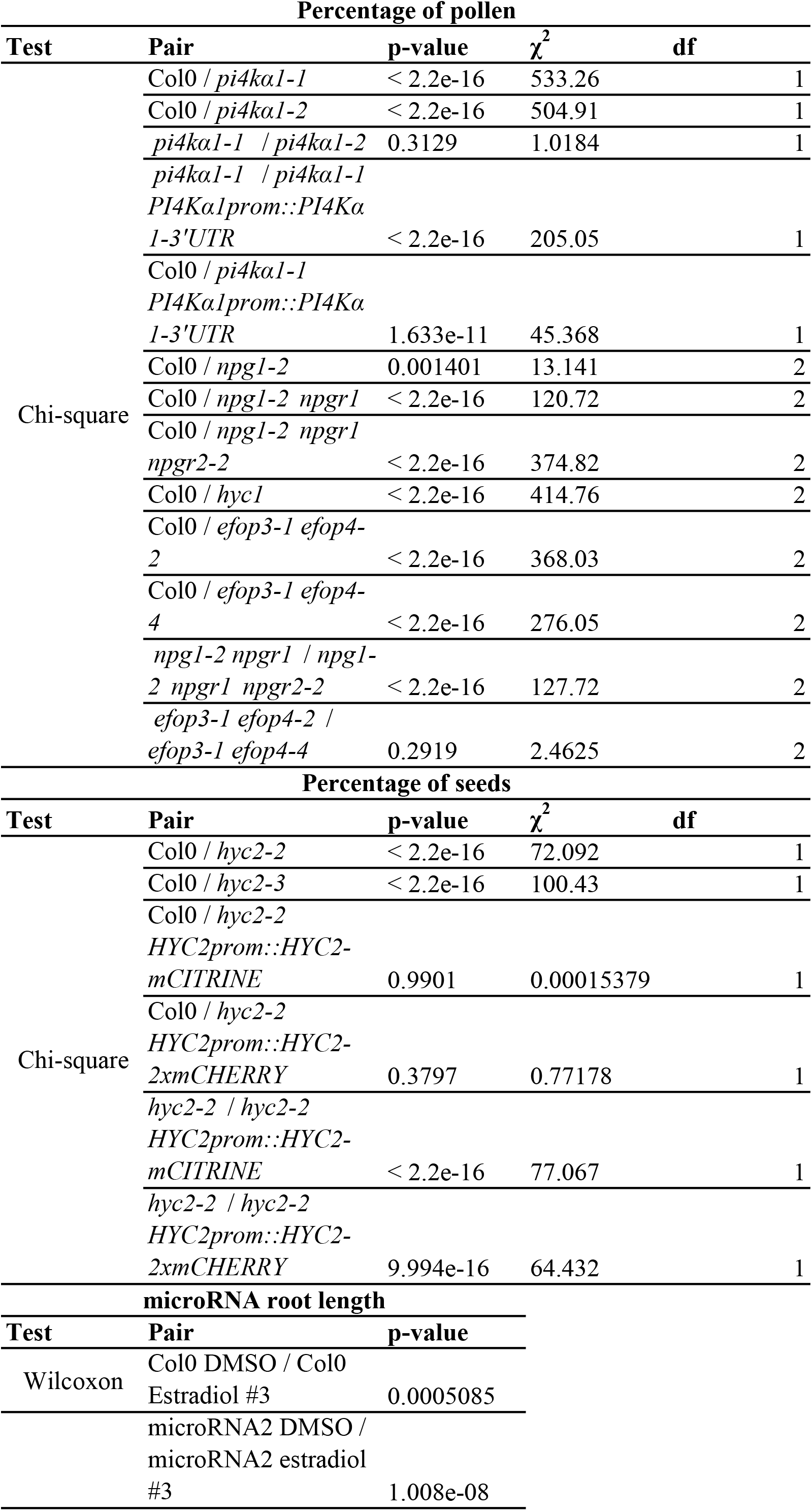
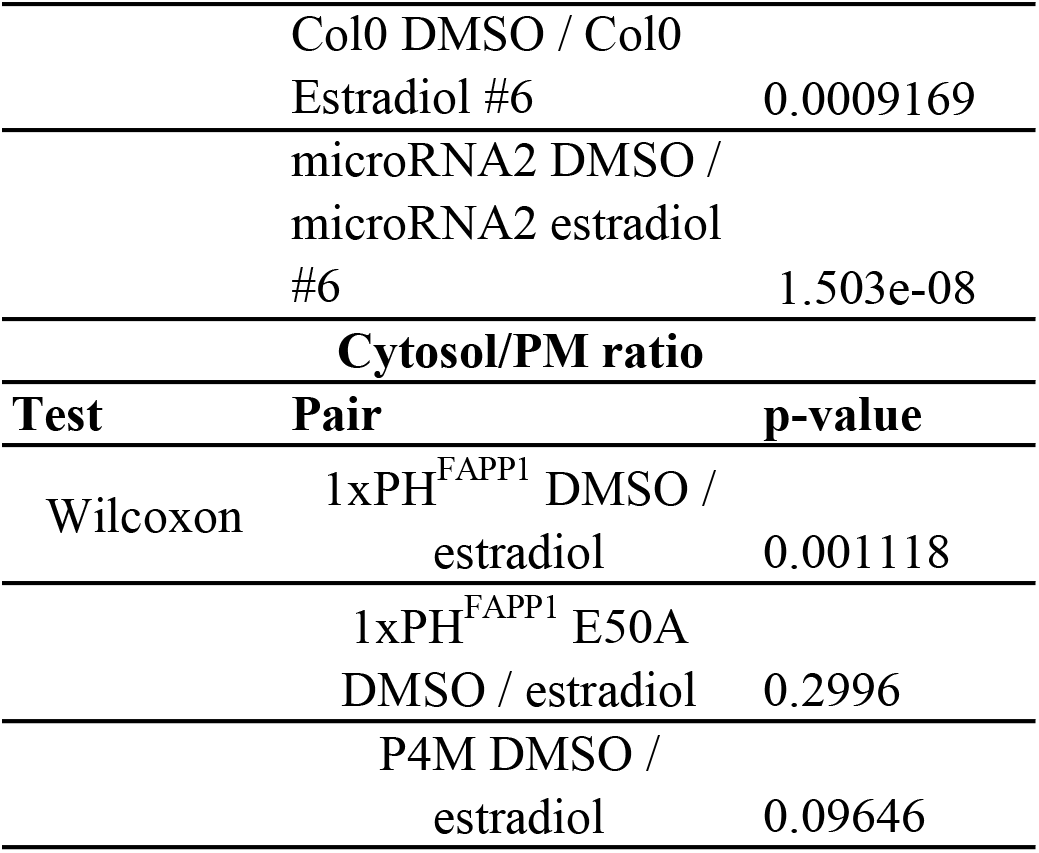
Statistical analysis.

## Notes

### Competing Interest Statement

The authors have declared no competing interest.

### Summary of Updates

New Figure 2, Figure 4 revised, Figure 5 revised, new Figure 6, Figure 9 revised, new Figure 10

